# Pseudo-chromosome length genome assembly of a double haploid ‘Bartlett’ pear (*Pyrus communis* L.)

**DOI:** 10.1101/651778

**Authors:** Gareth Linsmith, Stephane Rombauts, Sara Montanari, Cecilia H. Deng, Jean-Marc Celton, Philippe Guérif, Chang Liu, Rolf Lohaus, Jason D. Zurn, Alessandro Cestaro, Nahla V. Bassil, Linda V. Bakker, Elio Schijlen, Susan E. Gardiner, Yves Lespinasse, Charles-Eric Durel, Riccardo Velasco, David B. Neale, David Chagné, Yves Van de Peer, Michela Troggio, Luca Bianco

**Affiliations:** Center for Plant Systems Biology, VIB, Ghent, Belgium; Department of Plant Biotechnology and Bioinformatics, Ghent University, Ghent, Belgium; Fondazione Edmund Mach, San Michele all’Adige (TN), Italy; University of California Davis, Department of Plant Sciences, Davis, CA USA; The New Zealand Institute for Plant & Food Research Limited (PFR), Mt Albert Research Centre, Auckland, New Zealand; IRHS, INRA, Agrocampus-Ouest, Université d’Angers, SFR 4207 Quasav, 42 rue Georges Morel, F-49071 Beaucouzé, France; ZMBP, Allgemeine Genetik, Universität Tübingen, Auf der Morgenstelle 32, D-72076 Tübingen, Germany; USDA-ARS National Clonal Germplasm Repository, 33447 Peoria Road, Corvallis, OR 97333; Wageningen UR – Bioscience P.O. Box 16, 6700AA, Wageningen, The Netherlands; The New Zealand Institute for Plant & Food Research Limited (PFR), Palmerston North Research Centre, Palmerston North, New Zealand; CREA Research Centre for Viticulture and Enology, Via XXVIII Aprile 26, 31015 Conegliano (TV), Italy; Center for Microbial Ecology and Genomics Department of Biochemistry, Genetics and Microbiology, University of Pretoria, Pretoria 0028, South Africa

## Abstract

We report an improved assembly and scaffolding of the European pear (*Pyrus communis* L.) genome (referred to as BartlettDHv2.0), obtained using a combination of Pacific Biosciences RSII Long read sequencing (PacBio), Bionano optical mapping, chromatin interaction capture (Hi-C), and genetic mapping. A total of 496.9 million bases (Mb) corresponding to 97% of the estimated genome size were assembled into 494 scaffolds. Hi-C data and a high-density genetic map allowed us to anchor and orient 87% of the sequence on the 17 chromosomes of the pear genome. About 50% (247 Mb) of the genome consists of repetitive sequences. Comparison with previous assemblies of *Pyrus communis.* and *Pyrus x bretschneideri* confirmed the presence of 37,445 protein-coding genes, which is 13% fewer than previously predicted.

## Introduction

The genomics era has revolutionized research on fruit tree species and many of these genomes have recently been sequenced, or are currently being sequenced^1-2^. Nevertheless, although the cost for sequencing genomes has dropped considerably, obtaining high quality assemblies and annotations for complex plant genomes is still challenging ^3^. In addition to high numbers of repeats and transposable elements, high levels of heterozygosity complicate genome assembly for most fruit trees. Indeed, outcrossing fruit tree species often exhibit extremely high levels of heterozygosity with, for instance in apple ^4^, one single nucleotide polymorphism (SNP) every 50 base pairs (bp). The traditional solution to circumvent the challenge of heterozygosity is to sequence highly inbred plant material ^5-6^. However, such material may not always be available and many sequencing projects have used heterozygous samples, for sequencing of economically important cultivars ^7,8^.

Earlier assemblies of Asian pear (*Pyrus × bretschneideri*) ^8^, European pear (*Pyrus communis*) ^9^, and apple (*Malus × domestica*) ^7^ were based on heterozygous plant material, resulting in each case in erroneous and fragmented assemblies consisting of thousands of scaffolds. Both the Asian pear and apple genomes were subsequently re-assembled using different strategies to address the problem of extreme heterozygosity ^2,8^. In the case of Asian pear the genome was re-assembled using a BAC by BAC strategy, combined with Illumina sequencing ^8^. For apple, a double-haploid (DH) plant derived from the same cultivar, ‘Golden Delicious’, as the original reference genome was sequenced ^2^.

Here, we describe the assembly of the genome of the European pear (*Pyrus communis*) using a DH derived from the variety ‘Bartlett’, analogous to the strategy employed by Daccord et al. ^2^ in apple. The ‘Bartlett.DH’ developed at INRA, Angers, France ^10^ was chosen as it is derived from the same cultivar as employed for the previous European pear assembly, Bartlettv1.0, obtained by Roche 454 sequencing of extremely heterozygous plant material ^9^. This new genome sequence (BartlettDHv2.0) was assembled by combining short read Illumina and long read PacBio sequencing, optical mapping, Hi-C, and genetic maps. The BartlettDHv2.0 genome assembly improves the European pear assembly to 17 pseudo chromosomes and will be a critical tool for contemporary genomic studies in pear, including genome-wide association studies (GWAS) and genomic selection (GS) for the benefit of pear breeding.

## Results and Discussion

### Genome sequencing and assembly

A total of 31.4 Gb of PacBio RSII long read data was produced, comprising 3,665,270 reads with a read N50 of 14.2 kb. Reads longer than 10kb sum to 21.9 Gb. The RSII sequencing was supplemented by 123-fold coverage of Illumina (2×125bp) paired-end (PE) reads with a target insert size of 350 bp (61.5 Gb of sequence). Sequencing of two Hi-C libraries yielded 51.6 Gb of Illumina PE data as (2×125bp) reads. Kmer analysis of paired end Illumina data confirmed the homozygous nature of the ‘Bartlett.DH’ sample, with no heterozygosity peak visible in the 17-mer frequency distribution (Figure 1b vs. Figure 1a for Asian pear). Estimation of genome size from the 17-mer distribution provided an estimate of 528 Mb which agrees well with the 527 Mb genome size estimation made by Wu et al. ^8^ for Asian pear. The PacBio data therefore equates to 63-fold, long read coverage of the genome with 44 fold coverage in reads over 10kb.

**Fig1a.**
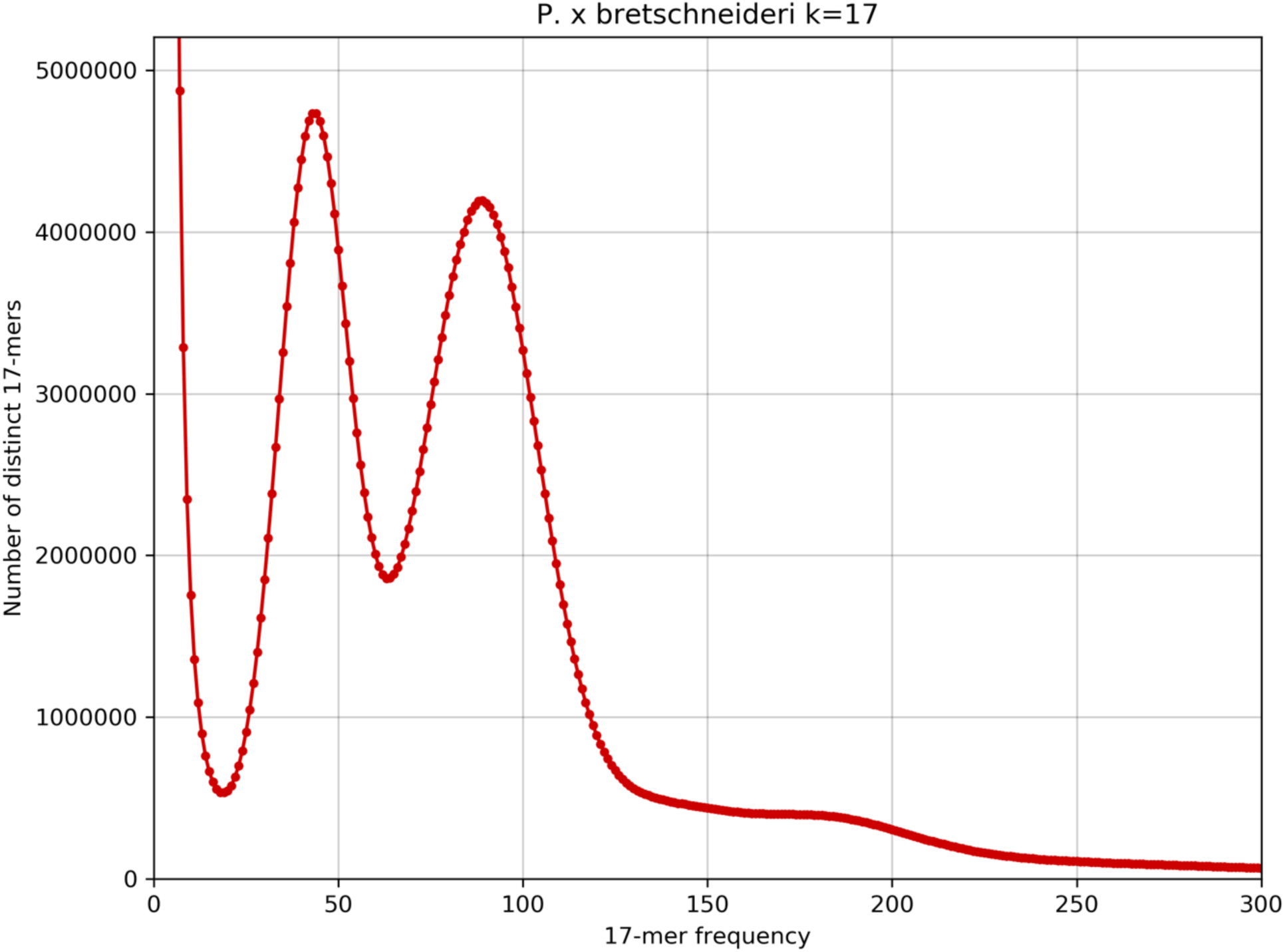
17mer frequency distribution of diploid *P* × *bretschneideri.* Using KAT^23^ v2.3.4, 17-mers were counted in all whole genome shotgun paired-end reads. The density plot of the number of unique kmer species (y axis) for each kmer frequency (x axis) is plotted. The homozygous peak is observed at a multiplicity (kmer coverage) of 86X, while the heterozygous peak is observed at 43X.

**Figure 1b.**
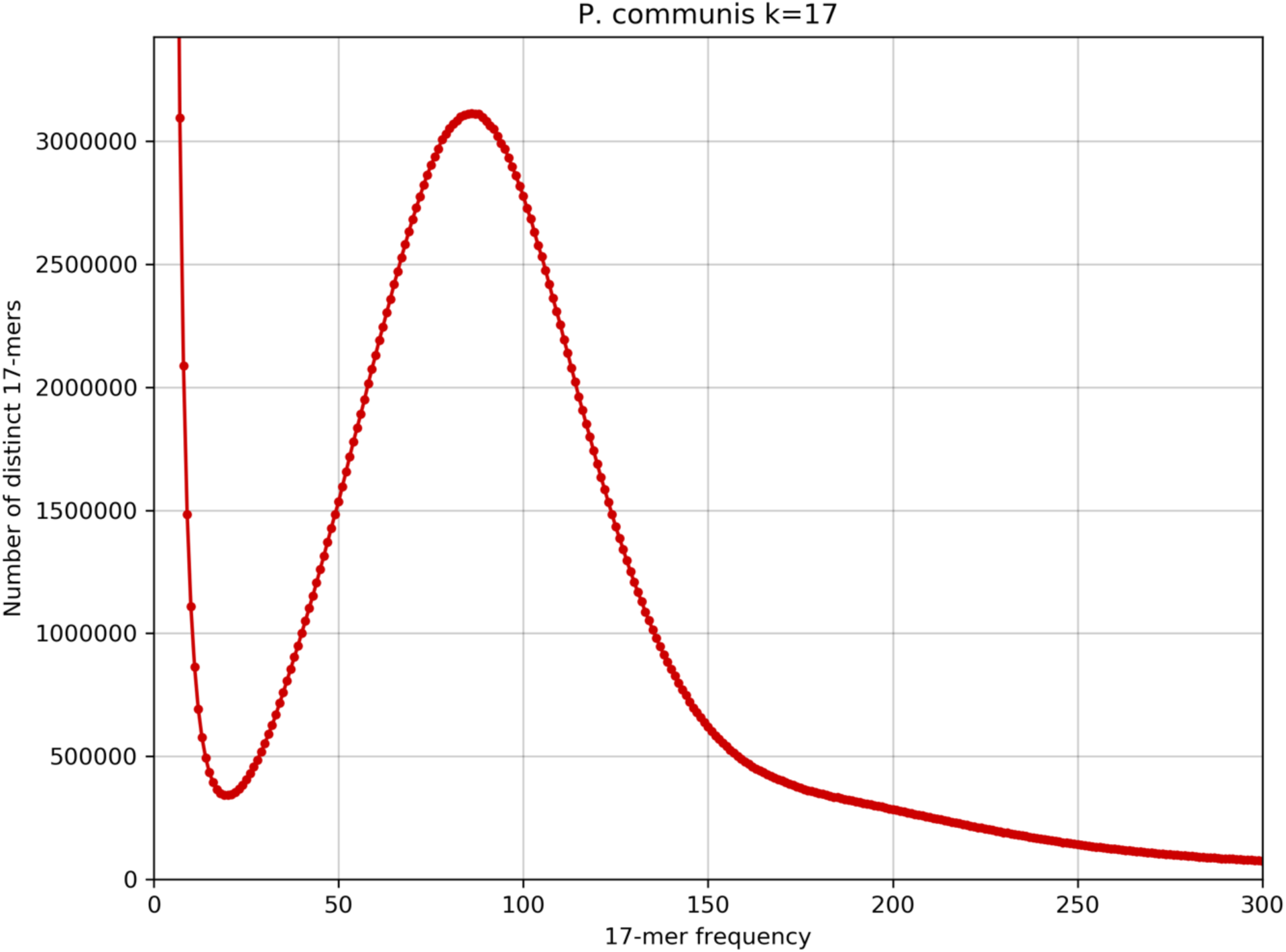
17mer frequency distribution of di-haploid *P. communis (BartlettDHv2.0).* Using KAT^23^ v2.3.4, 17-mers were counted in all whole genome shotgun paired-end reads. The density plot of the number of unique kmer species (y axis) for each kmer frequency (x axis) is plotted. The homozygous peak is observed at a multiplicity (kmer coverage) of 86X, while no heterozygous peak is observed.

The genome was assembled into 592 scaffolds totalling 496.9 Mb, or 94.0% of the expected genome size. The scaffold N50 is 6.5 Mb, which is a near 1,000-fold improvement over the Bartlettv1.0 assembly. Of these assembled scaffolds, 230 scaffolds totalling 445.1 Mb could be anchored to the 17 chromosomes of the pear genome using a combination of Hi-C data and the high-density genetic map. Thus 84.2% of the genome is anchored into 17 pseudomolecules with a further 51.8 Mb (477 smaller sequences) collected in LG0. These metrics are summarized in Table 1. Searching of telomeric sequences (5’-TTTAGGG-3’) enriched in the terminal parts of the pseudo-chromosomes allowed the identification of 22 telomers. Out of the 17 pseudo-chromosomes, all but Chr14 and Chr16 had at least one of the two telomers, and the presence of telomeric sequences could be found at both ends of seven of the pseudo-chromosomes (Chr4, Chr6, Chr8, Chr10, Chr11, Chr15 and Chr17). Additionally, SuperScaffold_290 was found to start with the 3’-CCCTAAA-5’ sequence, suggesting it may be located at the terminal the start of one chromosome. This hypothesis is also supported by three SNPs from this scaffold genetically mapping at the beginning of Chr02.

**Table 1.** Genome assembly metrics.

|  | Total assembled | % Genome | N50 | Number of Sequences |
| --- | --- | --- | --- | --- |
| Contigs | 501 Mb | 94.8% | 5.3 Mb | 620 |
| Scaffolds | 496.9 Mb | 94.0% | 6.5 Mb | 592 |
| Anchored into chromosomes | 445.1 Mb | 84.2% | 26.2 Mb | 17 |
| LG0 | 51.8 Mb | 9.8% | 0.19 Mb | 477 |

BUSCO analysis revealed 1,357 complete BUSCOs (94.3%) with 1.9% fragmented and 3.8% missing BUSCOs. Marey maps^11^, showing the relationship between genetic and physical distance across each chromosome, demonstrate good agreement between the assembly and the Bartlett genetic map (Supplementary figures S2-S18); an example showing Chromosome 1 is provided in Figure 2.

**Figure 2:**
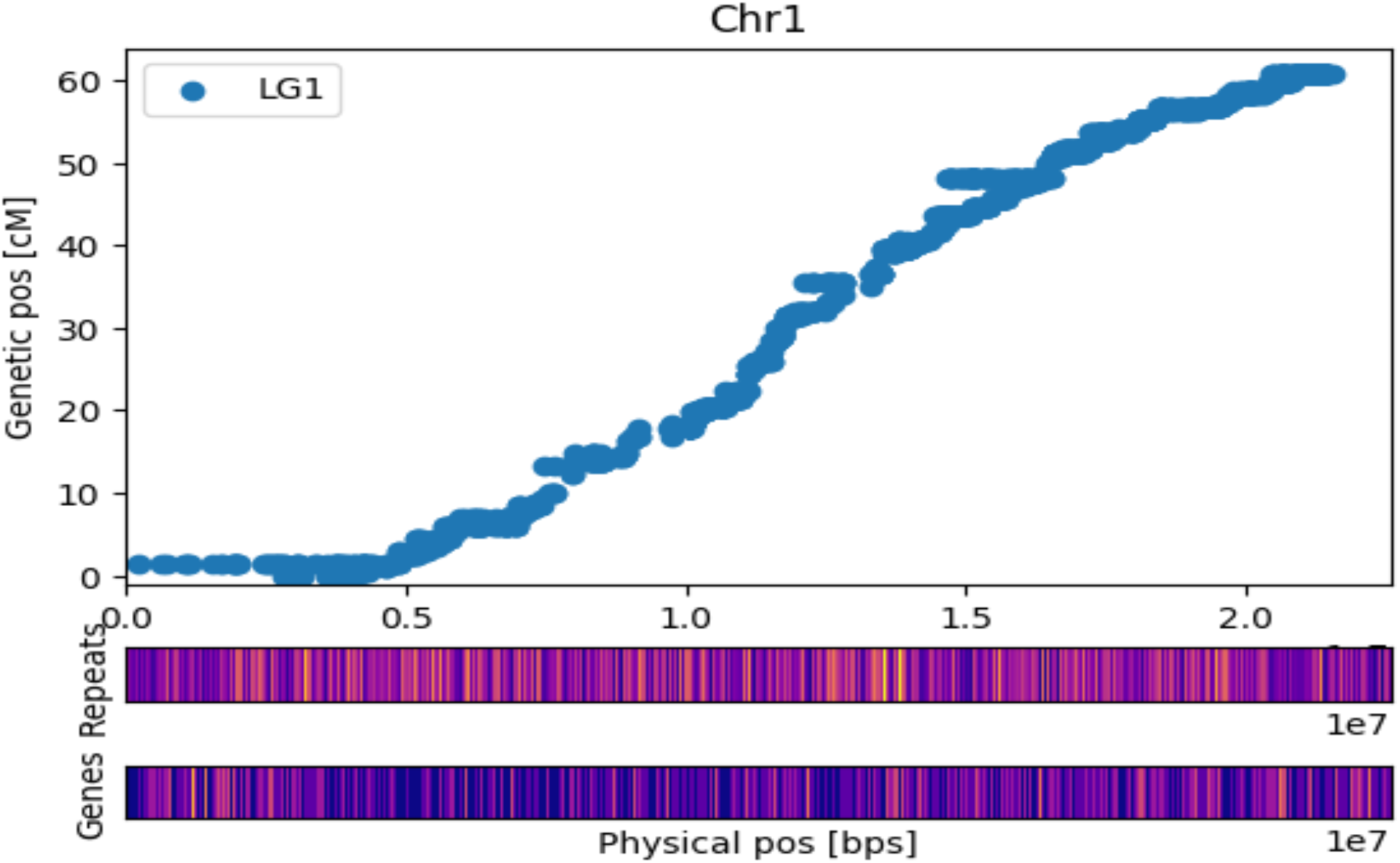
Marey plot of Chr1 with heatmap of Dispersed Repeats and Genes in bins of 200kb. The lighter the colour the more elements are present. Genetic positions refer to the high-density map of Bartlett. Dots represent the genetic and physic position (on BartlettDHv2.0) of 11,474 SNPs.

Using the high-density genetic map of Bartlett, a haplotype map for BartlettDHv2 was produced, allowing the identification of the two haplotypes of Bartlett in the BartlettDHv2 assembly, as well as the recombination breakpoints (Supplementary figure S1), confirming the gynogenesis origin of the BartlettDHv2 hypothesized by Bouvier et al^10^.

Summary statistics of two assemblies produced using Canu ^12^ and Falcon ^13^ are shown in Table 2. The Canu assembly has higher contiguity (501 Mb in 620 scaffolds), while the Falcon assembly produces a slightly larger, but more fragmented result (515 Mb in 1,282 scaffolds). Both assemblies were used for the optical mapping data analysis and results for both the Canu and Falcon assemblies are shown in Table 3. While the total amount of sequence is similar in both cases, the Canu assembly produced fewer conflicts with the optical mapping data than Falcon (13 vs. 38), as well as much longer scaffolds (scaffold N50 of 8.1 Mb vs 3.5 Mb in Canu and Falcon, respectively). Alignment with the high-density linkage map indicated that the Canu assembly produced fewer conflicts with the genetic map than the Falcon assembly (3 vs. 8). The Canu assembly was therefore selected as the contig assembly.

**Table 2:**
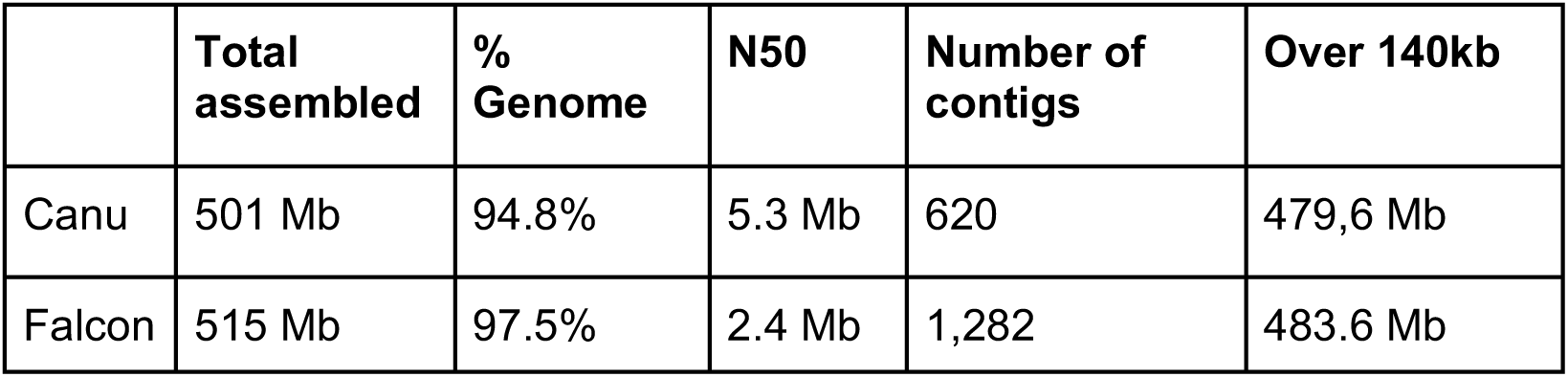
Summary statistics of best Canu and best Falcon contig assemblies.

|  | Total assembled | % Genome | N50 | Number of contigs | Over 140kb |
| --- | --- | --- | --- | --- | --- |
| Canu | 501 Mb | 94.8% | 5.3 Mb | 620 | 479,6 Mb |
| Falcon | 515 Mb | 97.5% | 2.4 Mb | 1,282 | 483.6 Mb |

**Table 3:** Summary statistics of the Canu and Falcon hybrid assemblies combined with the Bionano optical mapping data.

|  | **Bionano incorporated** | **% Genome** | **N50** | **Num scaffolds** | **Number of conflicts with optical map** |
| --- | --- | --- | --- | --- | --- |
| Canu + Bionano | 459.2 Mb | 87.0% | 8.1 Mb | 123 | 13 |
| Falcon + Bionano | 451.4 Mb | 85.4% | 3.5 Mb | 214 | 38 |

Consensus was called on the assembly using PacBio WGS, Illumina WGS and Illumina RNA-Seq data. A single iteration of consensus calling using raw PacBio data was followed by polishing with Illumina WGS data. This Illumina consensus calling was performed iteratively while monitoring the number of kmers shared between the assembly and the Illumina read data. This metric reached a maximum value after seven iterations and Illumina WGS consensus calling was halted at this point. Finally, iterative consensus calling was run using RNA sequencing (RNA-Seq) data instead of the WGS Illumina data in order to focus the consensus on coding sequence. The rationale for this was that small errors are particularly a problem in coding regions because they can introduce frameshifts that severely affect the annotation of genes. Metrics indicated that the consensus calling of coding regions was optimal after the second iteration. The second iteration of RNA-Seq consensus calling was therefore selected as the final scaffold assembly.

Combining scaffolds with proximity information from Hi-C sequencing enabled arrangement of the scaffolds into 17 ordered and oriented clusters representing the 17 chromosomes of the pear genome. Agreement of Hi-C clusters with the genetic map was not perfect but was very high, with 11 of the 17 Hi-C clusters being in perfect agreement with the genetic map. For such clusters, every anchored scaffold in that cluster is anchored to the same LG by the genetic map and no scaffold from another cluster was ever anchored to that LG. Comparison of the other 6 Hi-C clusters with the genetic map suggested that the Hi-C had correctly grouped and oriented chromosome arms. These clusters could be made to agree perfectly with the genetic map by splitting each of them into two. These remaining six clusters were therefore split and then re-joined in accordance with the genetic map.

### Comparison of BartlettDHv2.0 assembly with Bartlettv1.0 assembly

The Bartlettv1.0 assembly totals 507.7 Mb (excluding N’s), of which 99.8% (506.8 Mb of sequence in 141,034 out of the 142,083 original scaffolds) was aligned to the BartlettDHv2.0 assembly. Inter-assembly synteny is very strong, suggesting that although highly fragmented, the Bartlettv1.0 assembly was a veridical depiction of the genome. There is evidence of some haplotype separation in the Bartlettv1.0 assembly as 25,120 scaffolds totalling 25.6 Mb align to overlapping positions on the BartlettDHv2.0 assembly. Conversely, 1,974 scaffolds totalling 1.6 Mb, aligned to multiple places in the BartlettDHv2.0 assembly. These scaffolds represent repeats which are collapsed in the Bartlettv1.0 assembly, but not in the BartlettDHv2.0 assembly. This 1.6 Mb of repeat scaffolds from the Bartlettv1.0 assembly becomes 4.4 Mb of sequence in the BartlettDHv2.0 assembly, highlighting the importance of third generation, long read data in resolving the repetitive structures of plant genomes.

### Gene annotation and transcriptome sequence analysis

The combination of *ab initio* gene prediction with protein alignment and cDNA alignment prediction enabled the annotation of 37,445 protein-coding genes in the BartlettDHv2.0 assembly. In total 95% of these are supported by RNA-seq evidence. On average, gene models consisted of transcript lengths of 2,944 bp, coding lengths of 1,186 bp, and means of 10 exons per gene. These values are similar to those observed in Asian pear ^8^, apple ^2^, and the Bartlettv1.0 assembly ^9^ (Table 4). All gene models had matches in at least one of the public protein databases (nrprot or interpro), while 95% of them contained domains recognised in the interpro database. The average gene density in BartlettDHv2.0 assembly is 7.1 genes per 100 kb, with genes being more abundant in sub-telomeric regions, as previously observed in other sequenced plant genomes (Figure 2, Supplementary figures S2-S18).

**Table 4:**
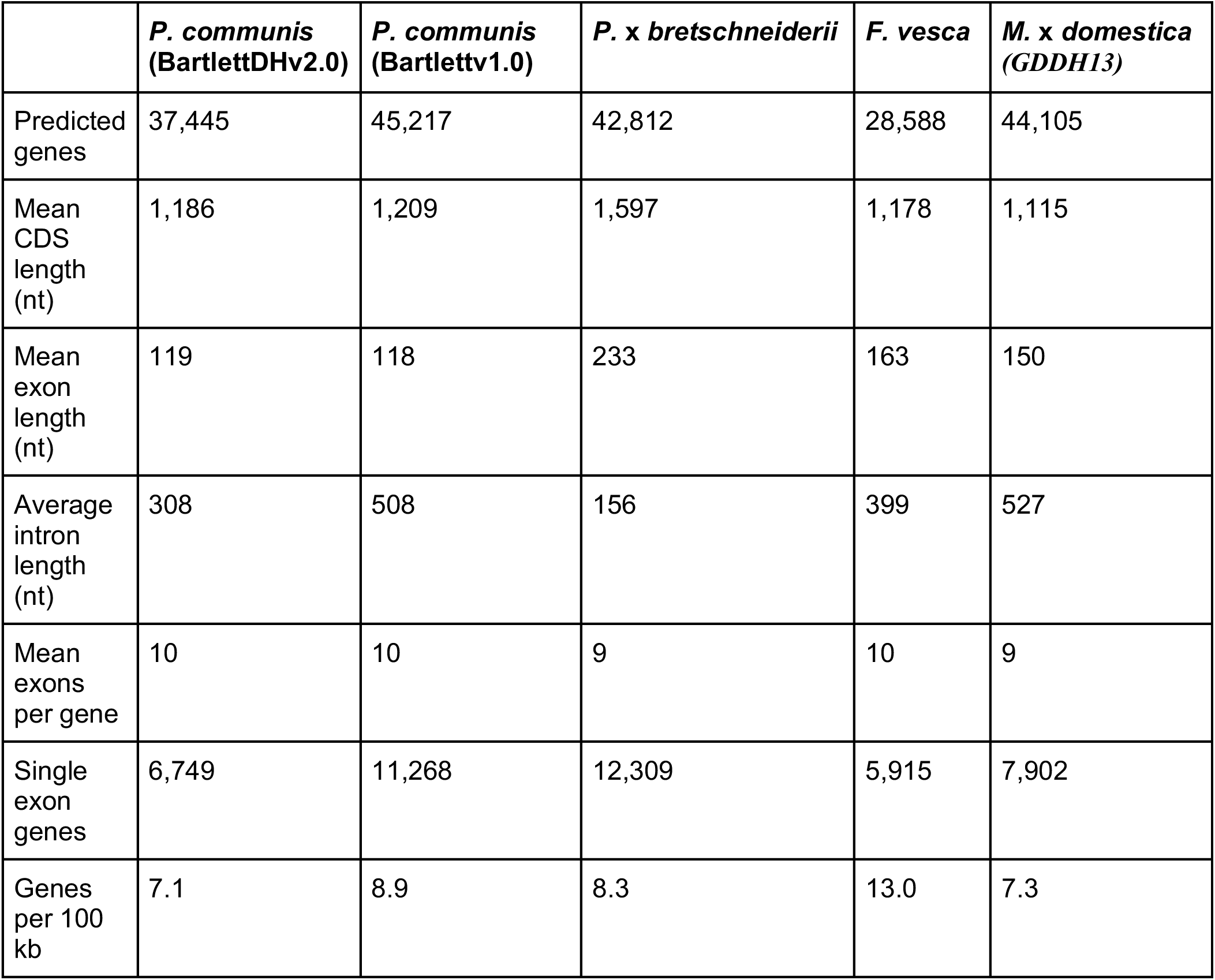
Summary statistics of gene annotation from selected Rosaceae species.

|  | *P. communis*<br>(BartlettDHv2.0) | *P. communis*<br>(BartlettV1.0) | *P. x bretschneiderii* | *F. vesca* | *M. x domestica*<br>(GDDH13) |
| --- | --- | --- | --- | --- | --- |
| Predicted genes | 37,445 | 45,217 | 42,812 | 28,588 | 44,105 |
| Mean CDS length (nt) | 1,186 | 1,209 | 1,597 | 1,178 | 1,115 |
| Mean exon length (nt) | 119 | 118 | 233 | 163 | 150 |
| Average intron length (nt) | 308 | 508 | 156 | 399 | 527 |
| Mean exons per gene | 10 | 10 | 9 | 10 | 9 |
| Single exon genes | 6,749 | 11,268 | 12,309 | 5,915 | 7,902 |
| Genes per 100 kb | 7.1 | 8.9 | 8.3 | 13.0 | 7.3 |

### Orthology analysis

The predicted protein sequences from European pear were compared with those from eight other species, *Pyrus x bretschneideri* ^8^, *Malus x domestica* (GDDH13)^2^, *Fragaria vesca* ^14^, *Prunus persica* ^15^, *Rosa chinensis* ^16^, *Rubus occidentalis* ^1^, *Vitis vinifera* ^17^, and *Arabidopsis thaliana* ^18^. Proteins were clustered into 20,677 orthologous groups (≥ 2 members), of which 8,877 (43%) were common to all nine genomes (Figure 3). Full results of the orthology analysis are available from the pear project database on request. A set of 414 gene clusters were identified as being specific to the three pome fruits analysed (i.e. to apple and the two species of pear). A set of 611 gene clusters were identified as being shared by the two pear species but not by apple. A set of 8 gene clusters was found to be specific to the European pear, while 22 gene clusters were specific to the Asian pear and 7 gene clusters were found to be specific to apple.

**Figure 3.**
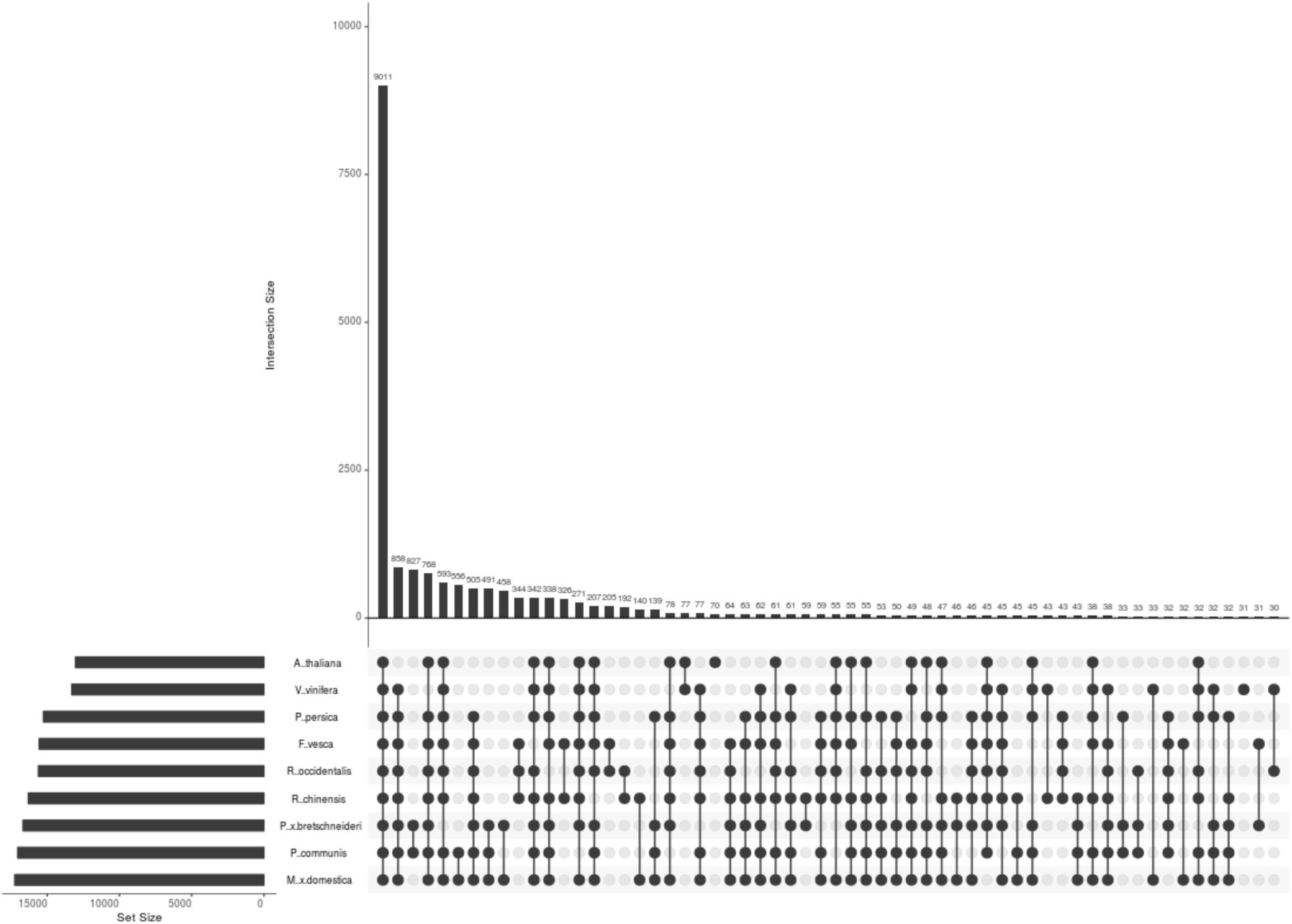
UpSet plot of protein clusters shared by nine species *P. x bretschneideri, P. communis, M. × domestica, F. vesca, P. persica, R. chinensis, R. occidentalis, V. vinifera*, and *A. thaliana*. UpSet plots describe the intersections of a set as a matrix were each column corresponds to a set, and each row corresponds to one segment in a Venn diagram. Cells are either empty indicating that this set is not part of that intersection, or filled showing that the set is participating in the intersection. The height of the bar represents the size of the corresponding intersection.

Gene clusters that were determined by the orthology analysis to be pear specific, or specific to one of the three Malinae species (Asian pear, European pear and apple) were queried in the other Malinae genomes by aligning gene sequences with Genome Threader ^19^. This gene sequence re-alignment revealed that, in most of these cases, gene clusters shown to be organism specific by the orthology analysis, revealed genes which were missed by the automatic annotation of the respective genome assemblies. All Asian pear and European pear specific gene clusters could be identified in one of the other Malinae genomes, while 5 gene clusters were found to be genuinely apple specific. Of the 611 pear specific gene clusters, 526 were found in the apple genome. Of the remaining 85 pear specific gene clusters 74 are supported by Rna-Seq in *Pyrus communis* 31 have a functional annotation and all 85 have either Rna-Seq support or a functional annotation. The gene structures resolved by alignment of Asian pear and apple genes were merged with the BartlettDHv2.0 annotation adding a further 209 gene models.

The results of this gene structure re-alignment highlight the limits of automated gene annotation and the importance of ongoing curation of gene structure annotations. An example of the importance of manual curation of gene models has recently been reported in kiwifruit, where more than 90% of the *in silico* predicted gene models were re-annotated compared to a previous draft version ^20^. The annotation of the BartlettDHv2.0 assembly has been loaded into the online resource for community annotation of eukaryotes (ORCAE) ^21^ to facilitate ongoing manual curation of gene models.

### Whole-genome duplication

Distributions of synonymous substitutions per synonymous site (*K*_S_) produced for the whole paranomes of *P. communis, P.* × *bretschneideri*, and *M.* × *domestica* all support the common whole-genome duplication (WGD) event shared by the Malinae. Signature WGD peaks in the *K*_S_ plots for the three species can be found at almost identical *K*_S_ values of ∼0.16 (Figure 4 a,b,c), as expected based on previous research ^2,7–9^. Comparison of these WGD *K*_S_ peaks with the *K*_S_ peaks of ortholog distributions between pears and apple and between pears/apple and rose (*Rosa chinensis*) ^16^ suggest that the WGD occurred quite a long time after the divergence of Amygdaloideae and Rosoideae and well before the divergence of pear and apple (unless substantial substitution rate acceleration/deceleration occurred in these lineages).

**Figure 4.**
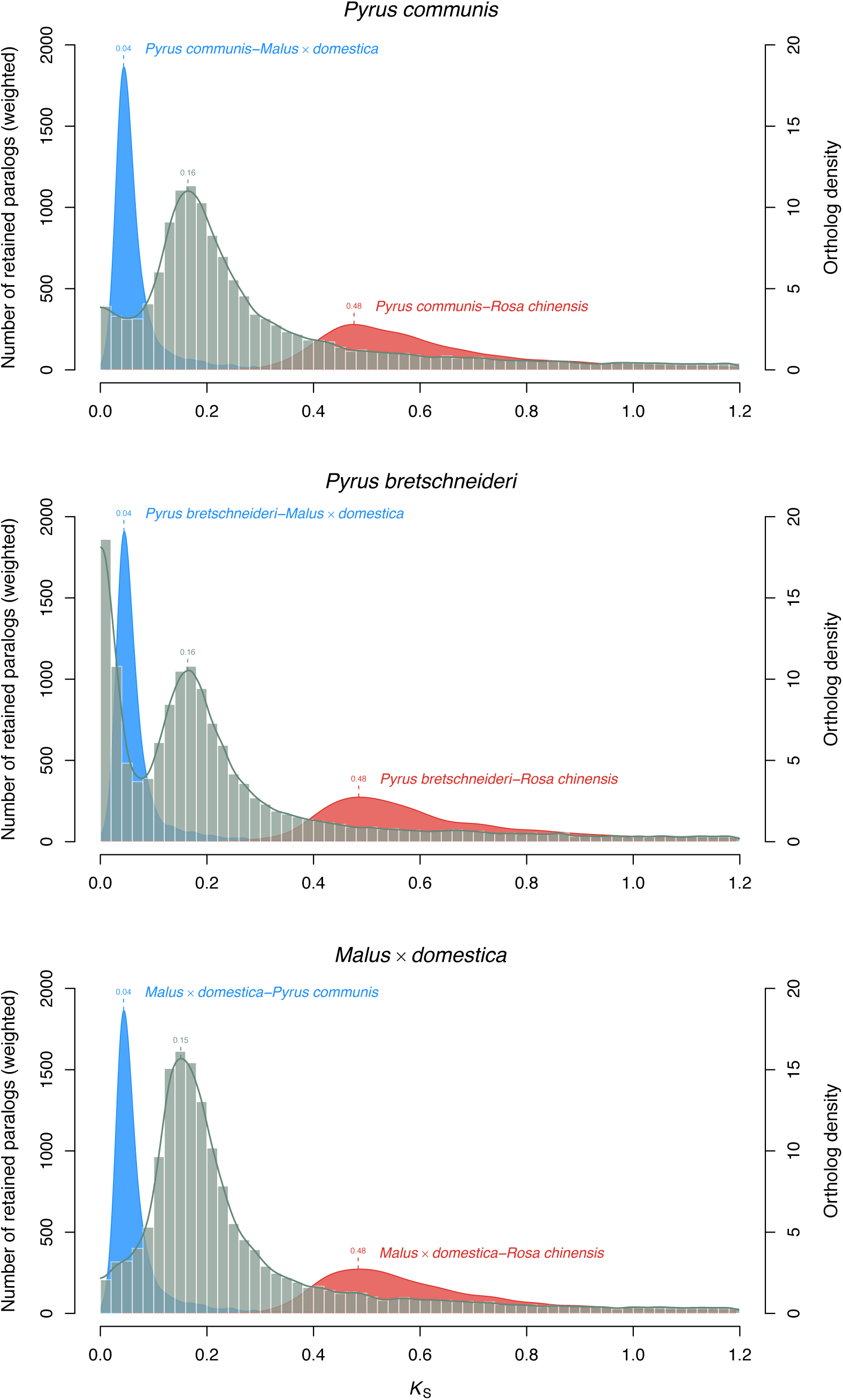
(a,b,c) Paralog K_s_ distributions of *P. communis BartlettDHv2.0, P.* × *bretschneideri* and *M.* × *domestica* GDDH13 (grey histograms and line, left-hand y-axes; a peak represents a WGD event) and one-to-one ortholog *K*_s_ distributions between indicated species (blue and red filled curves of kernel-density estimates, right-hand y-axes; a peak represents a species divergence event).

### Functional annotation and GO enrichment analysis

A combination of BLAST (NR prot) and interproscan searches enabled the annotation of 12,444 of the 37,445 genes (33%) with a functional description. Loading predicted transcripts into the TRAPID online annotation platform ^22^ enabled annotation of 24,257 (69%) genes with at least one GO term. GO enrichment analysis was performed within the TRAPID platform on gene sets of particular biological interest, i.e. pear specific gene families and pome specific gene families. No enriched GO terms were found for the pear specific gene families, while significantly enriched GO terms for the pome specific gene families are presented in the supplementary material.

### Repetitive element annotation

A combination of *de novo* and homology based repeat annotation identified a total of 247 Mb of transposable element sequences accounting for 49.7% of the assembly. As is typical for plant genomes, the most abundant transposable elements are retrotransposons of the long terminal repeat (LTR) family, totalling 32.6% of the genome. Although widely dispersed throughout the genome, transposon-related sequences were most abundant in centromeric regions.

The recent reassembly of the apple genome ^2^ revealed a previously undescribed LTR element dubbed ‘HODOR’ (or High Copy Golden Delicious Repeat) and the expansion of this element was implicated as having a potential role in the speciation of apple and pear. This element has now been verified in the pear genome. BLAST analysis revealed 232 full length HODOR copies in the BartlettDHv2.0 genome, only 29% of the number of full length copies identified in the apple genome. Although the HODOR element has, to date, only been identified in the apple and pear genomes, this finding must be treated with a degree of caution. The apple and pear genomes have been reassembled using the latest long read technology to arrive at chromosome scale assemblies. HODOR is a 9.2kb transposable element, and as such it may simply not have been completely assembled in previous Rosaceae genomes based on short read data. Nevertheless, BLAST searches reveal no trace of the HODOR element in the recent chromosome scale reassemblies of *Fragaria vesca* ^14^, *Rosa chinensis* ^16^, or *Rubus occidentalis* ^1^, all of which were developed from long read data.

Future in depth studies into the repeat content of Rosaceae genomes may reveal the point in the evolution of the Rosaceae, at which this element first emerged and how it relates to phenotypic differences among Rosaceae species.

### Chromosome structure

All 17 chromosomes of the European pear genome displayed strong nucleotide level synteny with the recent chromosome scale assembly of the apple genome ^2^ (Supplementary Figures S19b-S35b). Although only a scaffold level assembly of the Asian pear is publicly available at this time, 1,913 of the 2,182 scaffolds (82%) from the Asian pear assembly can be aligned to the European pear assembly. The aligned scaffolds sum to 495 Mb or 99.5% of the Asian pear assembly. Of the 1,913 aligned scaffolds, there are 882 scaffolds totalling 403.8 Mb (or 81% of the Asian pear assembly) which align unambiguously to the 17 assembled pseudomolecules. Numerous small-scale inversions with respect to European pear are evident within Asian pear scaffolds and any of these small-scale structural differences could prove to be of biological interest.

Self synteny of the genome based on colinear gene blocks reveals that the syntenic chromosome pairs for apple^7^ and pear ^10^ (LG3 and LG11, LG5 and LG10, LG9 and LG17, and LG13 and LG16) are clearly identifiable (Figure 6) and most collinear regions in strawberry correspond to two regions in European pear (Figure 7), as described for both apple and Asian pear ^9, 10^. Hence, the BartlettDHv2.0 assembly confirms that macrosyntenic chromosome structure is conserved across the Malinae.

**Figure 5.**
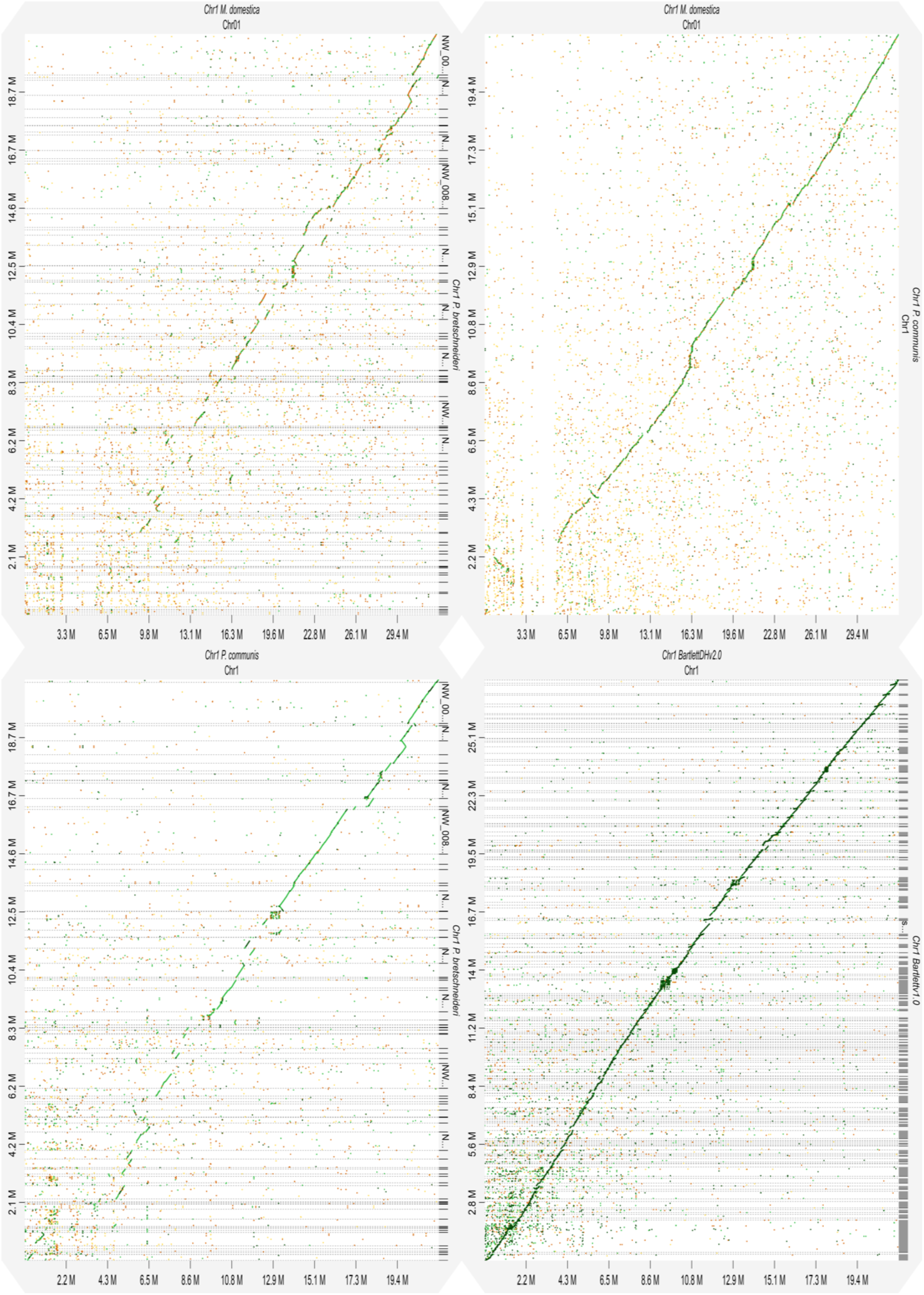
Chromosome 1 alignment dot plots. Dot plots are produced using the DGENIE software^60^ and alignments with minimap2 (v2.16). (Figure 5a) Dot plot of Chromosome 1 *P. × bretschneideri* to *P. communis - BartlettDHv2.0* (top left) (Figure 5b) Dot plot of Chromosome 1 *P. communis* BartlettDHv2.0 to *M. × domestica* - GDDH13 (top right) (Figure 5c) Dot plot of Chromosome 1 *P. × bretschneideri* to *M. × domestica – GDDH13* (bottom left) (Figure 5d) Dot plot of Chromosome 1 *P. communis* Bartlettv1.0 to *P. communis* BartlettDHv2.0 (bottom right)

**Figure 6.**
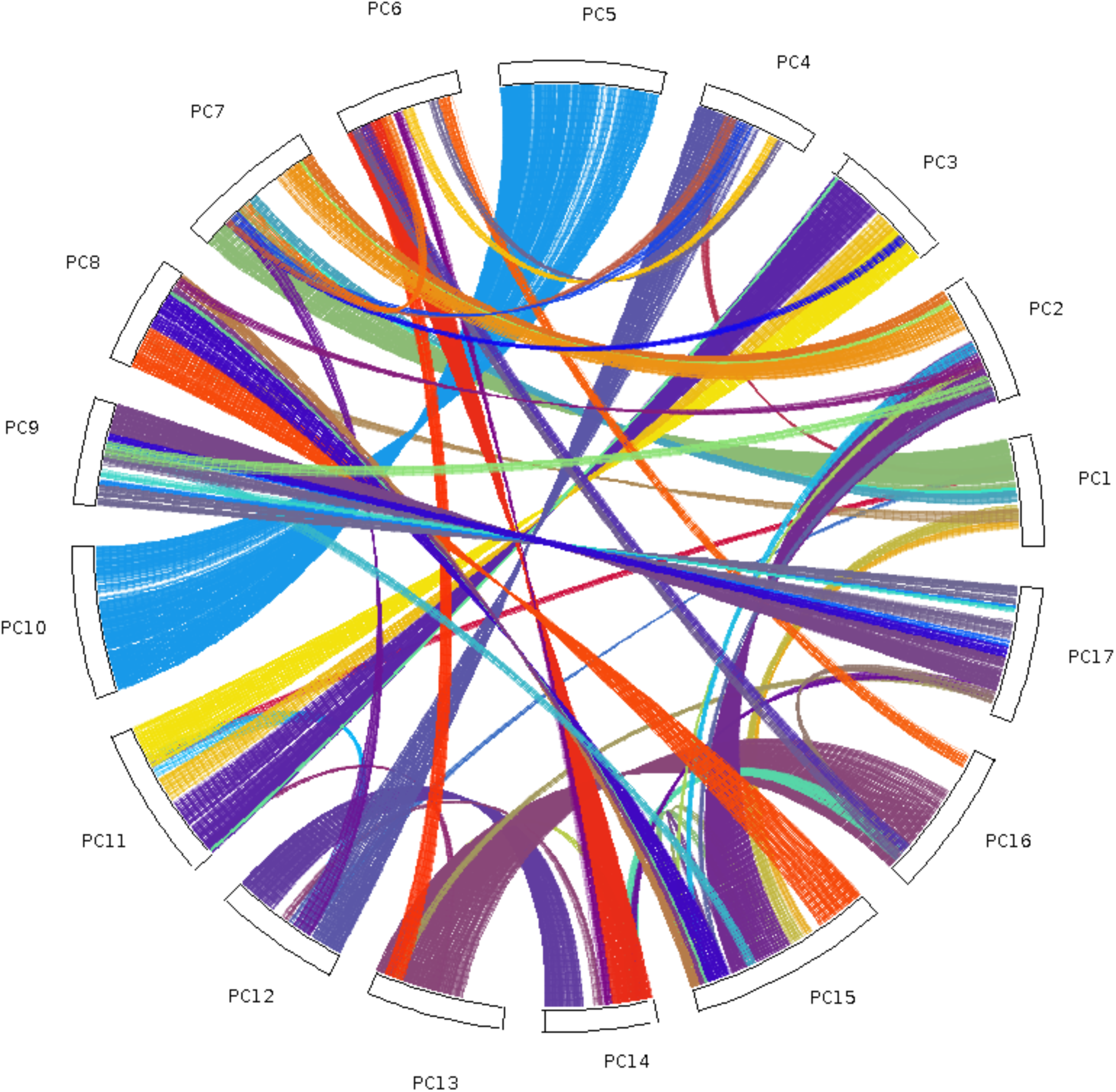
Self-collinearity of *P. communis* (BartlettDHv2). The coloured lines link collinearity blocks representing syntenic regions that were identified by MCScanX.

**Figure 7.**
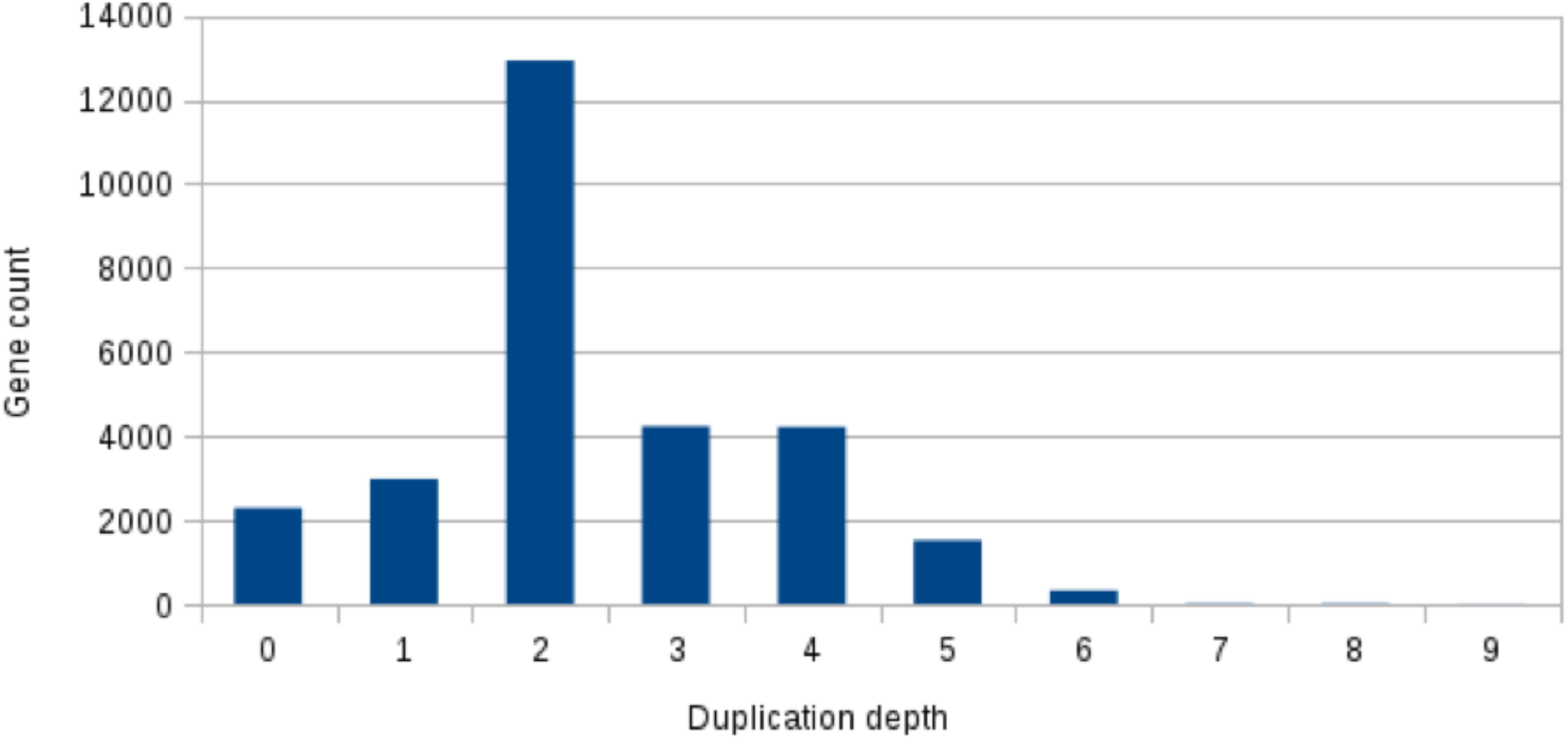
Duplication depth of *F. vesca* gene homologs in *P. communis* (BartlettDHv2). Inter species colinearity between *F. vesca* and *P. communis* was interrogated using MCScanX and at each gene locus of the *F. vesca* assembly the number of *P. communis* - *F. vesca* inter species colinear blocks (duplication depth) was counted. The number of *F. vesca* gene homologs having each copy number in the *P. communis* (BartlettDHv2) assembly is then plotted. It can be seen that most gene loci from *F. vesca* occur twice in *P. communis.*

### Revision in gene number in *Pyrus* species

Many Asian pear scaffolds align to overlapping positions on the BartlettDHv2.0 assembly. The same is also true of the Bartlettv1.0 assembly. These overlapping scaffolds most likely represent assembly of both haplotypes at the same genomic locus. Over-assembly is a danger when assembling a highly heterozygous genome and such separation of the haplotypes led to over-estimation of the gene number for apple ^2 7^ Re-examination of apple gene predictions and removal of overlapping gene models enabled Wu et. al. ^8^ to arrive at a new, lower estimate of the gene number for apple. Gene annotation of the BartlettDHv2.0 assembly resulted in a lower number of predicted genes than reported for the closely related Asian pear ^8^, or indeed the *P. communis.* Bartlettv1.0 assembly ^9^. When *P.* × *bretschneideri* gene models were aligned to the BartlettDHv2.0 assembly and overlapping gene models were collapsed down to a single locus, only 31,203 independent gene loci were identified, a reduction of 27% compared with the Asian pear assembly. Performing the same analysis with gene models from Bartlettv1.0 results in 37,997 independent loci. Thus, the removal of overlapping genes brings the number of gene predictions for the two *P. communis* assembly versions and the two sequenced *Pyrus* species much more closely in line (Table 5).

**Table 5.** Numbers of non-overlapping gene models in the three *Pyrus* gene annotations.

|  | **Total gene models** | **Non redundant gene models (no overlap on BartlettDHv2.0 assembly)** |
| --- | --- | --- |
| Bartlett v1.0 | 44,897 | 37,997 |
| Asian pear | 43,096 | 31,314 |
| BartlettDHv2.0 | 37,445 | 37,445 |

As further evidence of the over-assembly of the *P.* × *bretschneideri* genome the WGS read data from the *P.* × *bretschneideri* sequencing project was compared to the assembly in kmer space, using the kmer analysis toolkit^23^(Figure 8) and the same comparison was made for the BartlettDHv2.0 assembly (Figure 9). In these figures the kmer spectrum of the reads is colour coded to represent how many kmers of each frequency from the read’s spectrum ended up not included in the assembly, included once, included twice etc. Examination of Figure 8 reveals a significant amount of the content from the main peak of homozygous, single copy content is included twice, or even three times in the assembly (purple and green parts of the second peak). For the most part the heterozygous content has been properly collapsed in the *P.* × *bretschneideri* assembly (the black part of the first peak is covering almost 50% of the peak area) however, there is also heterozygous content which has been included twice in the assembly (purple part of first peak). Examination of. Figure 9 shows some slight duplication of content from the single copy peak (purple part at the top of the main peak) but this is much less severe than in the case of the *P.* × *bretschneideri* assembly. This finding is in accordance with the overlapping P. × bretschneideri scaffold alignments evident in the dot plots and accounts for the higher number of gene models reported for P. x bretschneideri

**Figure 8.**
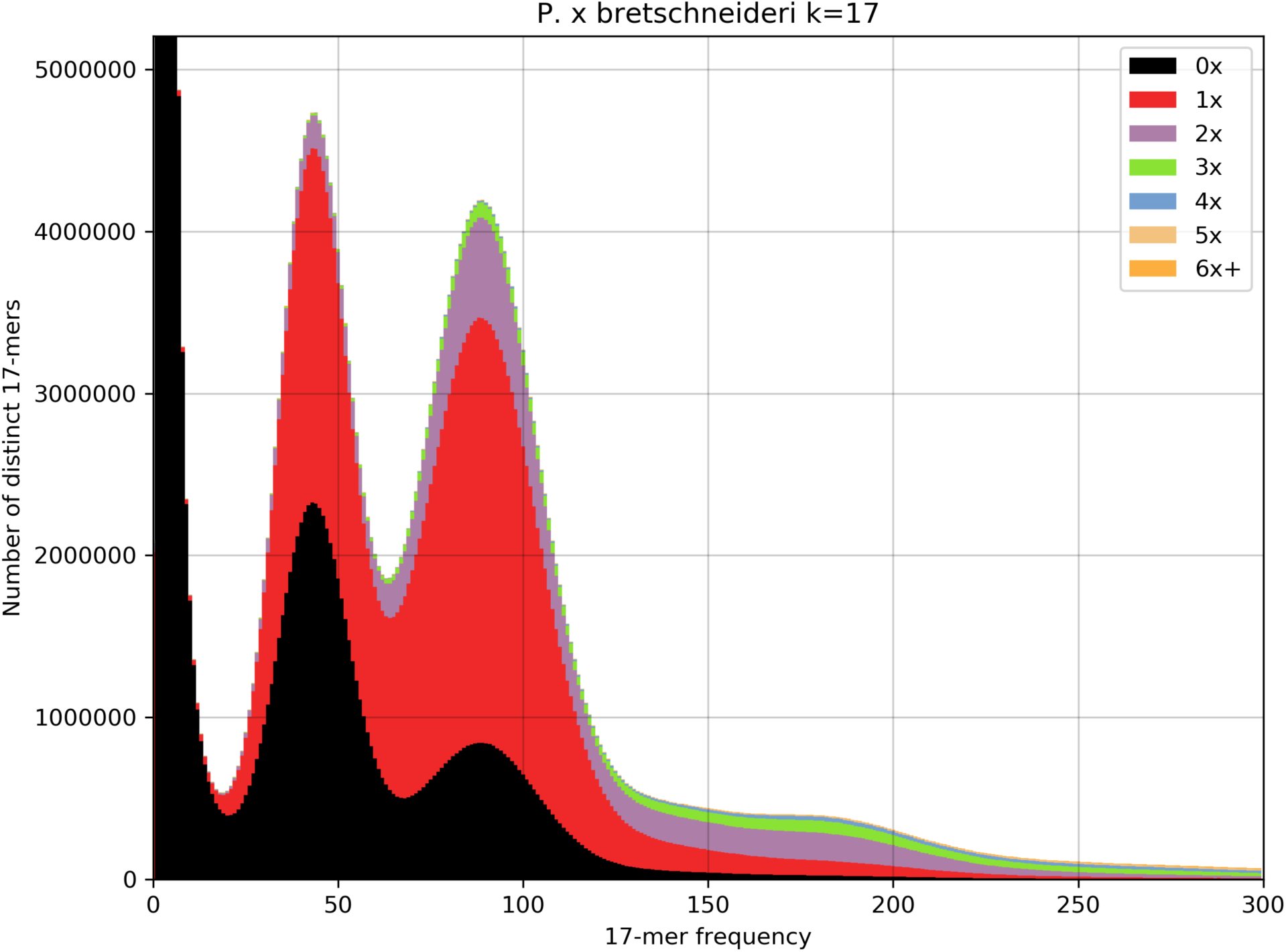
Kmer spectrum copy number plot for *P. × bretschneideri (k=17).* Using KAT^23^ v2.3.4, 17-mers were counted in all whole genome shotgun paired-end read and also in the *P. × bretschneideri* assembly. The kmer spectrum of the reads is then annotated with colours to denote the proportion of kmers with that frequency occurring once, twice, three times, etc. in the assembly. It is evident from this plot that a significant fraction of the kmer content which should appear once in the genome (content from the homozygous single copy peak at 86X) appears twice or even three times in the assembly (purple and green coloured parts of the single copy peak at 86X).

**Figure 9.**
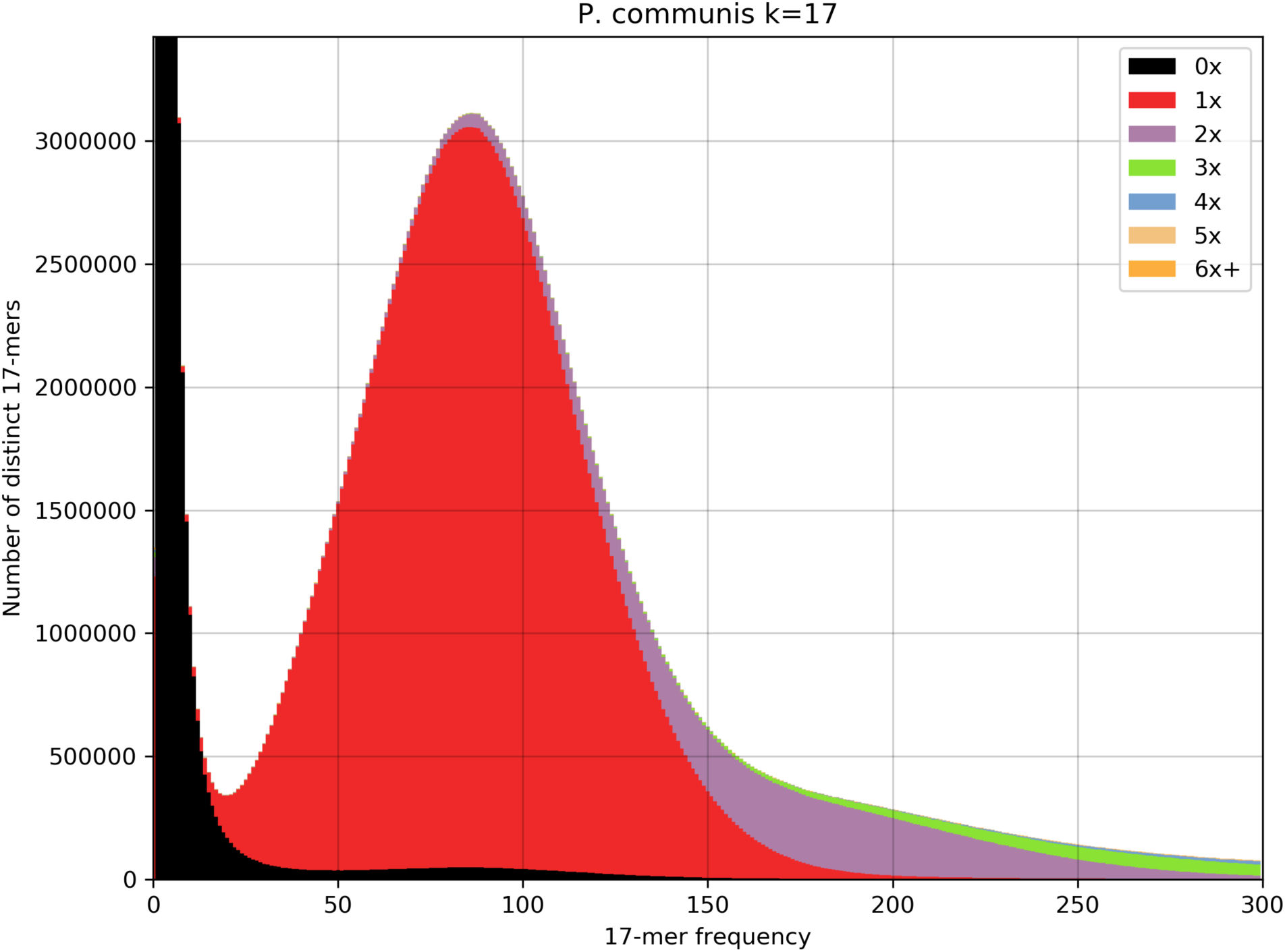
Kmer spectrum copy number plot for BartlettDHv2 *(k=17).* Using KAT^23^ v2.3.4, 17-mers were counted in all whole genome shotgun paired-end read and also in the BartlettDHv2 assembly. The kmer spectrum of the reads is then annotated with colours to denote the proportion of kmers with that frequency occurring once, twice, three times, etc. in the assembly. It is evident from this plot very little of the kmer content which should appear once in the genome (content from the homozygous single copy peak at 86X) is duplicated in the assembly (purple part of the single copy peak at 86X). The vast majority of the kmers that appear twice in the assembly (purple coloured parts of the figure) are in the small bump in the reads kmer spectrum centred at 172X i.e. they are kmers from the duplicated part of the genome.

### Conclusions

Cost effective, high throughput, long read sequencing is democratising the effective assembly of complex genomes, particularly the repeat rich genomes of plants. These advances in sequencing technology have enabled the improvement, or complete re-assembly of the draft genome sequences which have been typical of non-model organisms, including those of *Pyrus* species. This new improved assembly of the genome of *P. communis* will enable step changes in the progression of genome based technologies for pear breeders, analogous to those being developed for *Malus* following publication of the Golden Delicious v3.0 assembly ^24^. These include the ability to undertake genomic selection, and develop genetic markers based on candidate genes for traits of interest to breeders. These markers could be identified in the genome assembly following QTL mapping, or genome wide association studies. Such technologies will enable more efficient and targeted breeding of new varieties of pear with attributes that are desired by consumers and are also grower-friendly.

Values in column 2 for Bartlettv1.0 and Asian pear are from Chagné et al. ^9^ and Wu et al. ^8^ respectively.

## Materials and Methods

### Breeding the doubled haploid plant from ‘Bartlett’

In 1994, the European pear variety ‘Bartlett’ (synonymous ‘Williams’) was crossed as a female parent with the variety ‘Passe Crassane’ (male). Among the 971 seedlings obtained after sowing in the greenhouse in 1996, one showed the typical phenotype of pear haploid plants, i.e. a smaller size compared to diploid seedlings, with a slender stem and narrow, thin leaves of a pale green colour ^25^. This haploid plant (referenced W65) was confirmed by flow cytometry and propagated *in vitro* until development was sufficient for chromosome doubling experiment which was performed in 1998 with oryzalin based on a protocol adapted from apple^26^. The doubled haploid plant W65DH (here called ‘Bartlett.DH’) was confirmed as homozygous by isozyme and microsatellite markers ^25^ and also with the recently developed 70K SNP array ^33^ (data not shown). ‘Bartlett.DH’ was grafted on rootstock ‘Adams’ and is kept in an experimental orchard at INRA, Angers, France.

### Sample preparation and sequencing

For Illumina sequencing, genomic DNA from ‘Bartlett.DH’ was purified from young rolled leaves and young meristem tissue using the NucleoSpin Plant II DNA extraction kit (Macherey-Nagel GmbH, Düren, Germany), following the manufacturer’s instructions. One Illumina PE library was constructed at CNAG-CRG, Barcelona, Spain, with 340bp insert size according to KAPA Library Preparation Kit with no PCR Library Amplification/Illumina series (Roche-Kapa Biosystems) protocol and sequenced on HiSeq2000 (v4) in a single lane. For the BioNano and PacBio single molecule real time sequencing, genomic DNA was extracted using a modified nuclei preparation method ^27^ identical to that used in ^2^ followed by an additional phenol-chloroform purification step. Thirty SMRT cells were sequenced on the Pacific Biosciences RSII platform with the P5-C3 chemistry at the Genome Center at UC Davis.

### Hi-C library preparation and sequencing

The in situ Hi-C library preparation was performed according to a protocol established for rice seedlings with minor modifications ^28^. The libraries were made from two biological replicates of ‘Bartlett.DH’; for each replicate, 0.5 g of fixed leaves were used as the starting material. Due to the presence of large amount of cellular debris after isolation of nuclei, the nuclear pellet was divided into five parts prior to chromatin digestion with *Dpn*II. The Hi-C libraries were sent to the Australian Genome Research Facility (Melbourne, Australia) for sequencing using one lane of 100 bp PE sequencing using a HiSeq2000 (Illumina Inc.).

### BioNano Genomics genome mapping

Agarose plug embedded nuclei were Proteinase K treated for two days followed by RNAse treatment (Biorad CHEF Genomic DNA Plug Kit). DNA was recovered from agarose plugs according to IrysPrep™ Plug Lysis Long DNA Isolation guidelines (BioNano Genomics). Of the isolated DNA, 300 ng was used for subsequent DNA nicking using *Nt.Bsp*Q1 (NEB) incubating for 2 hours at 50°C. Labeling, repair and staining reactions were done according to IrysPrep™ Assay NLRS (30024D) protocol. Finally, labeled DNA molecules were analysed on a BioNano Genomics Irys instruments with optimized recipes using one Irys chip, one flowcell, 9 runs, with 270 cycles in total.

Data was collected and processed using IrisView software V 2.5 together with a XeonPhi (version v4704) accelerated cluster and special software (both BioNano Genomics, Inc.). A *de novo* map assembly was generated using molecules equal or bigger than 140 Kb, and containing a minimum six labels per molecule. In total, the molecules used for assembly encompassed 291 Mb equivalent space and on average 8 labels per 100Kb molecule size. For the assembly process, stringency settings for ‘alignment’ and ‘refineAlignment’ were set to 1e-8 and 1e-9 respectively. The assembly was performed by applying 4 iterations, where each iteration consisted of an extension and merging step.

Hybrid scaffolding was done using ‘hybrid scaffolding_config_aggressive’ settings of IrysView.

### Genome assembly and scaffolding

The genome assembly workflow began with *de novo* assembly of contigs from the PacBio long reads using two tools, Canu (version (1.5) and Falcon (version 0.5). For each assembler the most important assembly parameterswere systematically varied (Supplementary Methods), as defined by the tool developers, and by consideration of assembly theory (e.g. overlap length, overlap identity for overlap layout consensus assembly). Optimal settings were selected by comparison of assembly statistics (total size assembled and contig N50) and by alignment of Illumina PE data to the assembly with bowtie2 ^27^ (using the ‘very fast’ preset). For all PacBio assemblies the consensus step was performed by running Quiver (Genomic Consensus version 2.3.3) ^28^ (with default parameters) on raw PacBio contigs and using the full 63X of PacBio data.

Assembled contigs were further joined into scaffolds using a combination of BioNano optical mapping data, Hi-C chromatin conformation capture data, and genetic maps. The best assemblies from Canu and Falcon were independently combined with BioNano optical mapping data using the IrysView software to develop the Canu + BioNano(CB) and Falcon + BioNano (FB) assemblies, respectively. The BioNano scaffolding process identified conflicts between the assembled contigs and the optical map, indicating some degree of misassembly in both Canu and Falcon results.

### Assembly Polishing

Pilon (version1.21) ^29^ was run iteratively on the assembly, with Illumina sequence realigned to the polished assembly at each iteration and then alignments passed to Pilon to call the next consensus. Kmer spectrum comparisons were made using the kmer analysis toolkit (KAT) (version 2.3.4) ^23^ and the metric used to assess each iteration was the number of kmers shared between the assembly and the Illumina reads. In a second consensus phase, RNA-Seq reads were aligned as single end (SE), to the genome using Hisat (version 2.1.0) ^29^ with default parameters. This time the effectiveness of consensus calling was assessed by analysis of full length alignments of assembled RNA-Seq transcripts. All transcripts designated as ‘complete’ by Evigene ^30^ were aligned to the genome with BLAT (version 3.4) 31 minimum match identity 90%). Alignments were filtered to retain only full length alignments (i.e. from query start to query end). Finally, the number of gaps in the alignments (query gaps + target gaps) was used as a metric with the rationale that this serves as a proxy for the number of indels in alignments of assembled mRNA sequence.

### Scaffold Validation using a high-density genetic map

A high-density genetic map was developed using a 100 individual ‘Old Home’ × ‘Bartlett’ F_1_ population and the Axiom™ Pear 70K Genotyping Array ^32^, Markers were filtered to have less than 5% missing data and fit segregation ratios of 1:1 and 1:2:1 (α = 0.01). Mapping was conducted in an iterative process using the maximum likelihood algorithm in JoinMap 5 ^33^. After each round of mapping, a graphical genotyping approach was used to identify and fix marker order errors and regions with low marker density caused by segregation distortion. Markers that fitted segregation ratios of 2:1 and 2:3:1 (α = 0.01) were added to the dataset after a high-quality framework map was constructed to improve the low-density regions.

The Bartlett parental map produced by JoinMap included 11,474 markers. This map was used to validate and anchor the scaffolds from both the CB and FB assemblies. SNP probe sequences from the array ^32^ used in the construction of the genetic map were mapped to the assembly with BLAT (version 3.4) ^31^. Alignments were filtered to retain only markers perfectly matched to unique loci in the assembly as well as those with a maximum of two mismatches in the second best hit. The resulting alignments were queried to identify problematic scaffolds mapped with SNP probes from different LGs. The number of scaffolds with SNP probes mapped from different LGs was used as a metric in the quality assessment of the FB and CB assemblies. After selection of the CB assembly, its scaffolds were broken at the 3 positions where SNP mapping switched from one LG to another. Each scaffold breaking was performed by dividing the scaffold at the position 500bp past the last good SNP marker.

### Scaffold clustering and genome anchoring using Hi-C

Hi-C reads were aligned to the polished scaffolds in CB with Bowtie2 (version 2.3.3.1) ^34^. Based on the alignments, CB scaffolds were arranged into 17 ordered and oriented clusters using the LACHESIS software ^35^. As an internal check, the process was completed on two different random 75% sub-samplings of the Hi-C data, as well as on the full data set. The clusters produced by all three of these LACHESIS runs were identical. LACHESIS produces groups of scaffolds which are ordered and oriented relative to each other. These scaffold groupings were compared with the genetic map and the consistency of these sources of information was assessed. The SNP probe mapping at the scaffold validation step was compared with the clusters produced by LACHESIS.

### Illumina assembly

The Illumina data was also assembled on its own, using the de-Bruijn graph based assembler SOAPdenovo2 (version 2.04) ^36^ This assembly was used in various ways during the course of the pear genome project (for further scaffold validation, for training the *ab initio* gene predictors, etc.). The Illumina data was assembled twice. The first pass contigs were screened using the Kraken^37^ software and an index built from the entire RefSeq database. Reads aligning to contaminant contigs were removed and the remaining data was assembled again.

### Repeat Annotation

Repbase (v 16.02; ^38^) was used to identify repeats by using RepeatMasker (version 4.0.5) ^39^. RepeatModeler (version 1.0.8) ^39^ was used to build *de novo* repeats. HODOR sequences ^2^ were identified by blasting the apple HODOR sequence onto the assembly.

### Transcriptome assembly

The 26.6 Gb ‘Bartlett’ RNA-Seq data (SRA accession numbers SRR1572981 to SRR1572991) was assembled *de novo*, using Trinity (version 2.2.0) ^40^ and also genome guided, using both Cufflinks (version 2.2.1) ^41^ and Trinity-GG (version 2.2.0) ^42^. All transcripts from these three assemblies were pooled and input into the EviGene pipeline ^30^ which produces a non-redundant transcript database classified into putative primary and alternative transcripts.

### Gene annotation

Gene prediction was guided by the non-redundant transcriptome assembly, as well as by spliced alignments from three sources: CDS from closely related species (apple and Asian pear), proteins from these and other less related plant species (Arabidopsis, rice, tomato), and RNA-Seq read data aligned onto the genome. All assembled European pear transcripts classified as both full length and primary by EviGene were input to the ORF finder Transdecoder (version 3.0.0) ^43^ to give a set of predicted CDS sequences. These predicted CDS and CDS from closely related species were aligned to the genome using BLAT (version 3.4) ^31^ and Genome Threader (version 1.7.0) ^19^. Protein alignments were performed using Genome Threader. Mapping of all these evidence sources was first made to Illumina contigs and a training set for the training of *ab initio* gene predictors was constructed by manual annotation of genes on these contigs. Both Augustus (version 3.3) ^44^ and Eugene (version 4.2) ^45^ gene predictors were trained using this manually annotated training set.

Spliced alignments of RNA-Seq reads to the genome provide strong evidence for the structure of genes by delineating intron-exon boundaries. RNA-Seq data downloaded from NCBI/SRA were aligned to the pear genome using HiSat (version 2.1.0) ^29^ with custom parameters. This evidence was leveraged by providing Augustus ^44^ with ‘hints’ files detailing the intron-exon boundaries and providing Eugene ^45^ with splice site models generated by the SpliceMachine software (version 1.2) ^46^. Spliced alignments of assembled transcripts were leveraged by passing them to the PASA pipeline (version 2.3.1) ^40,47^ which constructs a genome based transcriptome assembly. PASA assembled transcripts were then processed by Transdecoder to produce a set of ORFs as genome based GFF coordinates.

*Ab initio* gene predictions were performed with Augustus and Eugene using models trained on the manually annotated Illumina sequence. Augustus was executed with hint files conveying information about the spliced mappings of RNA-Seq reads, assembled transcripts, CDS sequences and proteins and the repeat annotation of the genome. Similarly, these supporting hints were supplied to Eugene and the prediction was run on repeat masked sequence (with soft masking). The *ab initio* gene models from Augustus and Eugene were combined with the PASA gene models as well as the gene models produced by Genome Threader alignment of proteins, CDS, and assembled transcripts. The Evidence modeler software (version 1.1.1) ^48^ was used to combine these different gene models and evidence sources. Finally, the Evidence modeler annotation was taken and used to retrain Eugene. A final Eugene iteration using this Evidence modeler annotation as an evidence track helped to clean up the splice boundaries of some coding sequences.

### *K*_S_-based paralog and ortholog age distributions

Paralog age distributions of synonymous substitutions per synonymous site (*K*_S_) were constructed as previously described ^49^, except using PhyML ^50^ instead of average linkage hierarchical clustering for tree construction. Briefly, to build the paranome, an all-against-all BLASTP search was performed with an E-value cutoff of 1 × 10^-10^, followed by gene family construction using the mclblastline pipeline (v10-201, micans.org/mcl ^51^). Gene families larger than 400 members were removed. Each gene family was aligned using MUSCLE (v3.8.31 ^52^), and *K*_S_ estimates for all pairwise comparisons within a gene family were obtained through maximum likelihood (ML) estimation using the CODEML program ^53^ of the PAML package (v4.4c ^54^). Gene families were then subdivided into subfamilies for which *K*_S_ estimates between members did not exceed a value of 5. To correct for the redundancy of *K*_S_ values (a gene family of *n* members produces *n*(*n*–1)/2 pairwise *K*_S_ estimates for *n*–1 retained duplication events), a phylogenetic tree was constructed for each subfamily using PhyML ^50^ under default settings. For each duplication node in the resulting phylogenetic tree, all *m K*_S_ estimates between the two child clades were added to the *K*_S_ distribution with a weight of 1/*m* (where *m* is the number of *K*_S_ estimates for a duplication event), so that the weights of all *K*_S_ estimates for a single duplication event summed to one.

*K*_S_-based ortholog age distributions were constructed by identifying one-to-one orthologs between species using InParanoid ^55^ with default settings, followed by *K*_S_ estimation using the CODEML program as above. Coding sequences for *M.* × *domestica* and *P.* × *bretschneideri* were obtained from the apple GDDH13 genome project ^56^ and from the PLAZA dicot database ^57^.

### Gene family analysis

Proteins of *P. x bretschneideri* ^8^, *M. x domestica* ^2^, *F. vesca* ^14^, P. *persica* ^15^, *R. chinensis* ^16^, *R. occidentalis* ^1^, *V. vinifera* ^17^, and *A. thaliana* ^18^ were collected for all-against-all alignment to predicted proteins for *P. communis* with BLASTP ^57^ (evalue < 10^-4^). These alignments were passed to the OrthoFinder ^58^ software, which was run with default parameters.

### Collinearity and synteny

All-against-all protein alignments were also passed to the MCScanX software ^59^ to identify collinearity blocks. Self-collinearity of pear was plotted using the circle_plotter program bundled with MCScanX, after rebuilding the collinearity blocks with a minimum block size of 20 to reduce the noise level. Duplication depth of strawberry homologs in pear was counted with the dissect_multiple_alignment script bundled with MCScanX. DNA level synteny between *P. communis, P.* x *bretschneideri, M. × domestica*, and the two assembly versions for *P. communis* were all plotted using DGenie ^60^ with default parameters.

## Data availability

Genome assembly and gene predictions have been submitted to the Genome Database for Rosaceae^61^ and are freely available at the url https://www.rosaceae.org/species/pyrus/pyrus_communis/genome_v2.0 alongside tools like JBrowse and BLAST. The community annotation portal for *P. communis* will be made available for read only access at https://bioinformatics.psb.ugent.be/orcae/ following publication. To participate in the ongoing manual annotation efforts please contact Yves Van de Peer.

## Acknowledgments

The authors wish to thank the INRA experimental horticultural unit (UE Horti) for having maintained W65/Bartlett.DH in the field for 20 years, Elisa Ravon (INRA-IRHS) for her technical assistance in the initial DNA extraction and genotyping of W65, the California Pear Advisory Board and the Pear Pest Management Research Fund for providing funding to perform PacBio sequencing of the DH and genotyping of the mapping population and the Provincia Autonoma di Trento that partially funded this project. YVdP acknowledges support from the European Union Seventh Framework Programme (FP7/2007-2013) under European Research Council Advanced Grant Agreement 322739 – DOUBLEUP.

## Author contributions

GL and LB assembled the genome and performed assembly QA. LVB and ES performed Bionano optical mapping. GL and SR performed gene annotation and functional annotation. GL performed kmer analysis, repeat annotation, orthology analysis, synteny analysis, inter assembly comparisons, transcriptome assembly, Hi-C data analysis, interpreted the results and wrote the manuscript. CL prepared Hi-C libraries. CD contributed to the BUSCO analysis, to interspecies synteny analysis and performed contamination screening of Illumina assemblies. RL performed analysis of the whole genome duplication. Under the scientific authority of YL, PG identified, checked, doubled, multiplied and maintained W65/Bartlett.DH since 1996, with DNA finally extracted by JMC. SM, MT, JZ, BN, DC and CD contributed linkage maps. DN, MT, DC, CED, SM and RV contributed funding toward the sequencing. GL, SR, LB, MT, DC, DN, CD, YVDP and RV designed the project. DC, LB, MT, CD, SM, JZ, RL, SR, CL, ES, AC, SG and YVDP contributed to writing the manuscript. All authors approved the final manuscript.

