## Supplementary figures and tables for "Pseudo-chromosome length genome assembly of a double haploid ‘Bartlett’ pear (*Pyrus communis* L.)"

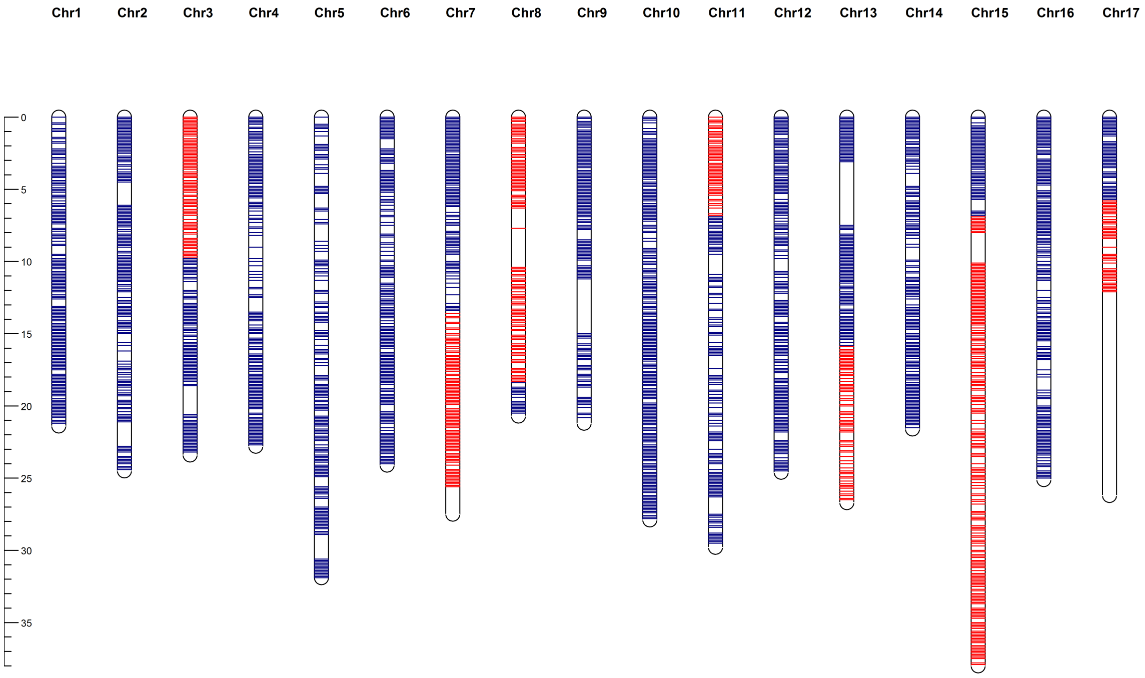


**Figure S1: Haplotype map** for the BartlettDHv2.0. The blue and red colors represent the two different alleles of the SNPs of Bartlett.

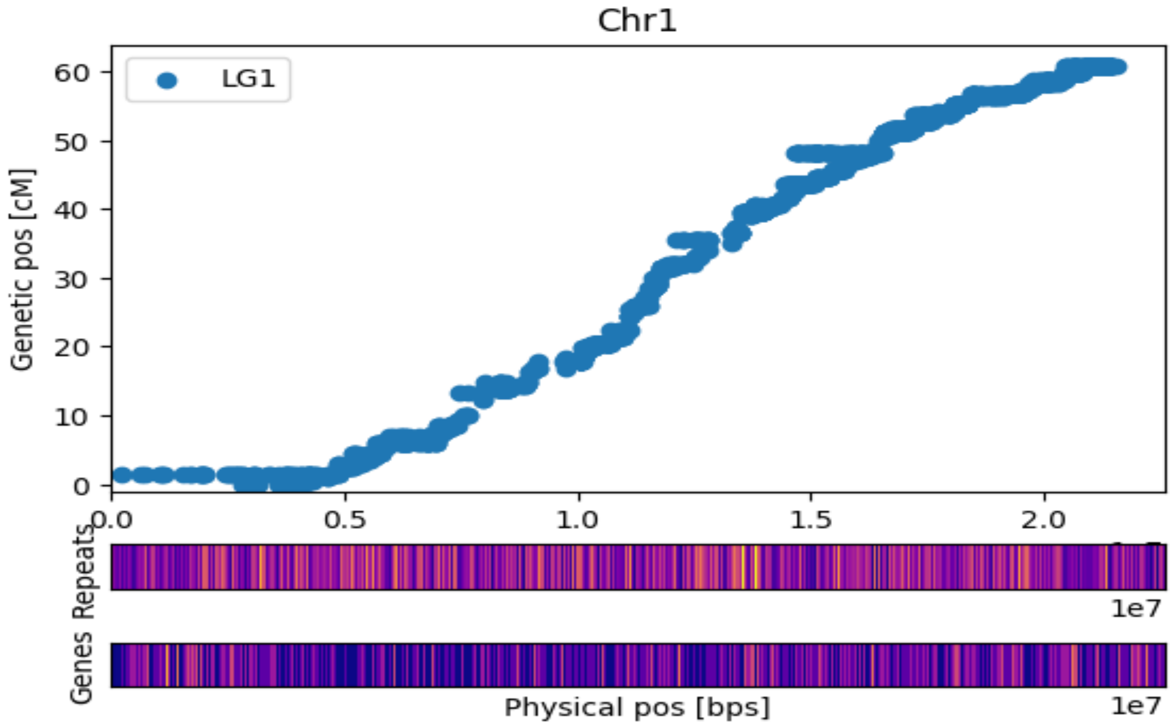


**Figure S2: Marey plot of Chr1 with heatmap of Dispersed Repeats and Genes in bins of 200kb. The lighter the color the more elements are present. Genetic positions refer to the high-density map of Bartlett. Dots represent the genetic and physic position (on of BartlettDHv2.0) of 11,474 SNPs.**

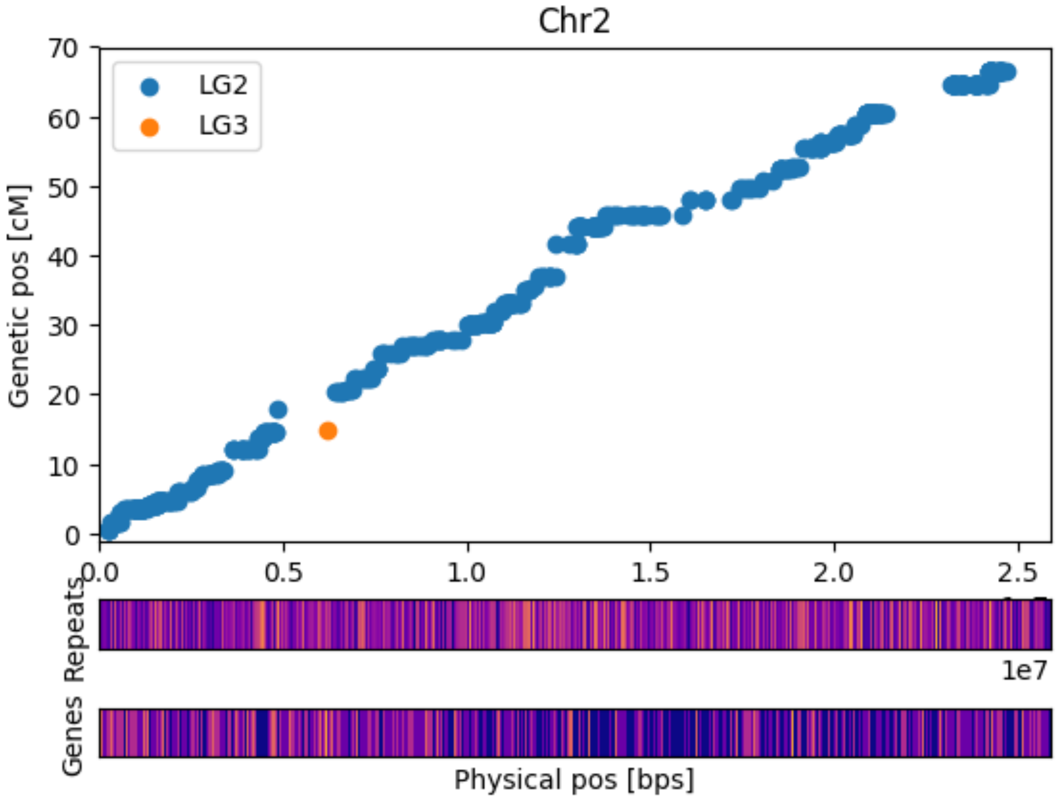


**Figure S3: Marey plot of Chr2 with heatmap of Dispersed Repeats and Genes in bins of 200kb. The lighter the color the more elements are present. Genetic positions refer to the high-density map of Bartlett. Dots represent the genetic and physic position (on of BartlettDHv2.0) of 11,474 SNPs.**

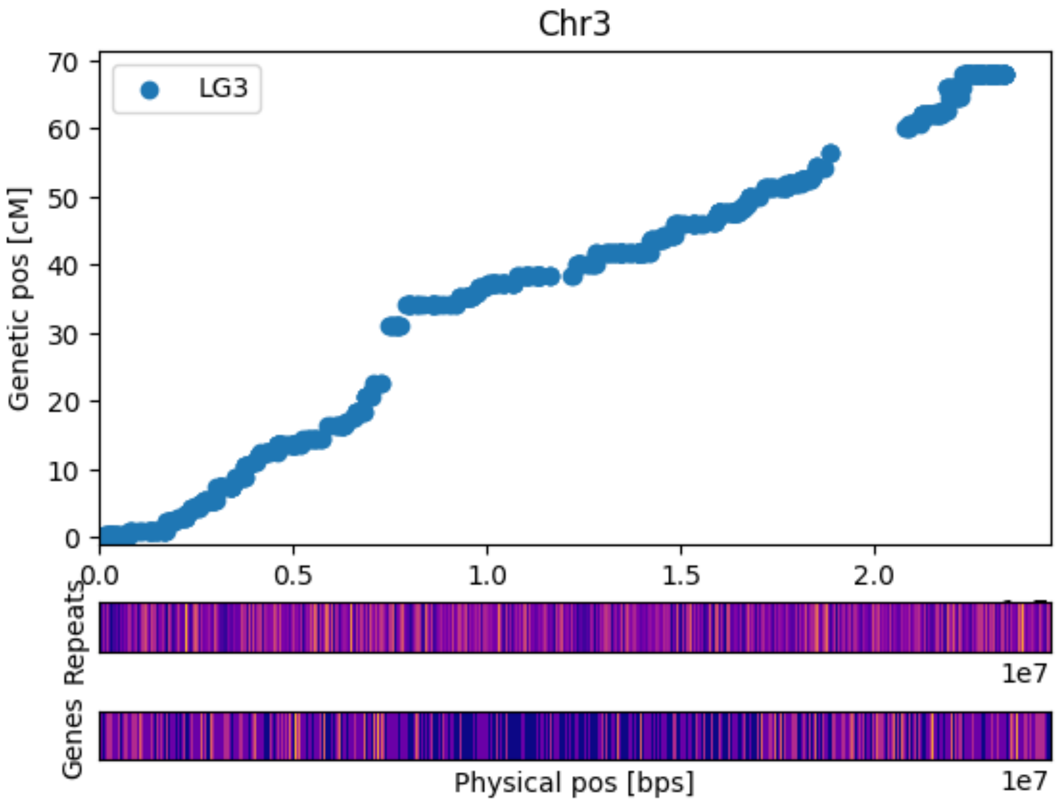


**Figure S4: Marey plot of Chr3 with heatmap of Dispersed Repeats and Genes in bins of 200kb. The lighter the color the more elements are present. Genetic positions refer to the high-density map of Bartlett. Dots represent the genetic and physic position (on of BartlettDHv2.0) of 11,474 SNPs.**

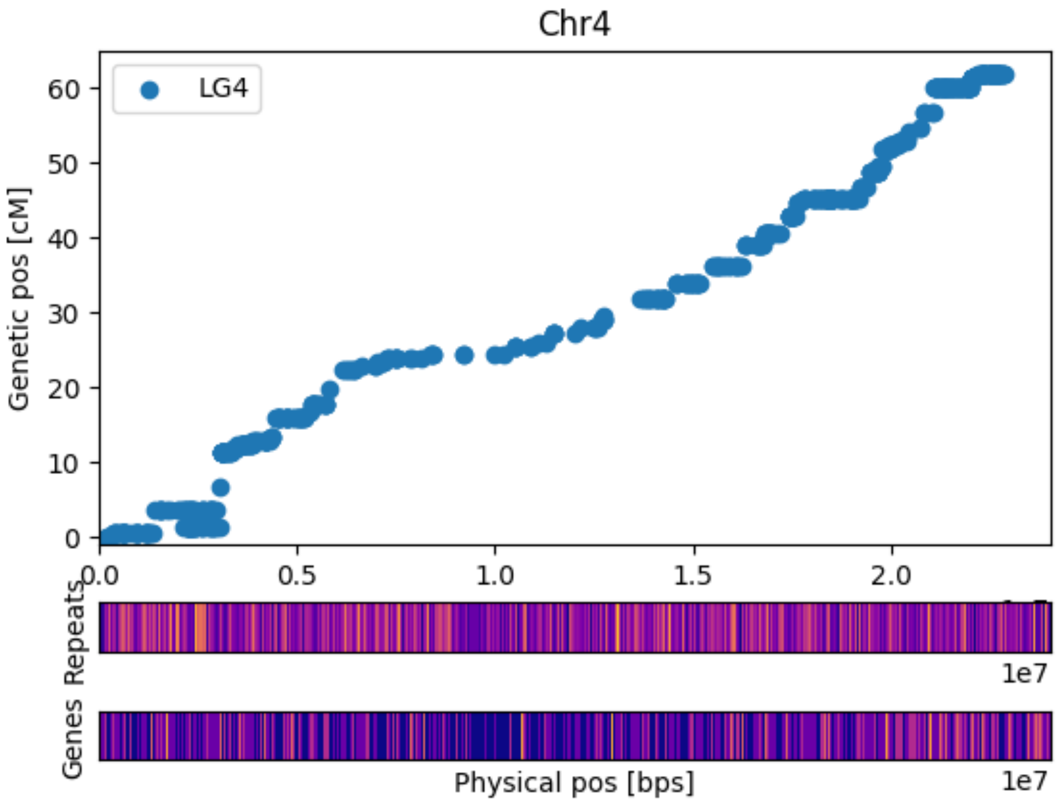


**Figure S5: Marey plot of Chr4 with heatmap of Dispersed Repeats and Genes in bins of 200kb. The lighter the color the more elements are present. Genetic positions refer to the high-density map of Bartlett. Dots represent the genetic and physic position (on of BartlettDHv2.0) of 11,474 SNPs.**

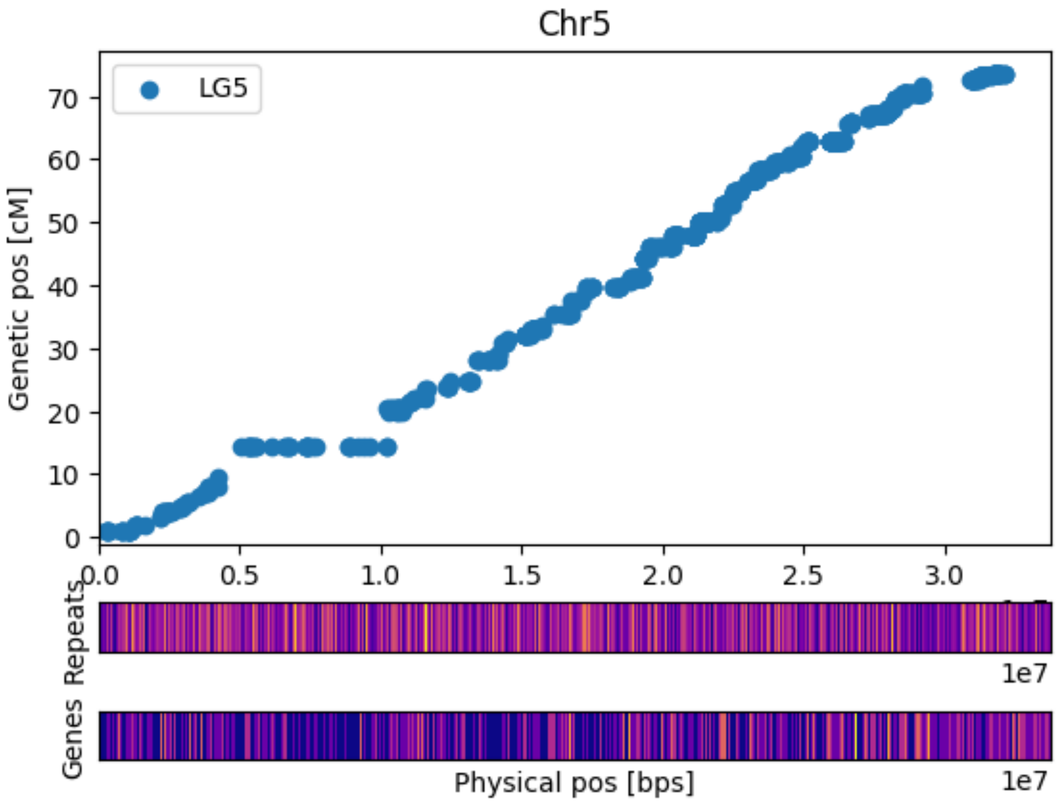


**Figure S6: Marey plot of Chr5 with heatmap of Dispersed Repeats and Genes in bins of 200kb. The lighter the color the more elements are present. Genetic positions refer to the high-density map of Bartlett. Dots represent the genetic and physic position (on of BartlettDHv2.0) of 11,474 SNPs.**

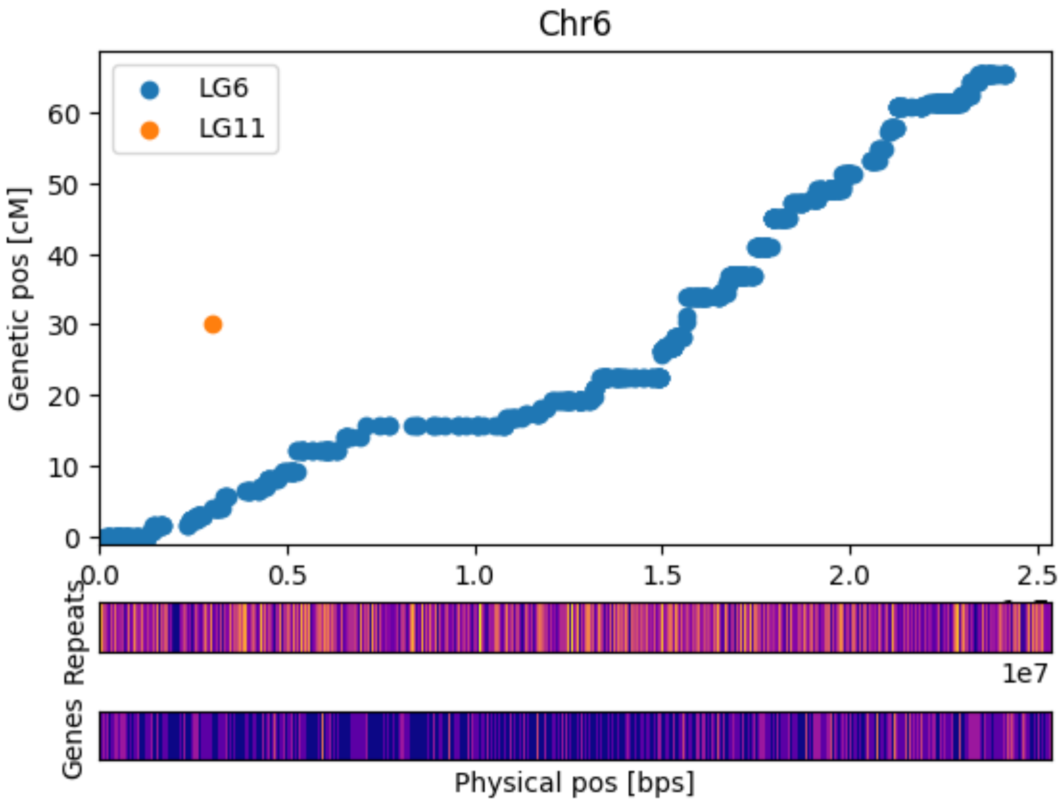


**Figure S7: Marey plot of Chr6 with heatmap of Dispersed Repeats and Genes in bins of 200kb. The lighter the color the more elements are present. Genetic positions refer to the high-density map of Bartlett. Dots represent the genetic and physic position (on of BartlettDHv2.0) of 11,474 SNPs.**

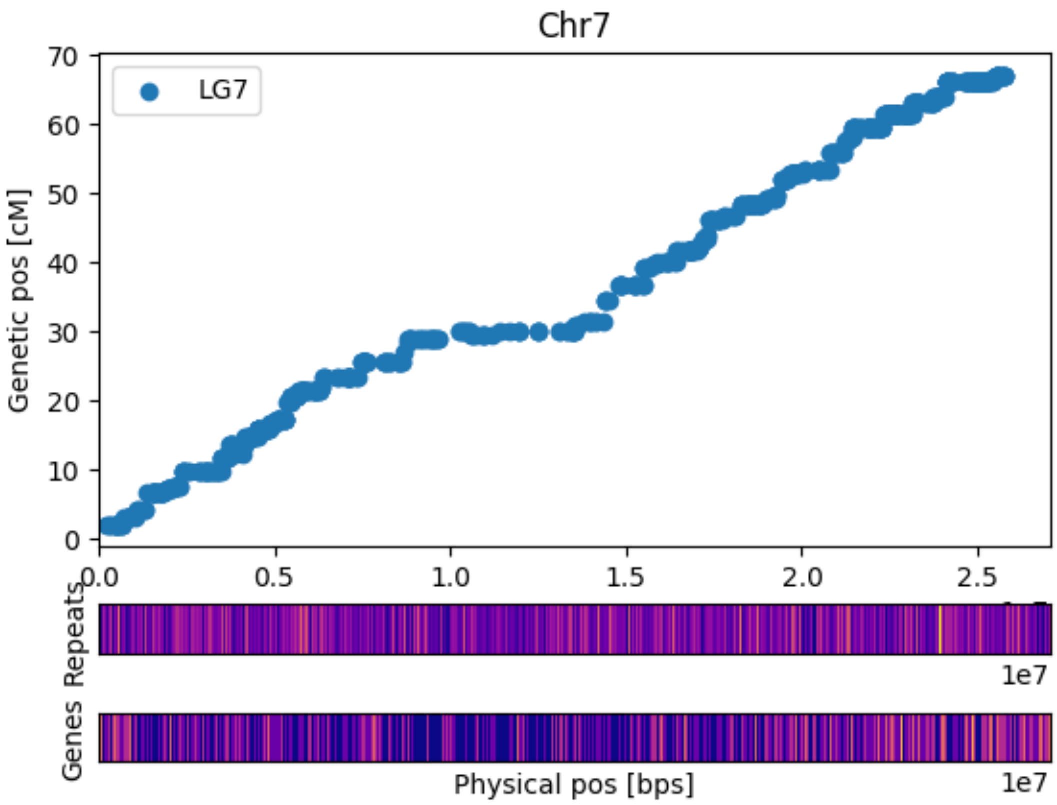


**Figure S8: Marey plot of Chr7 with heatmap of Dispersed Repeats and Genes in bins of 200kb. The lighter the color the more elements are present. Genetic positions refer to the high-density map of Bartlett. Dots represent the genetic and physic position (on of BartlettDHv2.0) of 11,474 SNPs.**

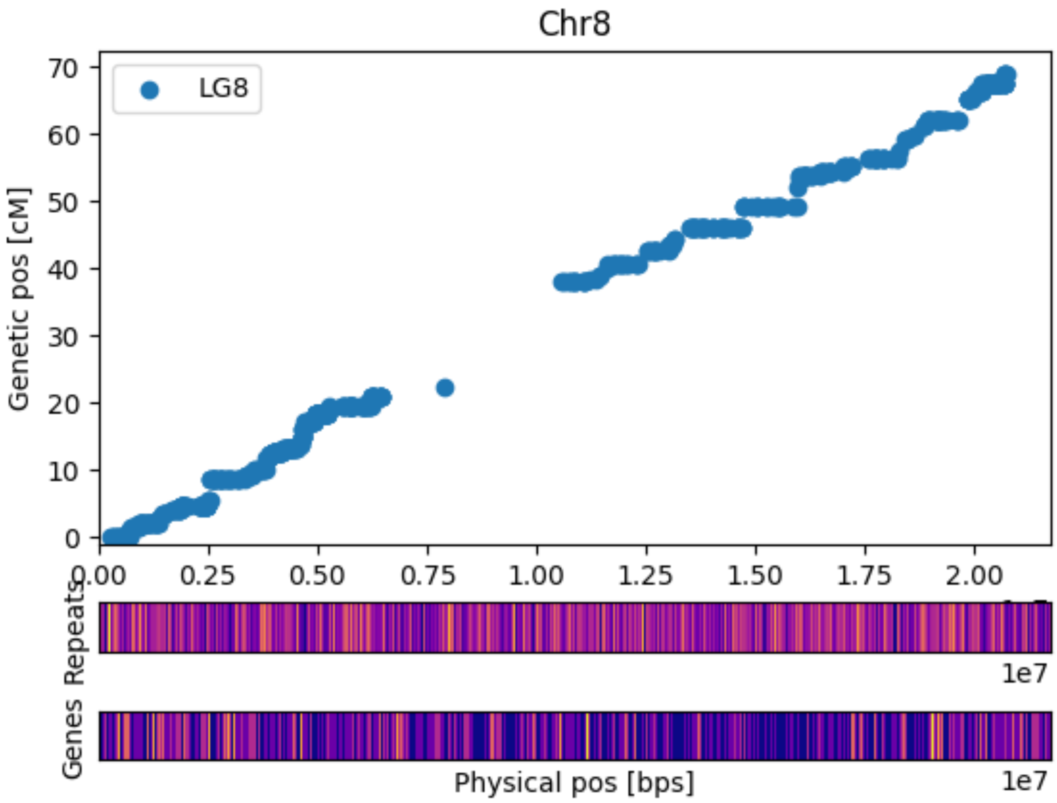


**Figure S9: Marey plot of Chr8 with heatmap of Dispersed Repeats and Genes in bins of 200kb. The lighter the color the more elements are present. Genetic positions refer to the high-density map of Bartlett. Dots represent the genetic and physic position (on of BartlettDHv2.0) of 11,474 SNPs.**

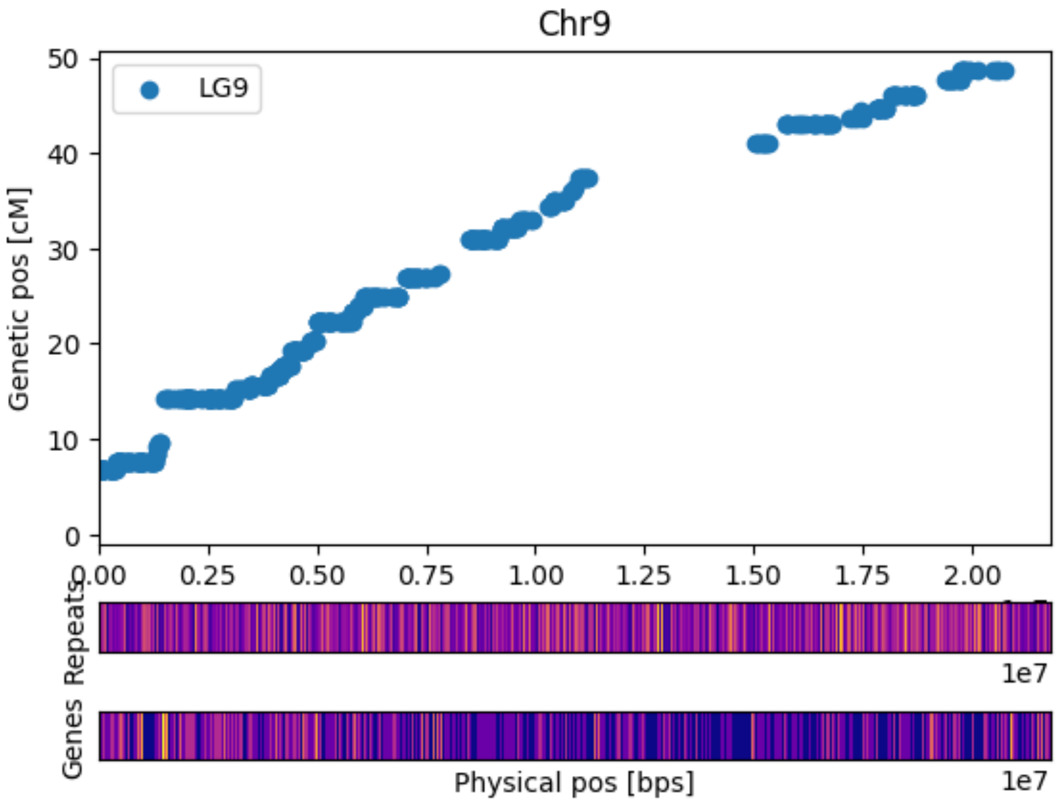


**Figure S10: Marey plot of Chr9 with heatmap of Dispersed Repeats and Genes in bins of 200kb. The lighter the color the more elements are present. Genetic positions refer to the high-density map of Bartlett. Dots represent the genetic and physic position (on of BartlettDHv2.0) of 11,474 SNPs.**

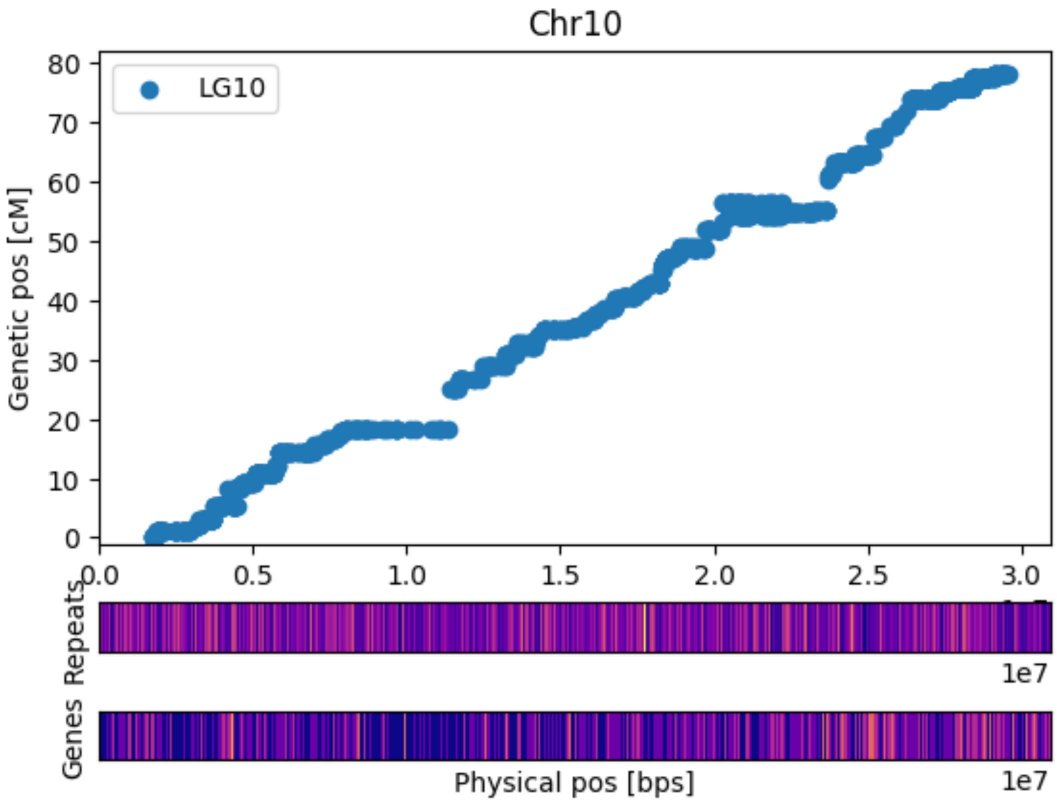


**Figure S11: Marey plot of Chr10 with heatmap of Dispersed Repeats and Genes in bins of 200kb. The lighter the color the more elements are present. Genetic positions refer to the high-density map of Bartlett. Dots represent the genetic and physic position (on of BartlettDHv2.0) of 11,474 SNPs.**

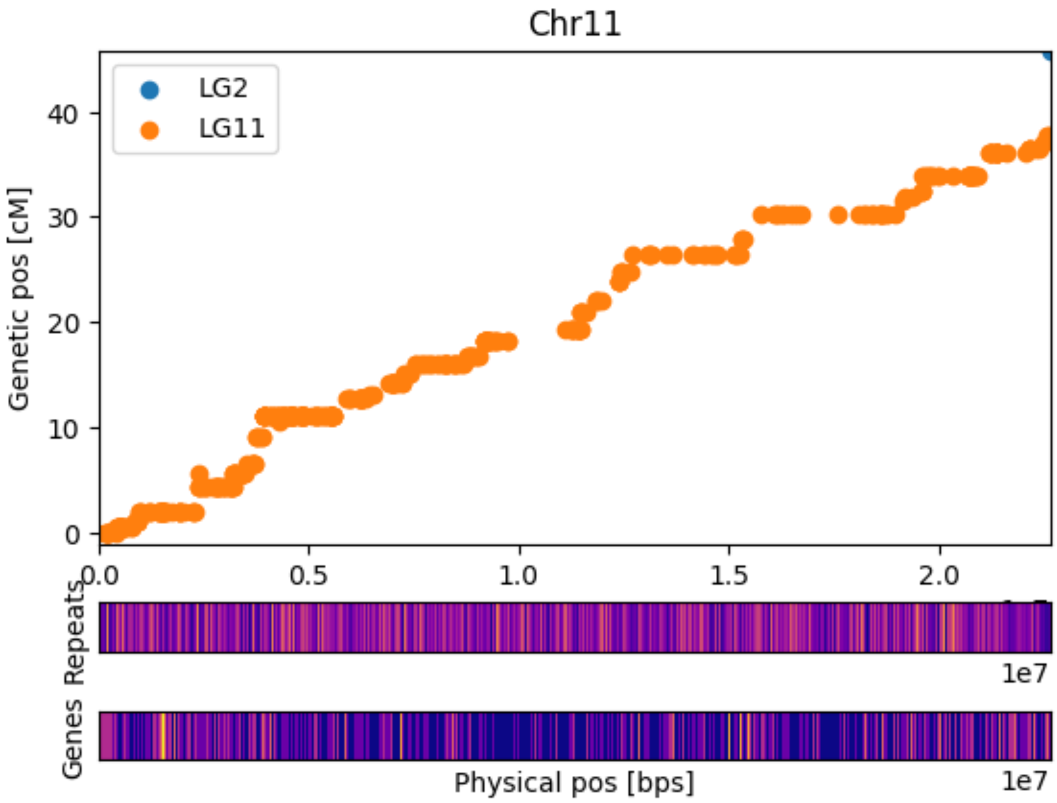


**Figure S12: Marey plot of Chr11 with heatmap of Dispersed Repeats and Genes in bins of 200kb. The lighter the color the more elements are present. Genetic positions refer to the high-density map of Bartlett. Dots represent the genetic and physic position (on of BartlettDHv2.0) of 11,474 SNPs.**

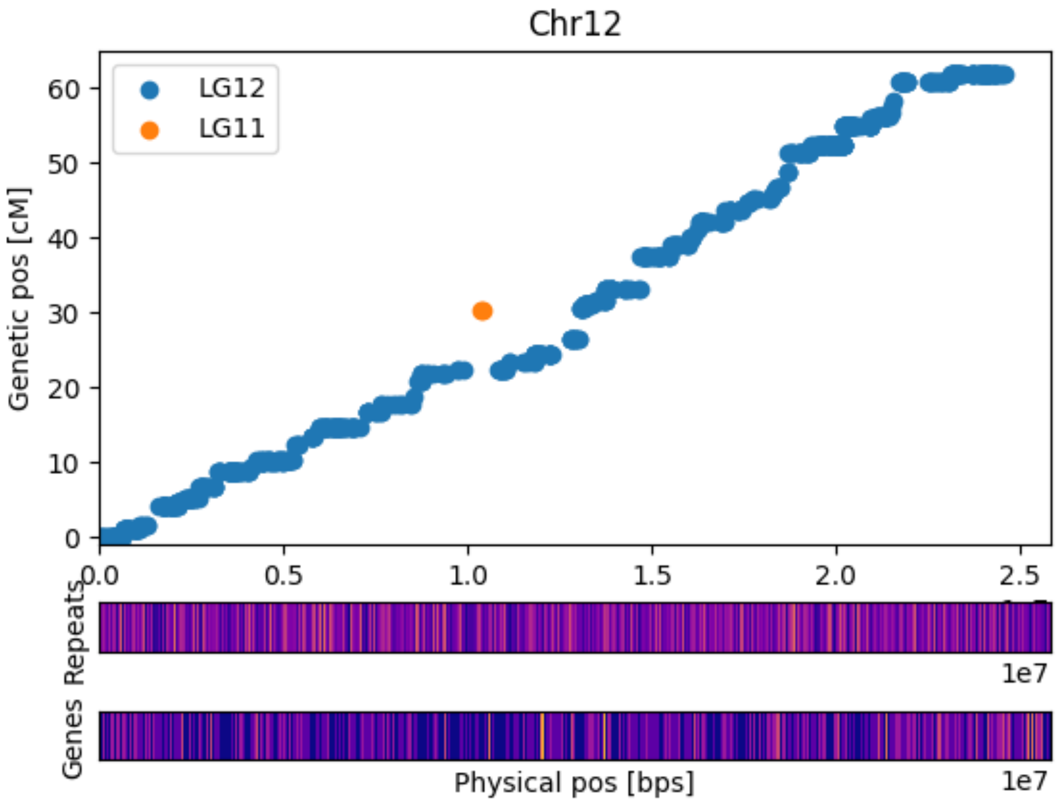


**Figure S13: Marey plot of Chr12 with heatmap of Dispersed Repeats and Genes in bins of 200kb. The lighter the color the more elements are present. Genetic positions refer to the high-density map of Bartlett. Dots represent the genetic and physic position (on of BartlettDHv2.0) of 11,474 SNPs.**

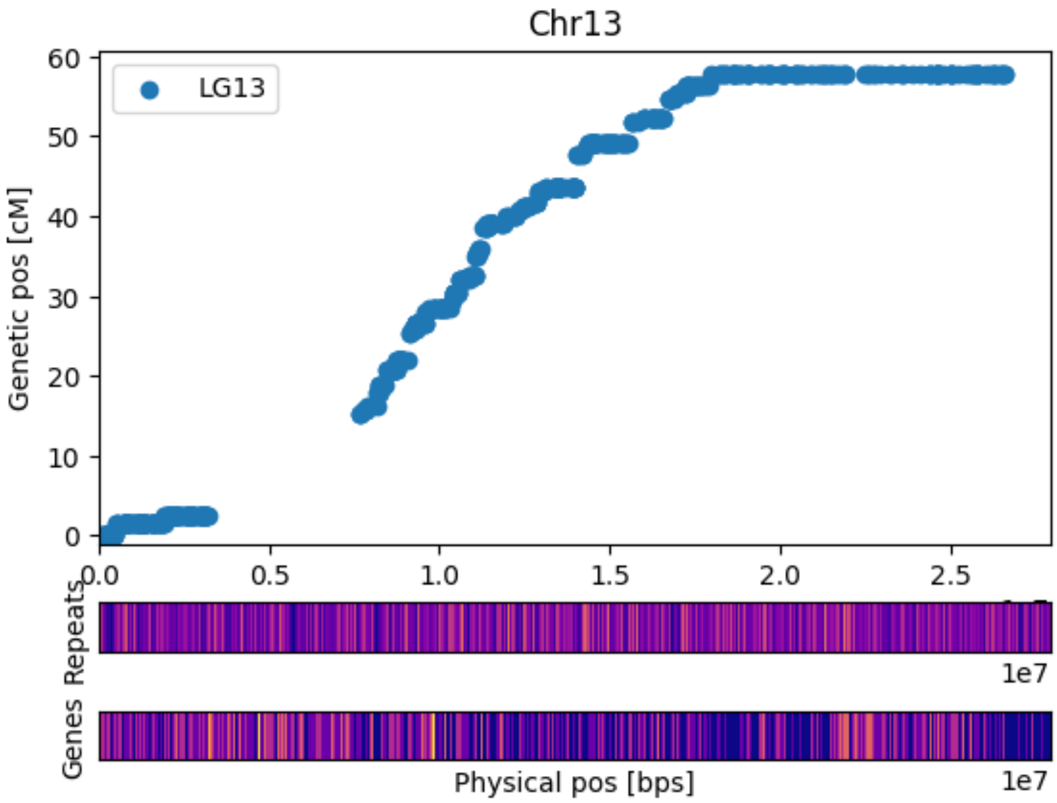


**Figure S14: Marey plot of Chr13 with heatmap of Dispersed Repeats and Genes in bins of 200kb. The lighter the color the more elements are present. Genetic positions refer to the high-density map of Bartlett. Dots represent the genetic and physic position (on of BartlettDHv2.0) of 11,474 SNPs.**

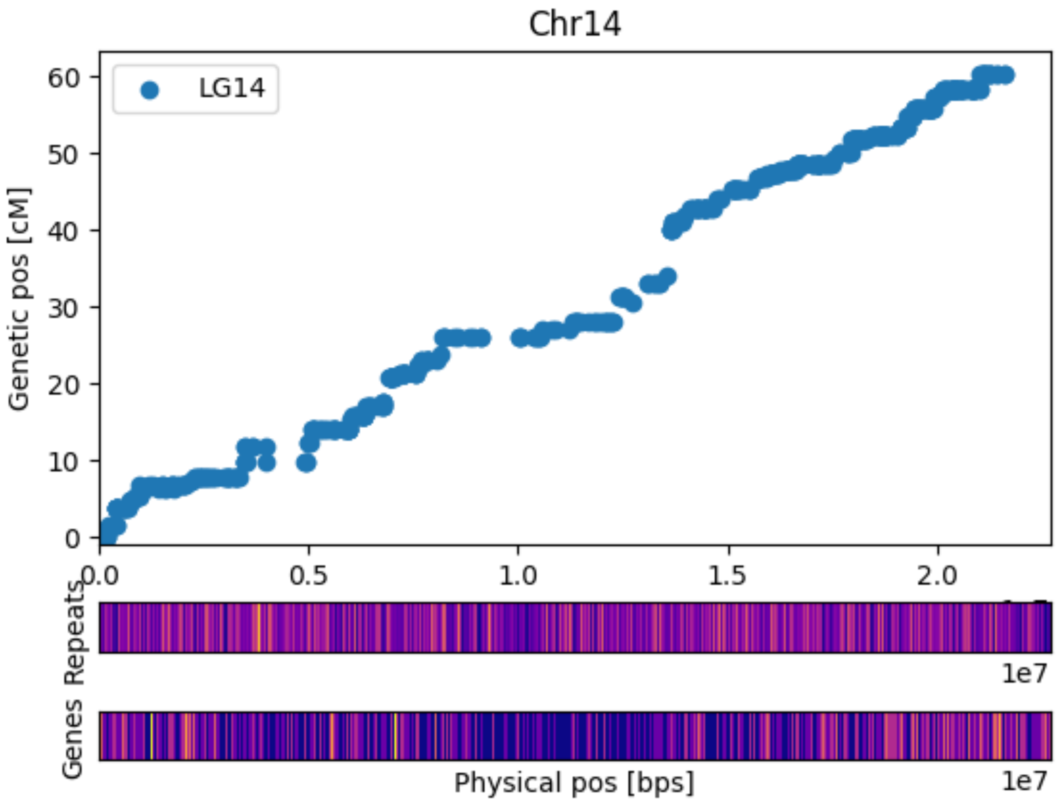


**Figure S15: Marey plot of Chr14 with heatmap of Dispersed Repeats and Genes in bins of 200kb. The lighter the color the more elements are present. Genetic positions refer to the high-density map of Bartlett. Dots represent the genetic and physic position (on of BartlettDHv2.0) of 11,474 SNPs.**

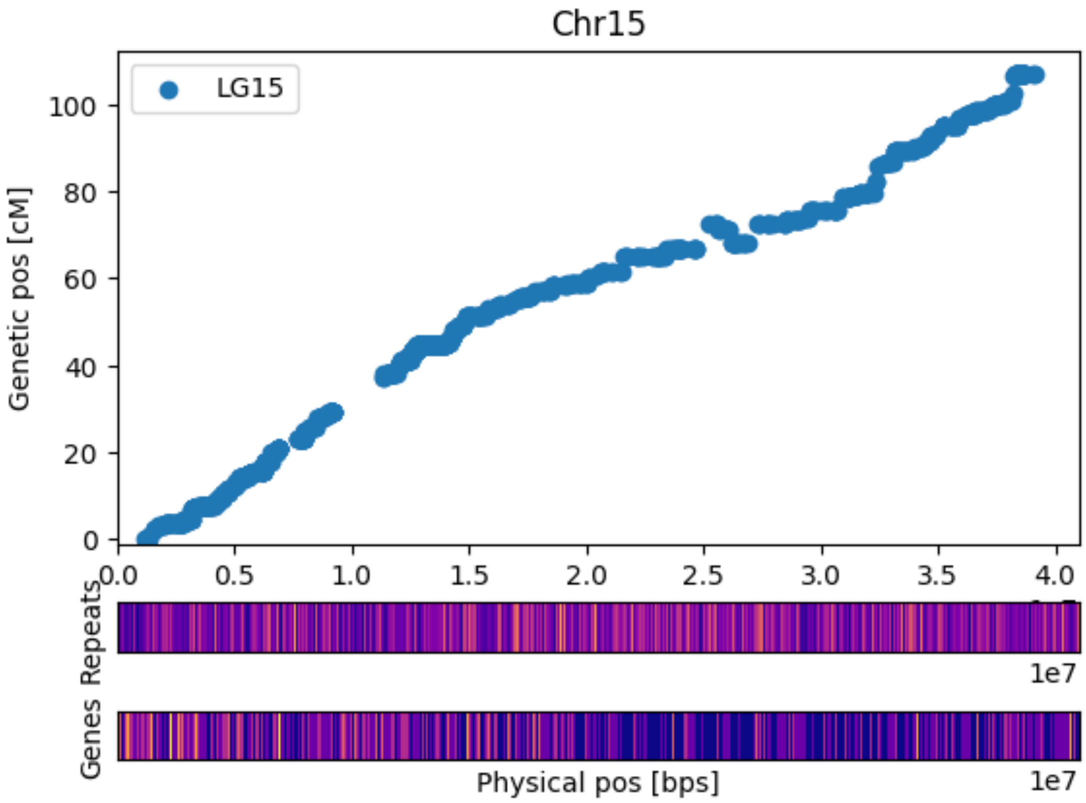


**Figure S16: Marey plot of Chr15 with heatmap of Dispersed Repeats and Genes in bins of 200kb. The lighter the color the more elements are present. Genetic positions refer to the high-density map of Bartlett. Dots represent the genetic and physic position (on of BartlettDHv2.0) of 11,474 SNPs.**

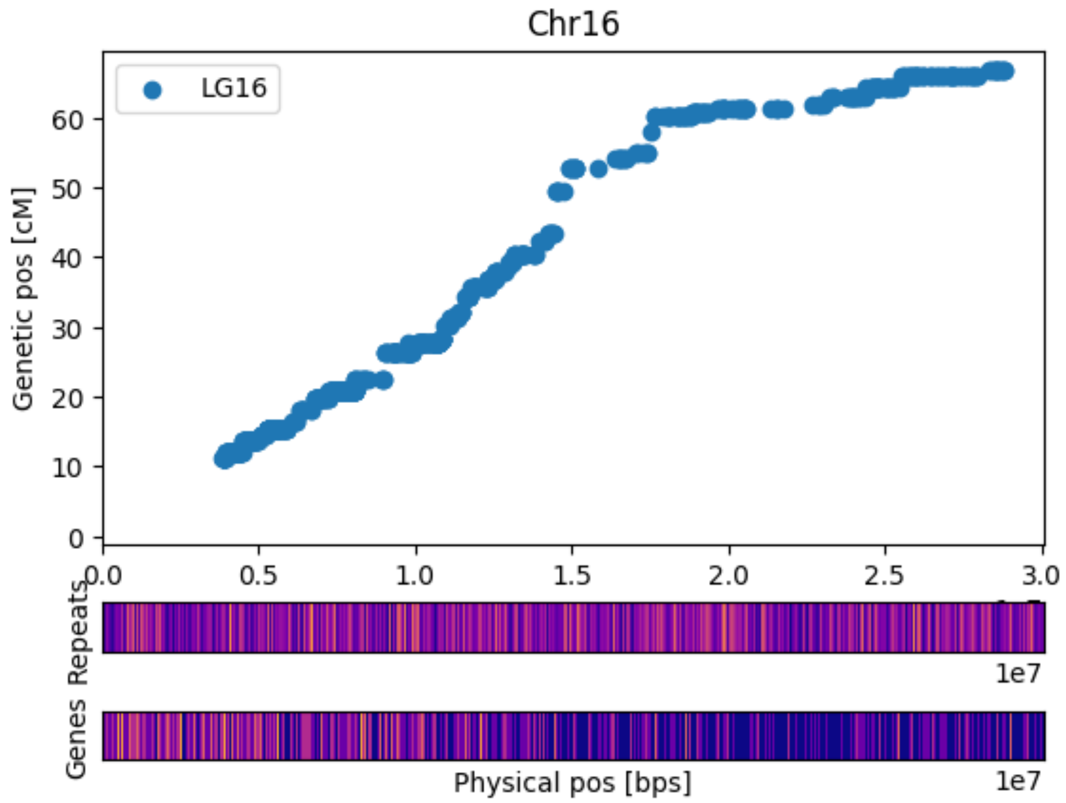


**Figure S17: Marey plot of Chr16 with heatmap of Dispersed Repeats and Genes in bins of 200kb. The lighter the color the more elements are present. Genetic positions refer to the high-density map of Bartlett. Dots represent the genetic and physic position (on of BartlettDHv2.0) of 11,474 SNPs.**

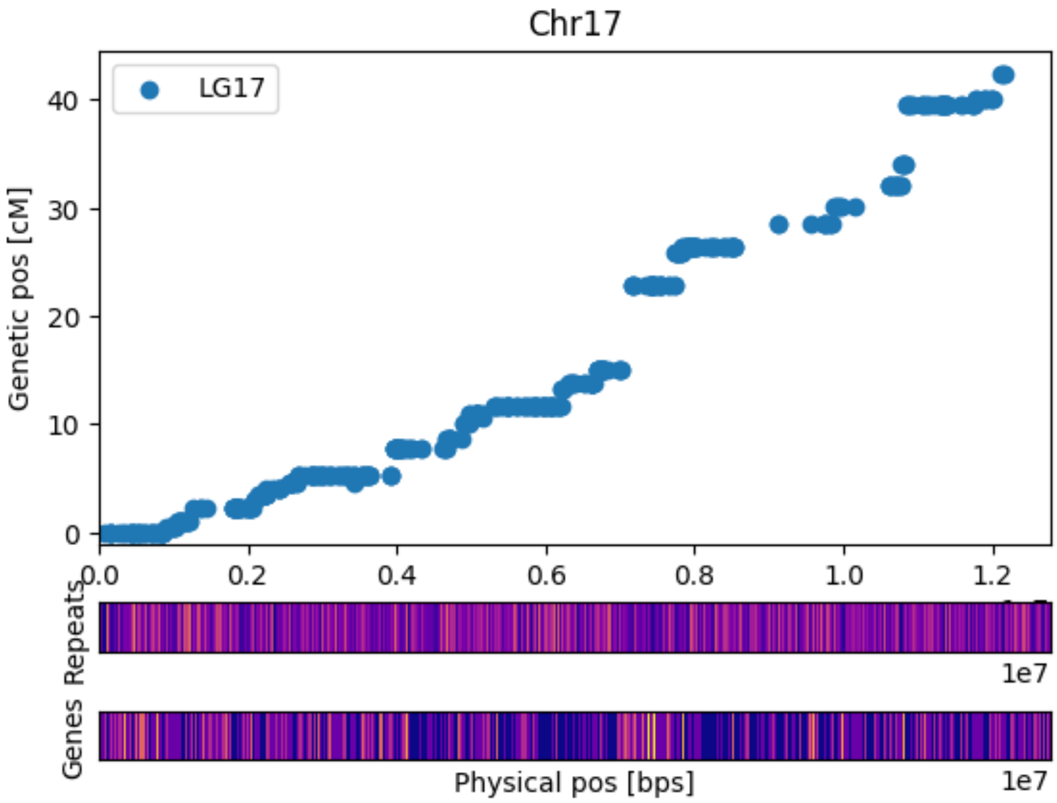


**Figure S18: Marey plot of Chr17 with heatmap of Dispersed Repeats and Genes in bins of 200kb. The lighter the color the more elements are present. Genetic positions refer to the high-density map of Bartlett. Dots represent the genetic and physic position (on of BartlettDHv2.0) of 11,474 SNPs. This Marey plot only shows data for the first 12Mbps of Chr17. Markers in the remaining region show a high level of segregation distortion (and therefore were not mapped genetically) that should be further investigated.**

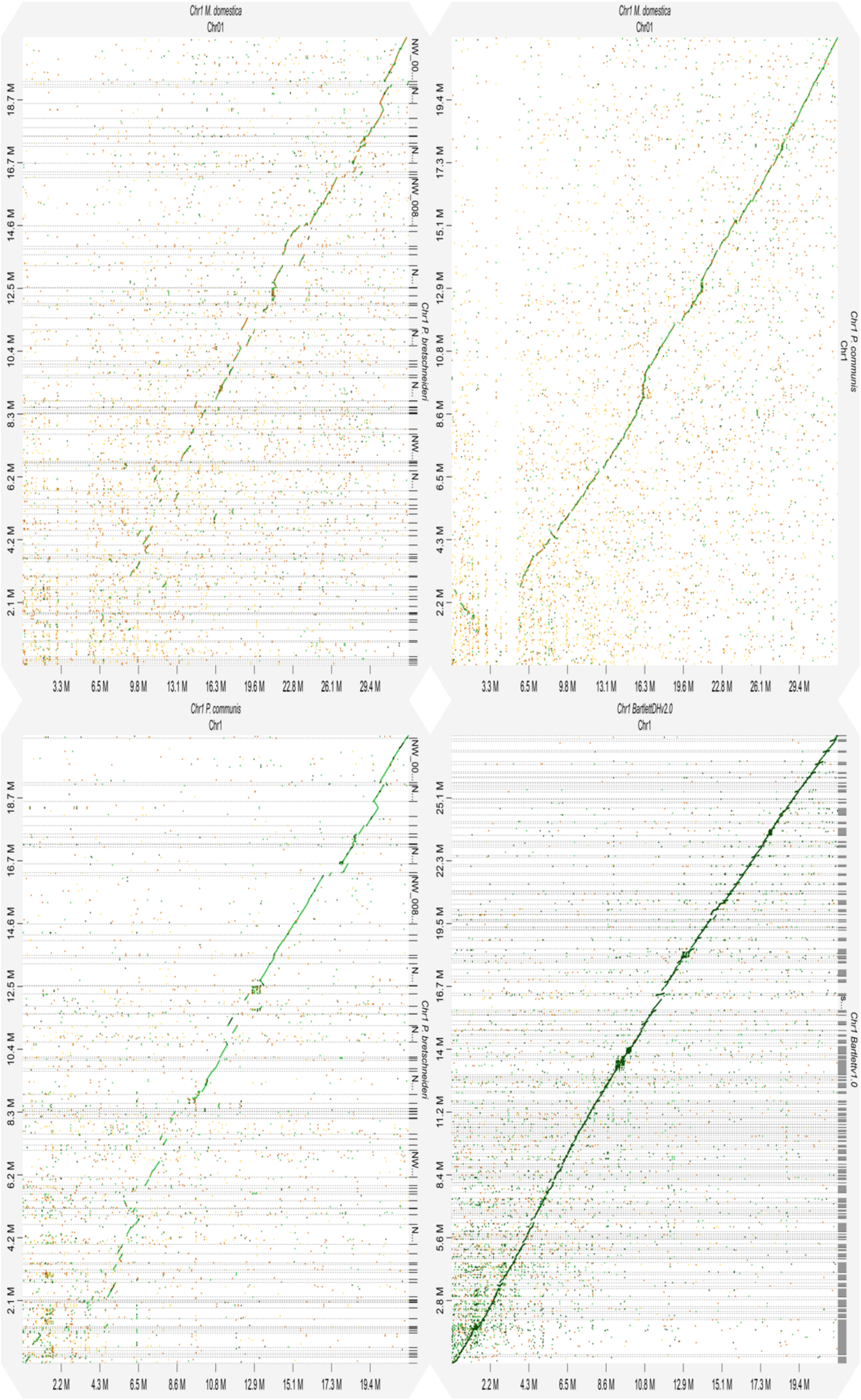


**Figure S19. Chromosome 1 alignment alignment dot plots. Dot plots are produced using the DGENIE software[60]and alignments with minimap2 (v2.16).**

(Fig S19a) Alignment of Chromosome 1 *P. x bretschneideri* to *P. communis* (top left)

(Fig S19b) Alignment of Chromosome 1 *P. communis* to *M. domestica* (top right)

(Fig S19c) Alignment of Chromosome 1 *P. x bretschneideri* to *M. domestica* (bottom left)

(Fig S19d) Alignment of Chromosome 1 *P. communis* of Bartlettv1.0 to of BartlettDHv2.0

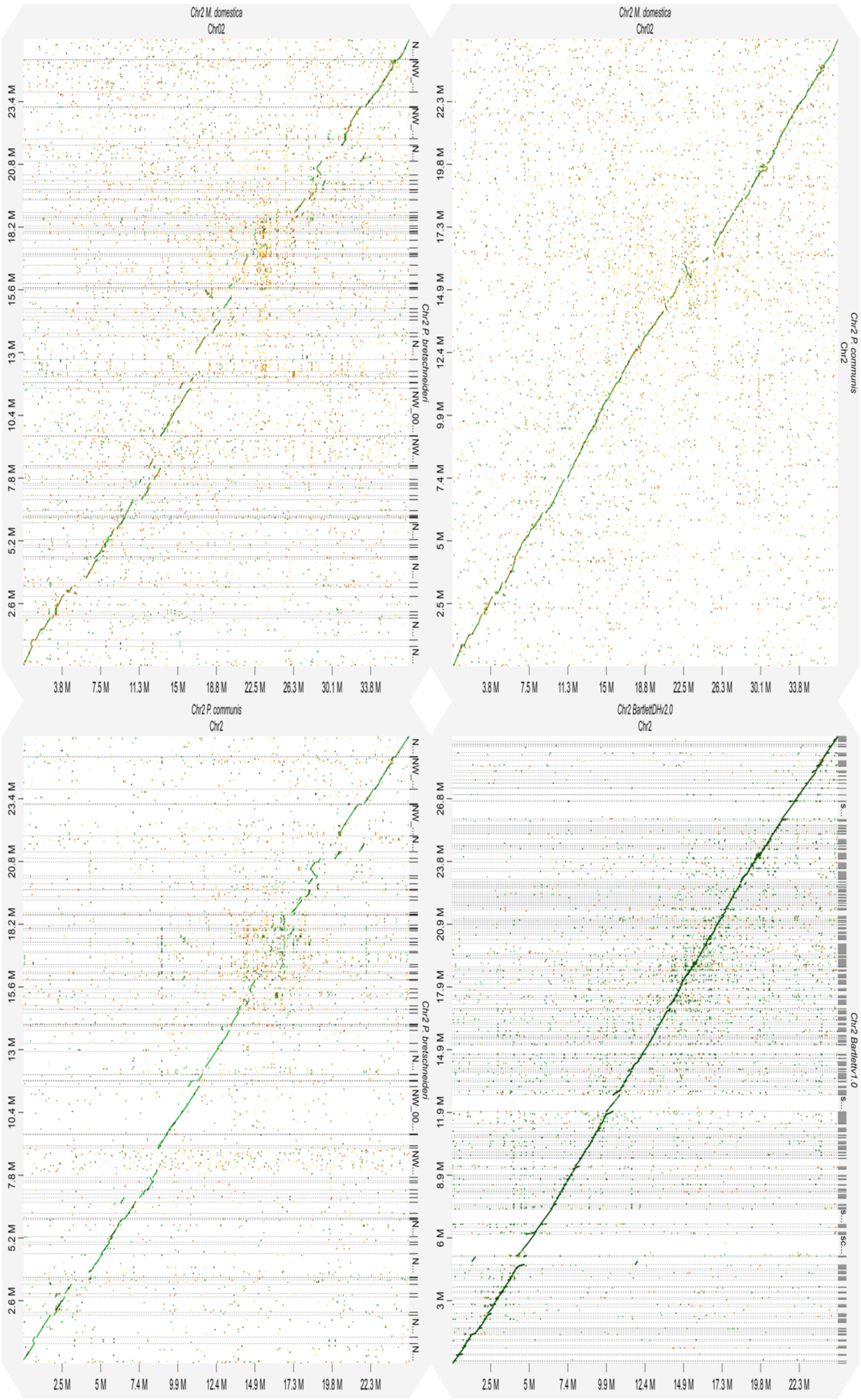


**Figure S20. Chromosome 2 alignment alignment dot plots. Dot plots are produced using the DGENIE software[60]and alignments with minimap2 (v2.16).**

(Fig S20a) Alignment of Chromosome 2 *P. x bretschneideri* to *P. communis* (top left)

(Fig S20b) Alignment of Chromosome 2 *P. communis* to *M. domestica* (top right)

(Fig S20c) Alignment of Chromosome 2 *P. x bretschneideri* to *M. domestica* (bottom left)

(Fig S20d) Alignment of Chromosome 2 *P. communis* of Bartlettv1.0 to of BartlettDHv2.0

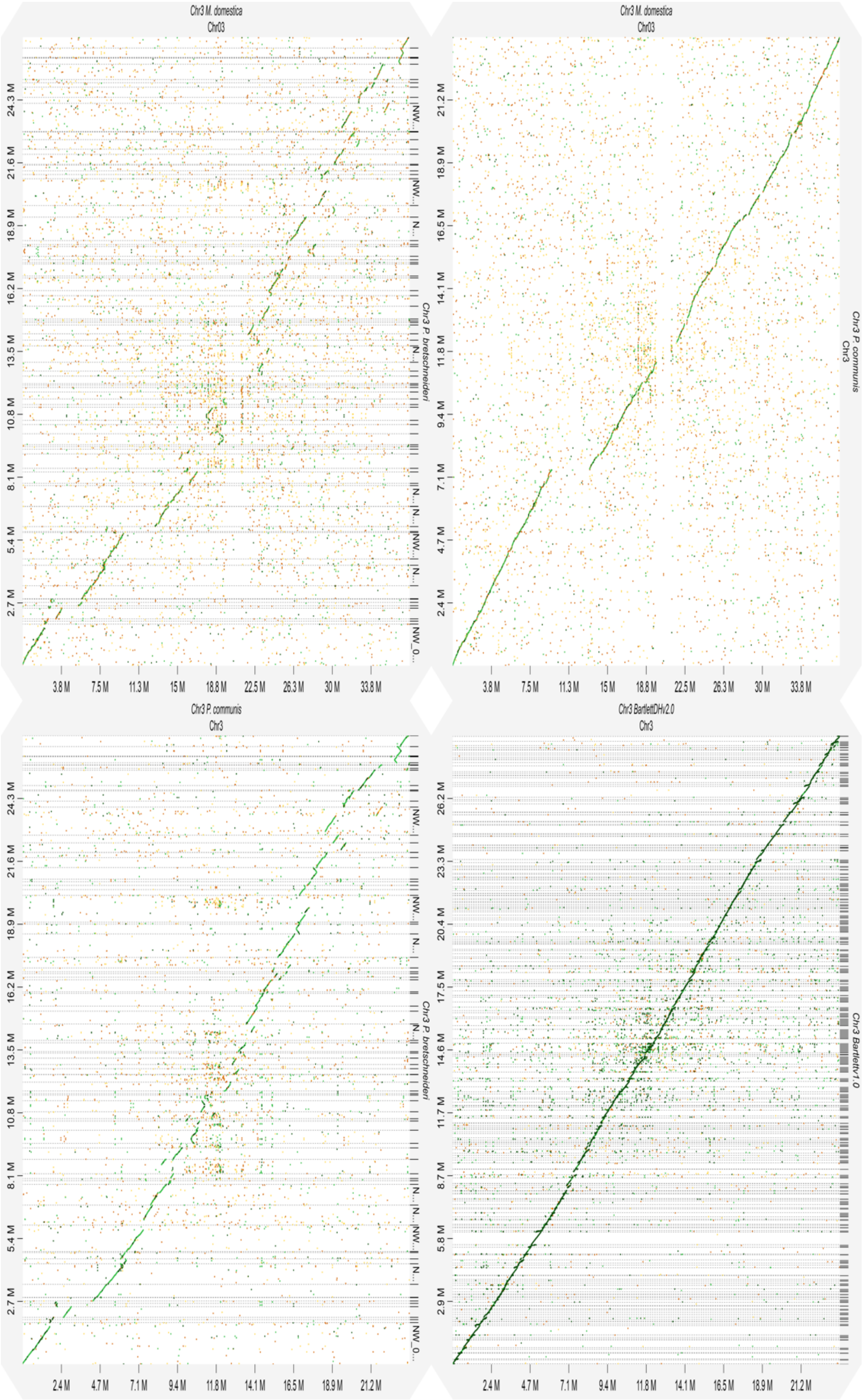


**Figure S21. Chromosome 3 alignment alignment dot plots. Dot plots are produced using the DGENIE software[60]and alignments with minimap2 (v2.16).**

(Fig S21a) Alignment of Chromosome 3 *P. x bretschneideri* to *P. communis* (top left)

(Fig S21b) Alignment of Chromosome 3 *P. communis* to *M. domestica* (top right)

(Fig S21c) Alignment of Chromosome 3 *P. x bretschneideri* to *M. domestica* (bottom left)

(Fig S21d) Alignment of Chromosome 3 *P. communis* of Bartlettv1.0 to of BartlettDHv2.0

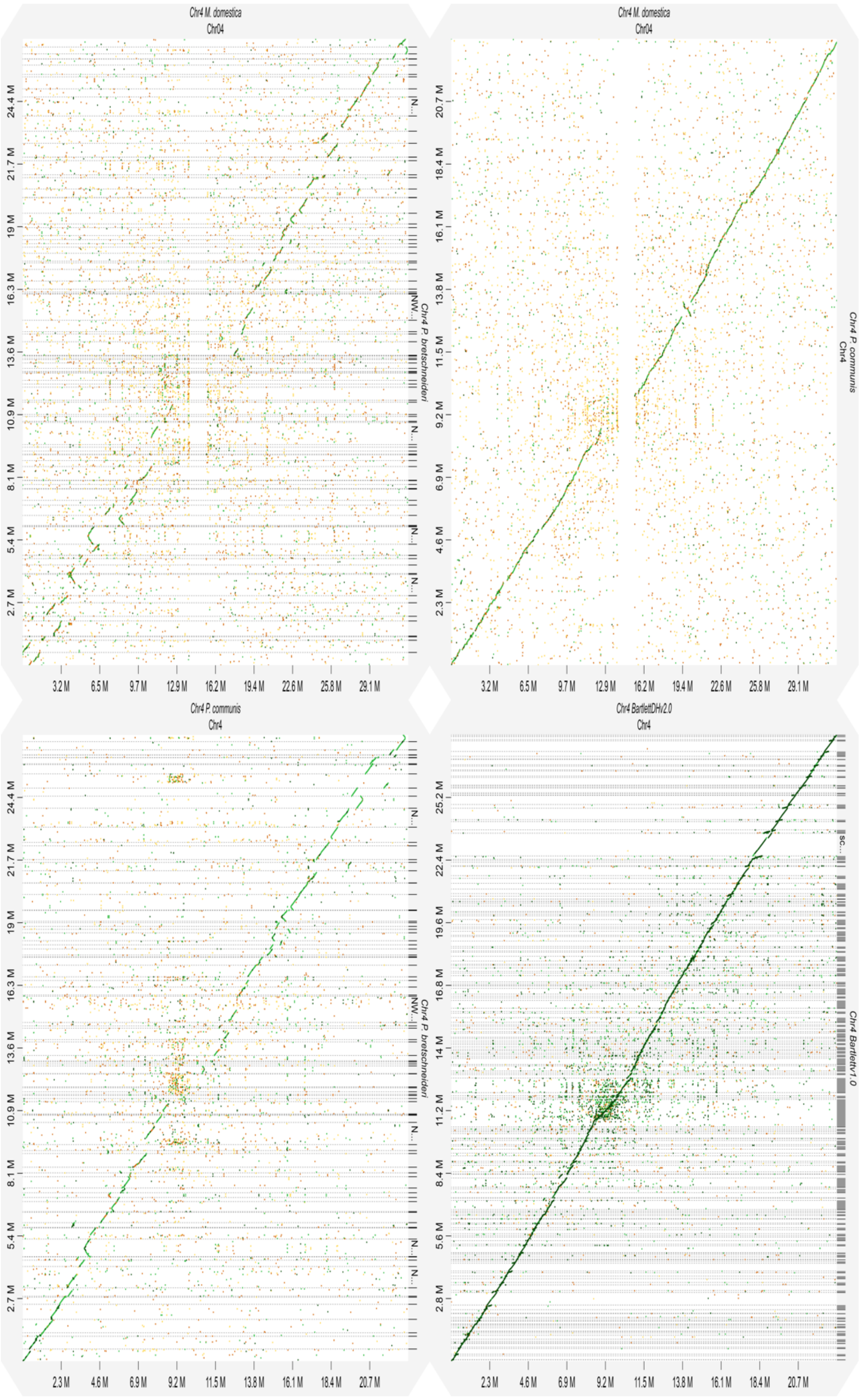


**Figure S22. Chromosome 4 alignment alignment dot plots. Dot plots are produced using the DGENIE software[60]and alignments with minimap2 (v2.16).**

(Fig S22a) Alignment of Chromosome 4 *P. x bretschneideri* to *P. communis* (top left)

(Fig S22b) Alignment of Chromosome 4 *P. communis* to *M. domestica* (top right)

(Fig S22c) Alignment of Chromosome 4 *P. x bretschneideri* to *M. domestica* (bottom left)

(Fig S22d) Alignment of Chromosome 4 *P. communis* of Bartlettv1.0 to of BartlettDHv2.0

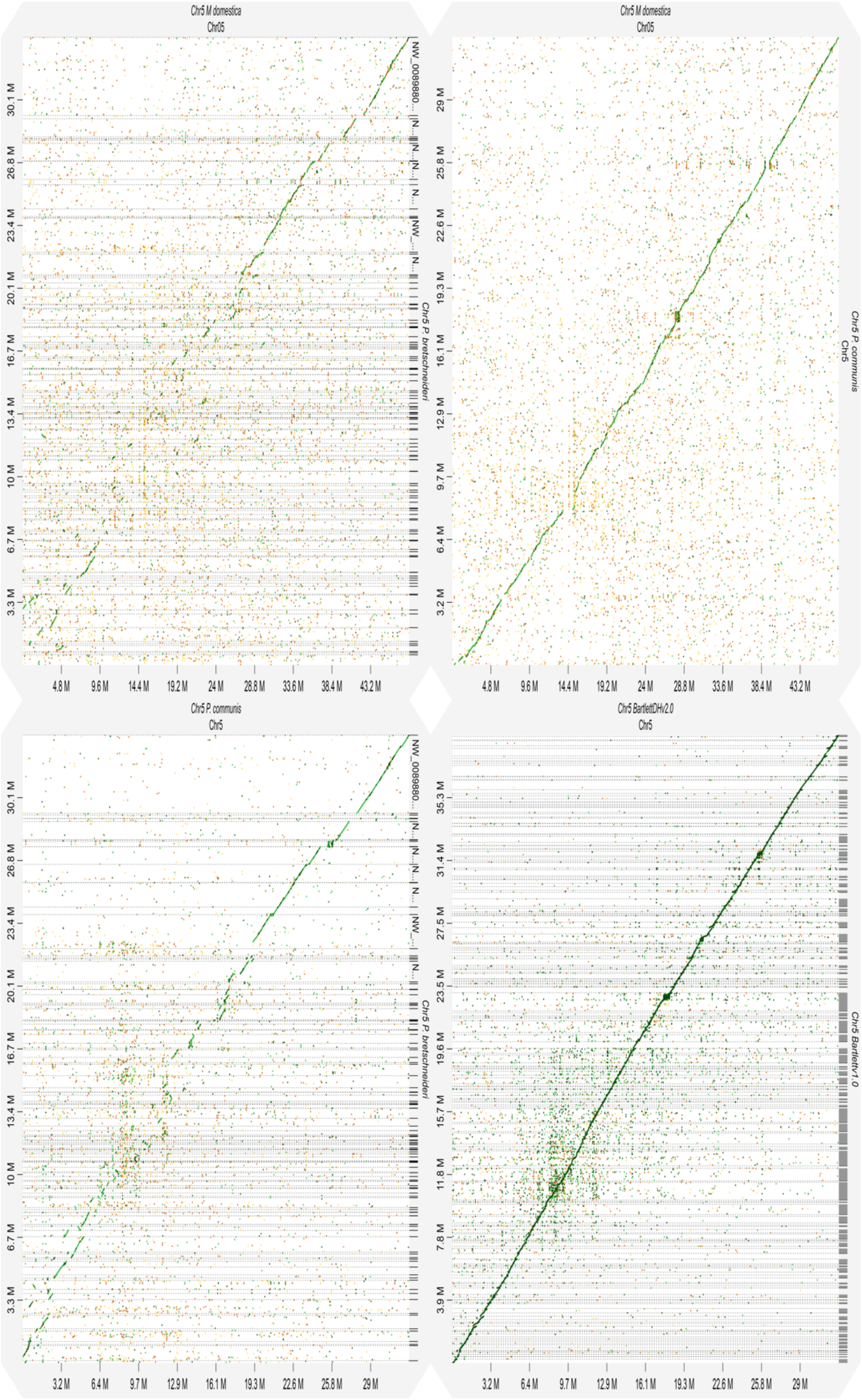


**Figure S23. Chromosome 5 alignment alignment dot plots. Dot plots are produced using the DGENIE software[60]and alignments with minimap2 (v2.16).**

(Fig S23a) Alignment of Chromosome 5 *P. x bretschneideri* to *P. communis* (top left)

(Fig S23b) Alignment of Chromosome 5 *P. communis* to *M. domestica* (top right)

(Fig S23c) Alignment of Chromosome 5 *P. x bretschneideri* to *M. domestica* (bottom left)

(Fig S23d) Alignment of Chromosome 5 *P. communis* of Bartlettv1.0 to of BartlettDHv2.0

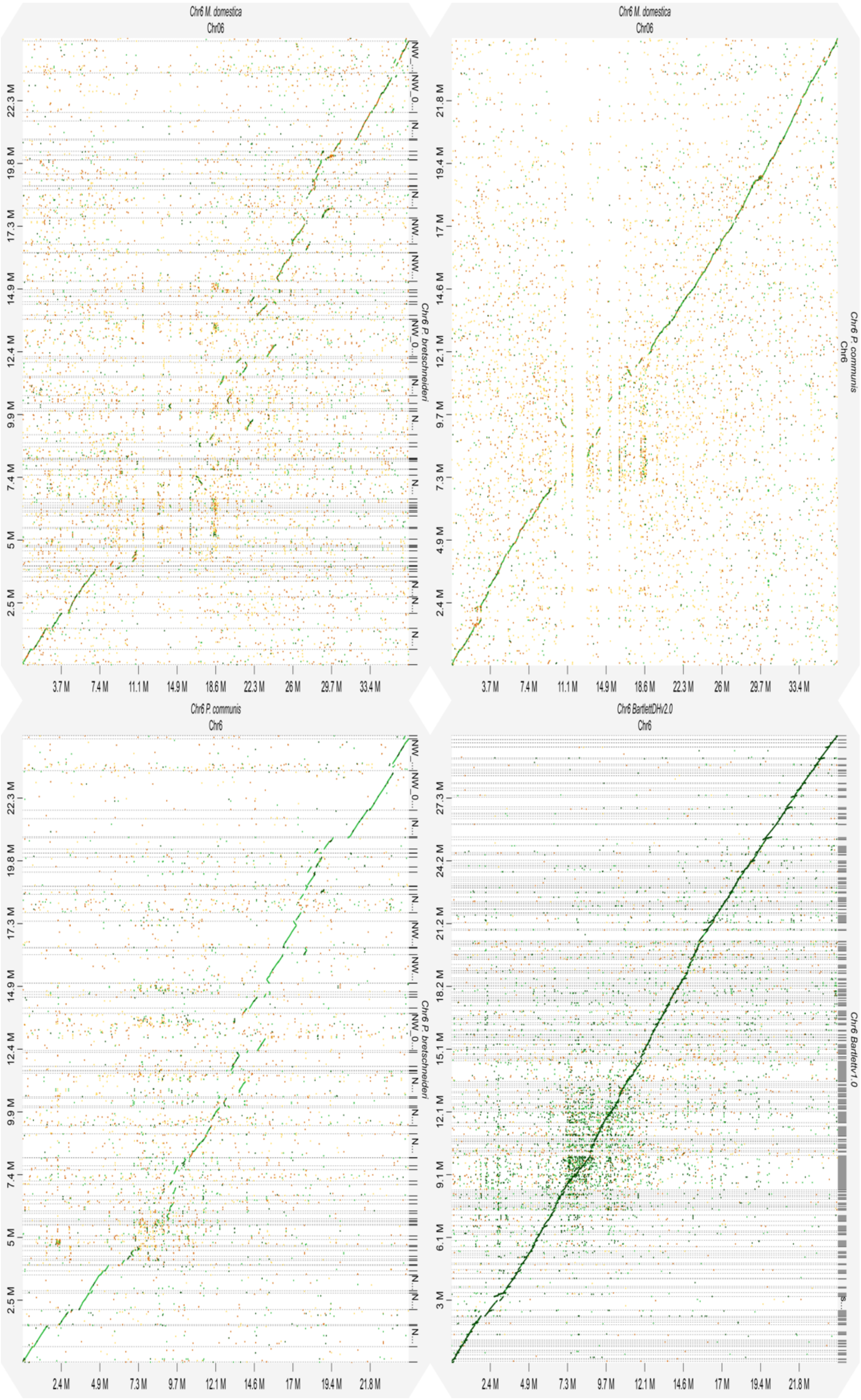

**Figure S24. Chromosome 6 alignment alignment dot plots. Dot plots are produced using the DGENIE software[60]and alignments with minimap2 (v2.16).**

(Fig S24a) Alignment of Chromosome 6 *P. x bretschneideri* to *P. communis* (top left)

(Fig S24b) Alignment of Chromosome 6 *P. communis* to *M. domestica* (top right)

(Fig S24c) Alignment of Chromosome 6 *P. x bretschneideri* to *M. domestica* (bottom left)

(Fig S24d) Alignment of Chromosome 6 *P. communis* of Bartlettv1.0 to of BartlettDHv2.0

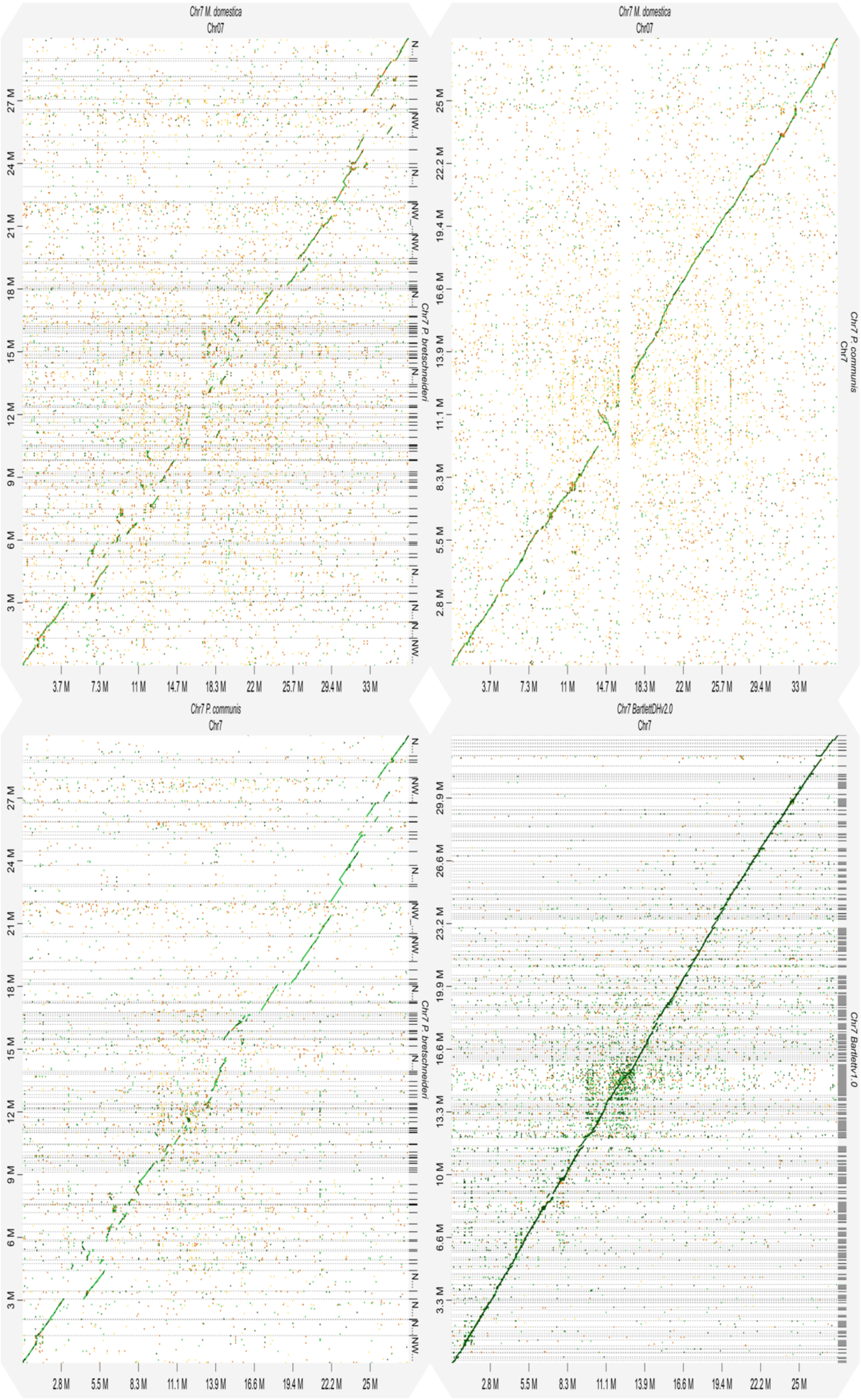

**Figure S25. Chromosome 7 alignment alignment dot plots. Dot plots are produced using the DGENIE software[60]and alignments with minimap2 (v2.16).**

(Fig S25a) Alignment of Chromosome 7 *P. x bretschneideri* to *P. communis* (top left)

(Fig S25b) Alignment of Chromosome 7 *P. communis* to *M. domestica* (top right)

(Fig S25c) Alignment of Chromosome 7 *P. x bretschneideri* to *M. domestica* (bottom left)

(Fig S25d) Alignment of Chromosome 7 *P. communis* of Bartlettv1.0 to of BartlettDHv2.0

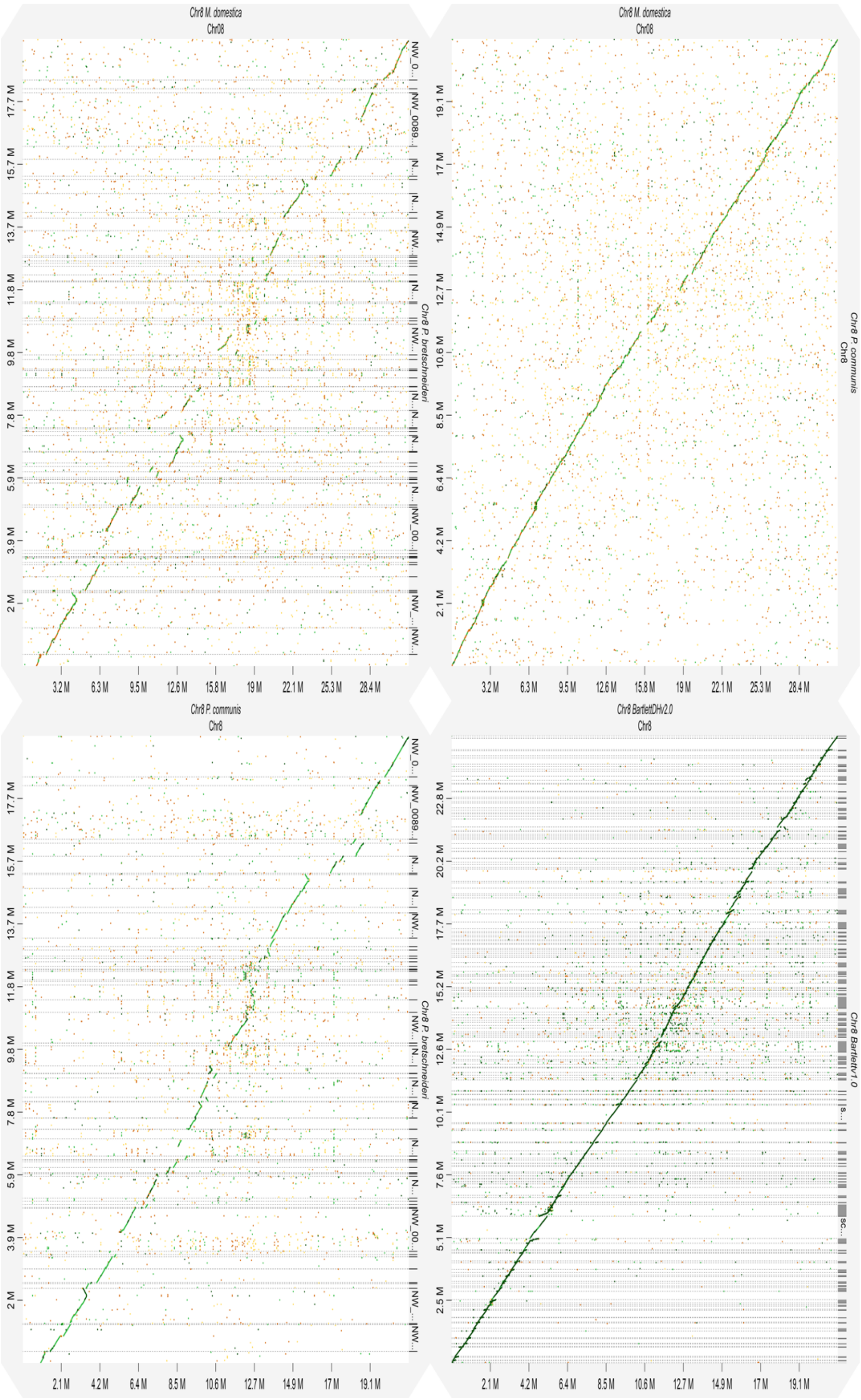


**Figure S26. Chromosome 8 alignment alignment dot plots. Dot plots are produced using the DGENIE software[60]and alignments with minimap2 (v2.16).**

(Fig S26a) Alignment of Chromosome 8 *P. x bretschneideri* to *P. communis* (top left)

(Fig S26b) Alignment of Chromosome 8 *P. communis* to *M. domestica* (top right)

(Fig S26c) Alignment of Chromosome 8 *P. x bretschneideri* to *M. domestica* (bottom left)

(Fig S26d) Alignment of Chromosome 8 *P. communis* of Bartlettv1.0 to of BartlettDHv2.0

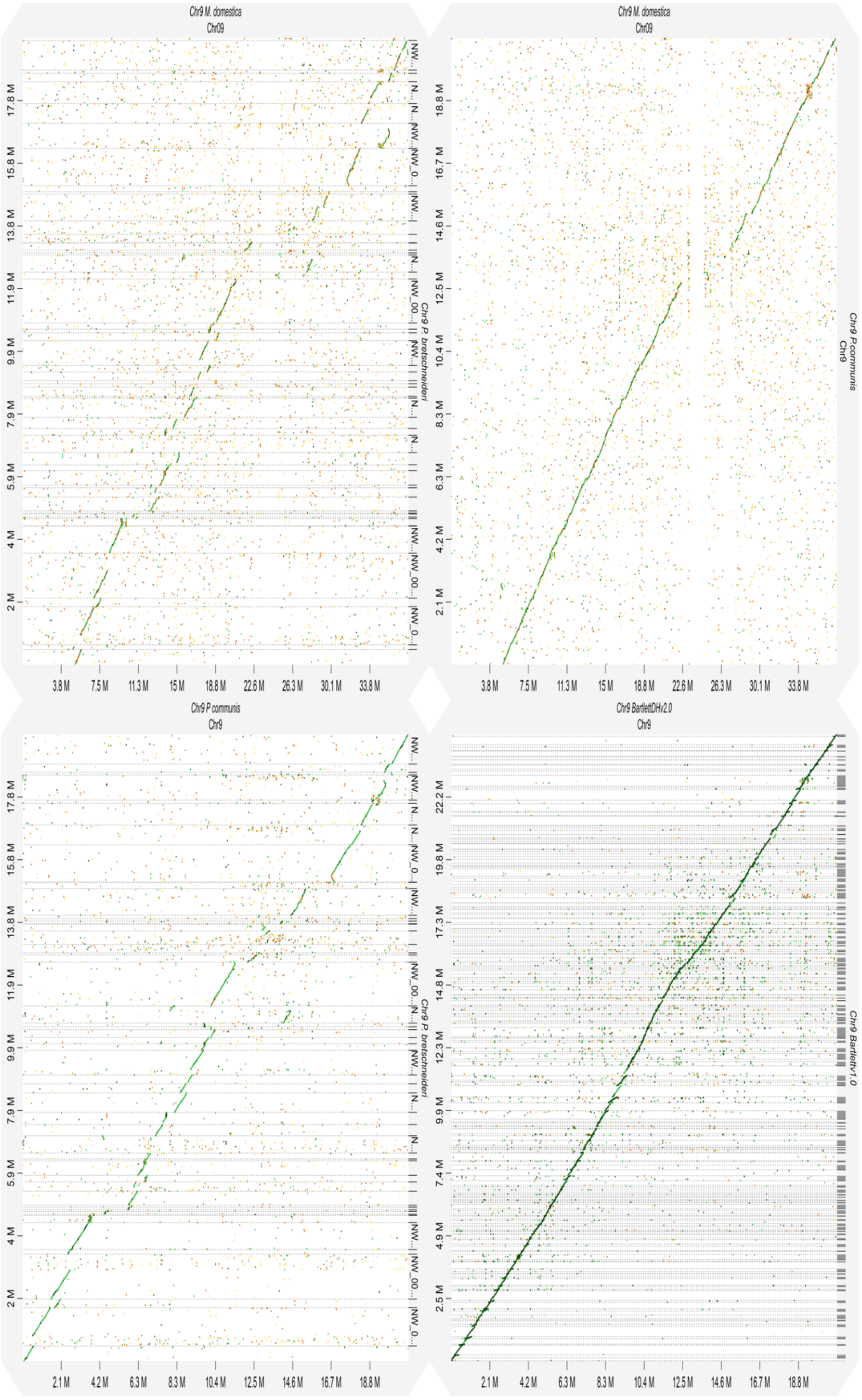


**Figure S27. Chromosome 9 alignment alignment dot plots. Dot plots are produced using the DGENIE software[60]and alignments with minimap2 (v2.16).**

(Fig S27a) Alignment of Chromosome 9 *P. x bretschneideri* to *P. communis* (top left)

(Fig S27b) Alignment of Chromosome 9 *P. communis* to *M. domestica* (top right)

(Fig S27c) Alignment of Chromosome 9 *P. x bretschneideri* to *M. domestica* (bottom left)

(Fig S27d) Alignment of Chromosome 9 *P. communis* of Bartlettv1.0 to of BartlettDHv2.0

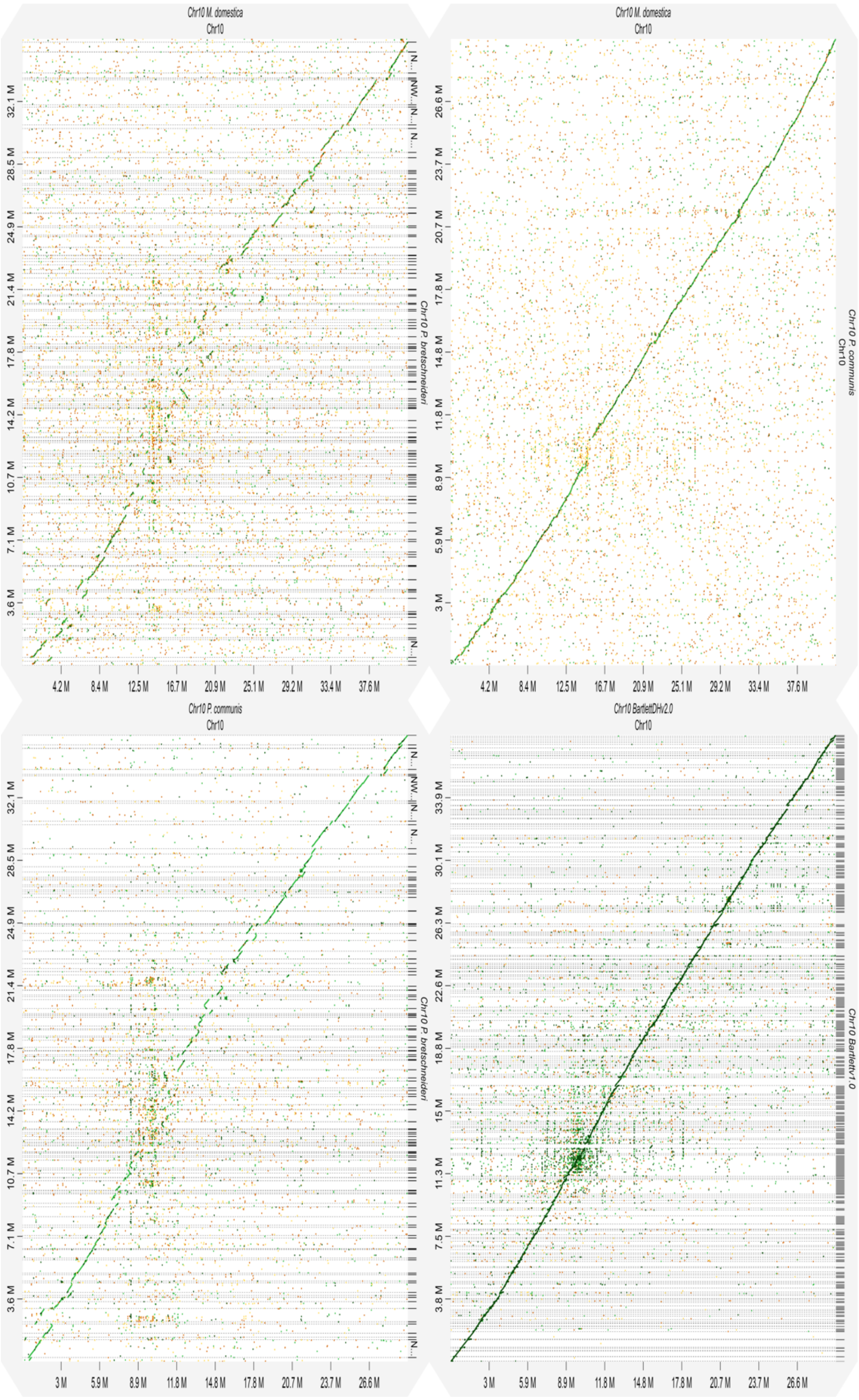


**Figure S28. Chromosome 10 alignment alignment dot plots. Dot plots are produced using the DGENIE software[60] and alignments with minimap2 (v2.16).**

(Fig S28a) Alignment of Chromosome 10 *P. x bretschneideri* to *P. communis* (top left)

(Fig S28b) Alignment of Chromosome 10 *P. communis* to *M. domestica* (top right)

(Fig S28c) Alignment of Chromosome 10 *P. x bretschneideri* to *M. domestica* (bottom left)

(Fig S28d) Alignment of Chromosome 10 *P. communis* of Bartlettv1.0 to of BartlettDHv2.0

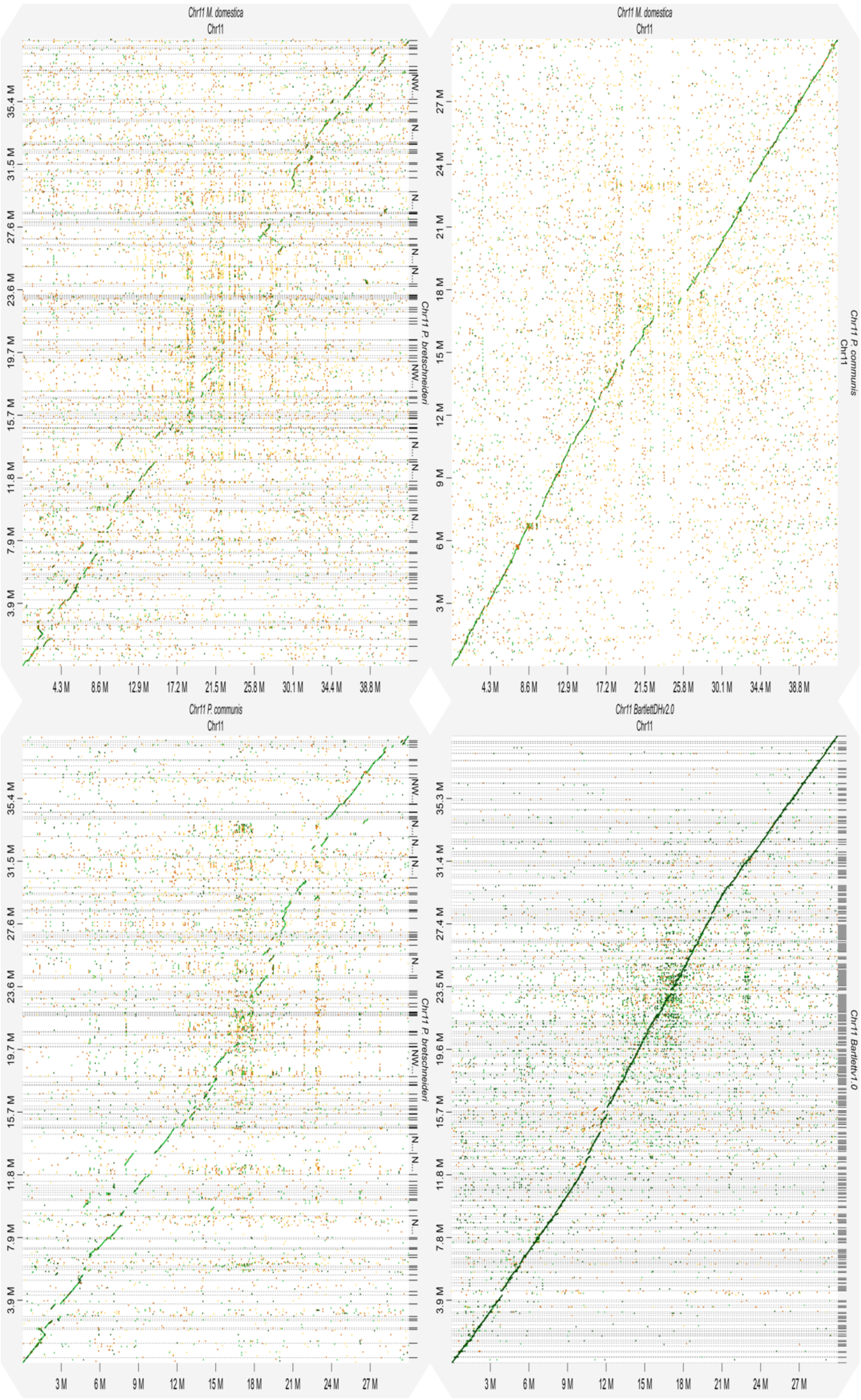


**Figure S29. Chromosome 11 alignment alignment dot plots. Dot plots are produced using the DGENIE software[60] and alignments with minimap2 (v2.16).**

(Fig S29a) Alignment of Chromosome 11 *P. x bretschneideri* to *P. communis* (top left)

(Fig S29b) Alignment of Chromosome 11 *P. communis* to *M. domestica* (top right)

(Fig S29c) Alignment of Chromosome 11 *P. x bretschneideri* to *M. domestica* (bottom left)

(Fig S29d) Alignment of Chromosome 11 *P. communis* of Bartlettv1.0 to of BartlettDHv2.0

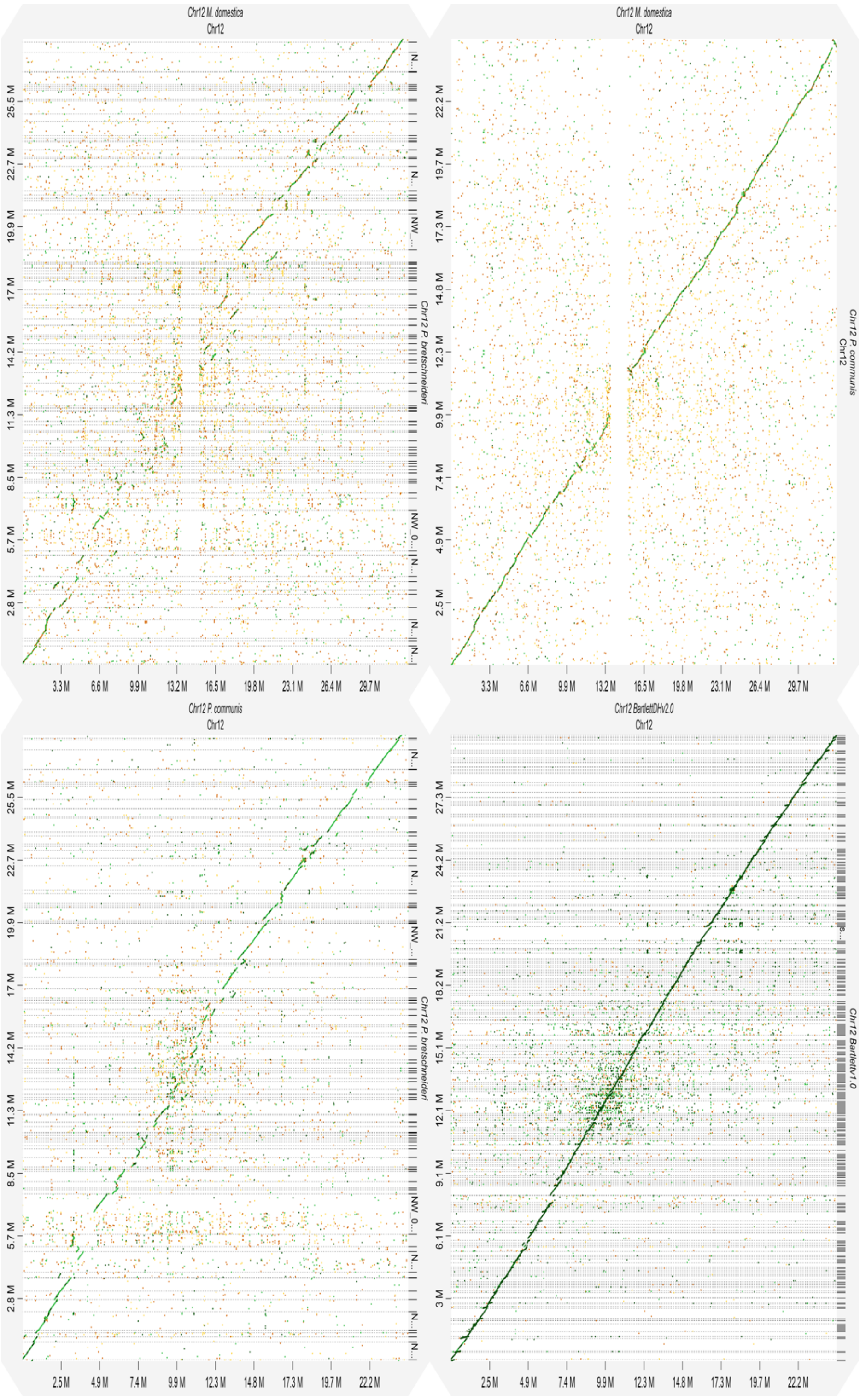


**Figure S230. Chromosome 12 alignment alignment dot plots. Dot plots are produced using the DGENIE software[60]and alignments with minimap2 (v2.16).**

(Fig S30a) Alignment of Chromosome 12 *P. x bretschneideri* to P. communis (top left)

(Fig S30b) Alignment of Chromosome 12 *P. communis* to *M. domestica* (top right)

(Fig S30c) Alignment of Chromosome 12 *P. x bretschneideri* to *M. domestica* (bottom left)

(Fig S30d) Alignment of Chromosome 12 *P. communis* of Bartlettv1.0 to of BartlettDHv2.0

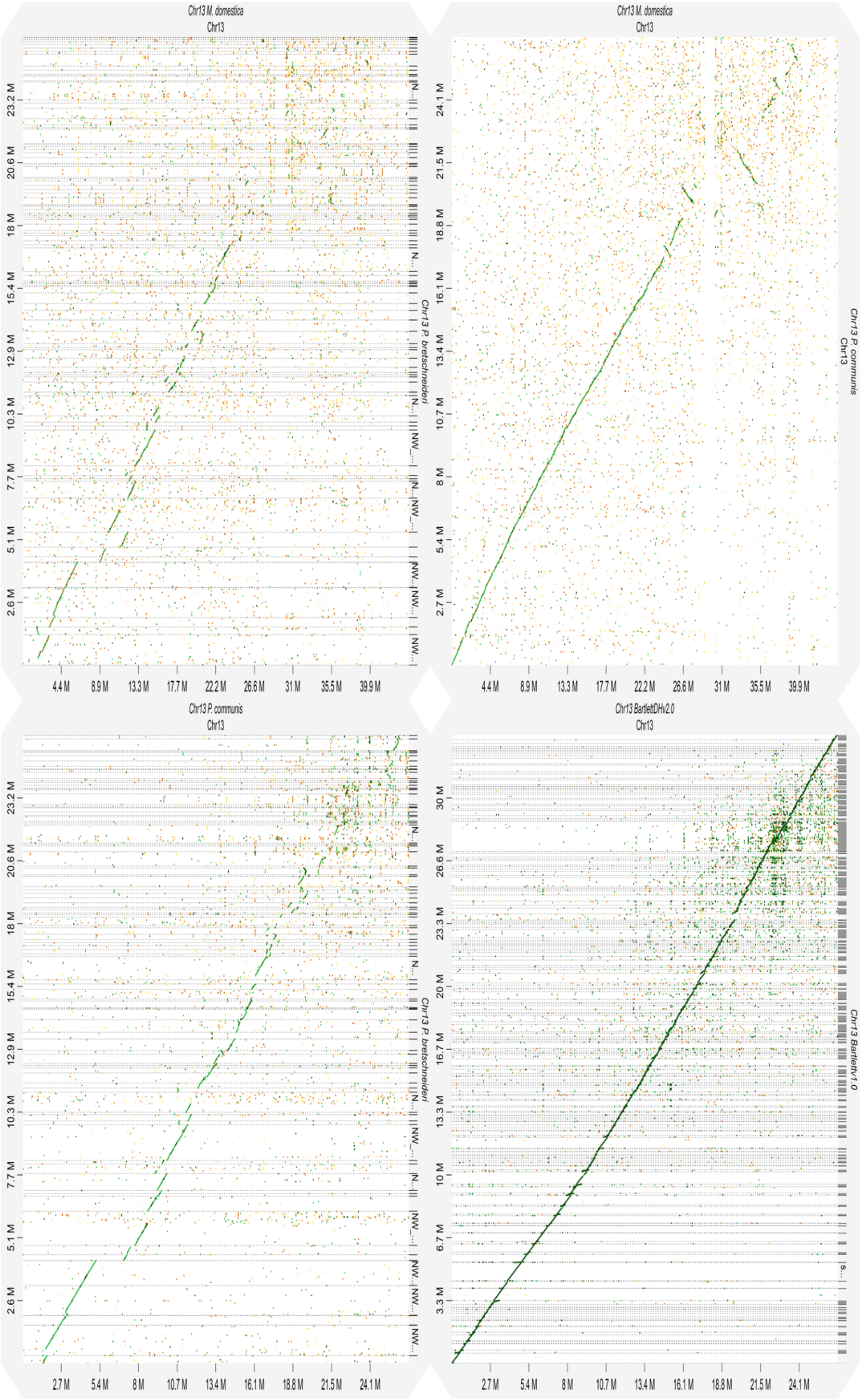


**Figure S31. Chromosome 13 alignment alignment dot plots. Dot plots are produced using the DGENIE software[60]and alignments with minimap2 (v2.16).**

(Fig S31a) Alignment of Chromosome 13 *P. x bretschneideri* to *P. communis* (top left)

(Fig S31b) Alignment of Chromosome 13 *P. communis* to *M. domestica* (top right)

(Fig S31c) Alignment of Chromosome 13 *P. x bretschneideri* to *M. domestica* (bottom left)

(Fig S31d) Alignment of Chromosome 13 *P. communis* of Bartlettv1.0 to of BartlettDHv2.0

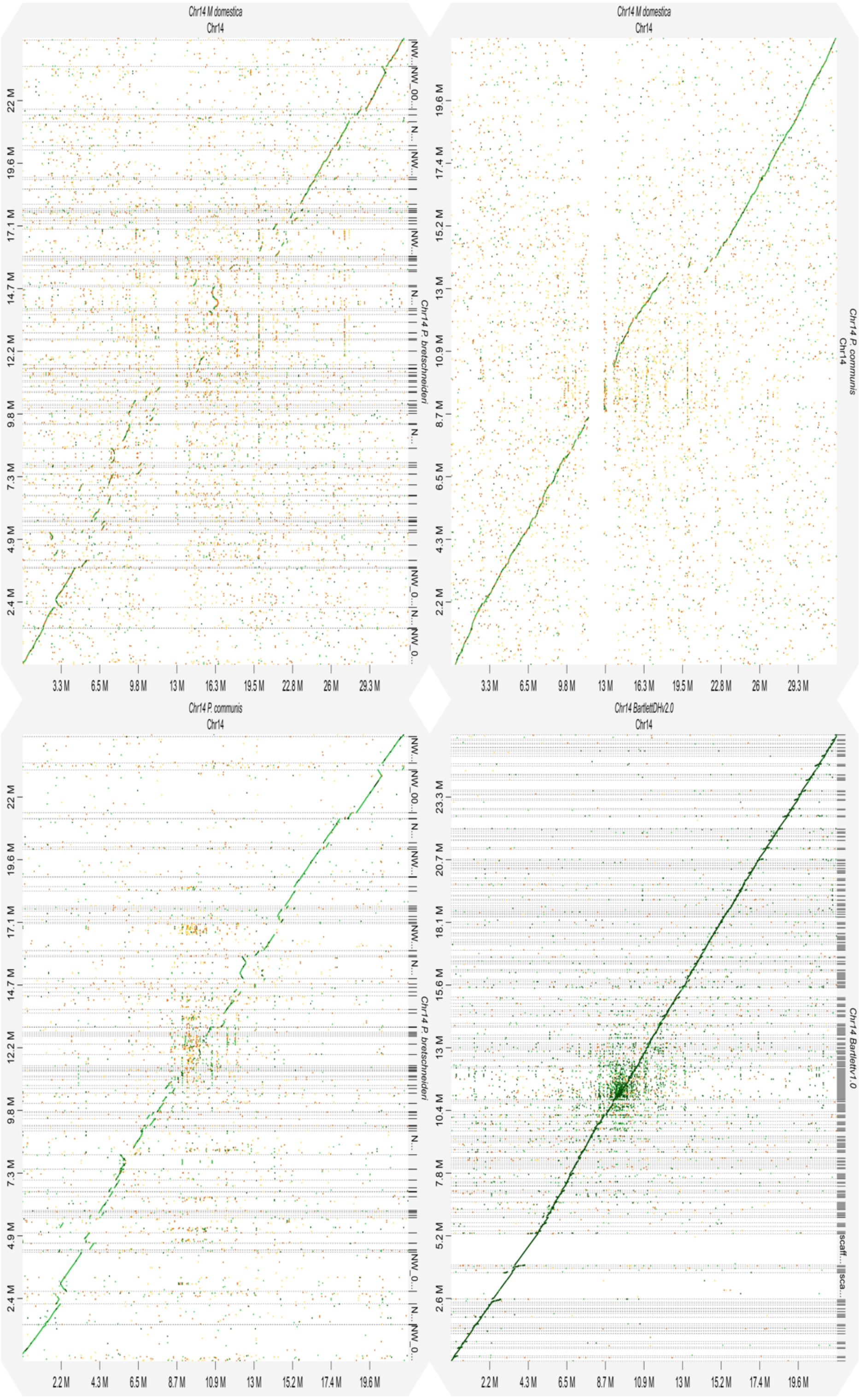


**Figure S32. Chromosome 14 alignment alignment dot plots. Dot plots are produced using the DGENIE software[60] and alignments with minimap2 (v2.16).**

(Fig S32a) Alignment of Chromosome 14 *P. x bretschneideri* to *P. communis* (top left)

(Fig S32b) Alignment of Chromosome 14 *P. communis* to *M. domestica* (top right)

(Fig S32c) Alignment of Chromosome 14 *P. x bretschneideri* to *M. domestica* (bottom left)

(Fig S32d) Alignment of Chromosome 14 *P. communis* of Bartlettv1.0 to of BartlettDHv2.0

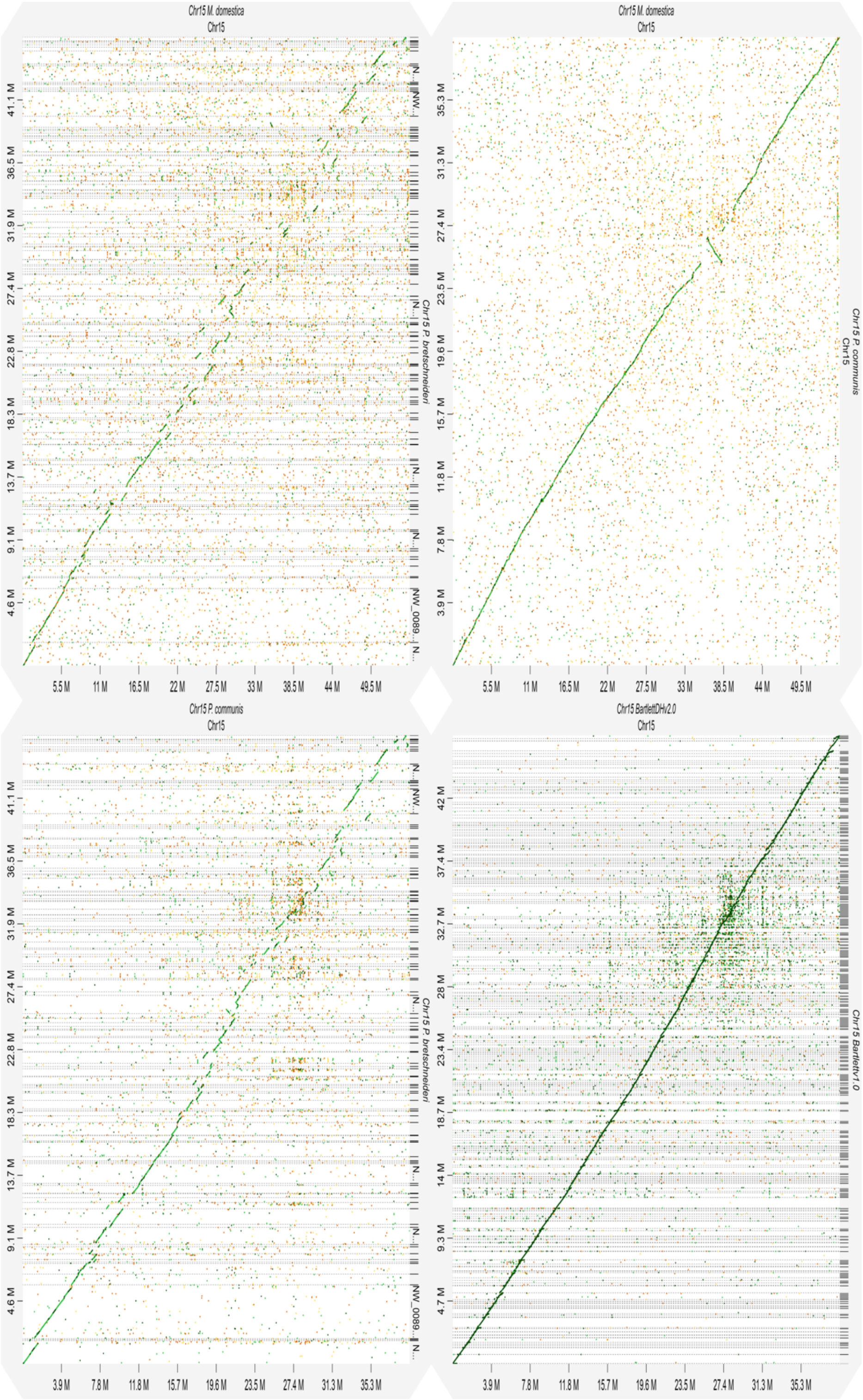


**Figure S33. Chromosome 15 alignment alignment dot plots. Dot plots are produced using the DGENIE software[60]and alignments with minimap2 (v2.16).**

(Fig S33a) Alignment of Chromosome 15 *P. x bretschneideri* to *P. communis* (top left)

(Fig S33b) Alignment of Chromosome 15 *P. communis* to *M. domestica* (top right)

(Fig S33c) Alignment of Chromosome 15 *P. x bretschneideri* to *M. domestica* (bottom left)

(Fig S33d) Alignment of Chromosome 15 *P. communis* of Bartlettv1.0 to of BartlettDHv2.0

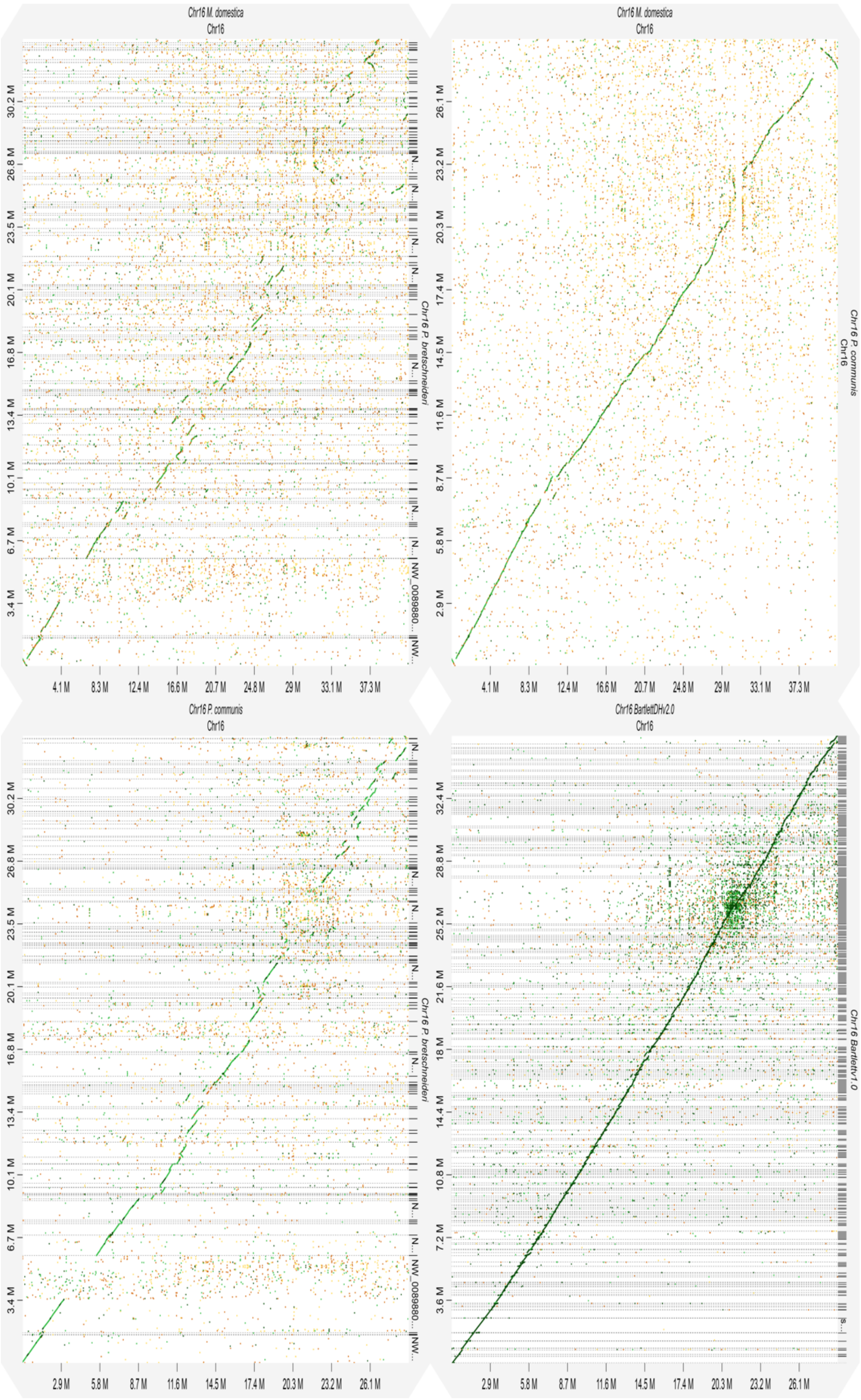


**Figure S34. Chromosome 16 alignment alignment dot plots. Dot plots are produced using the DGENIE software[60]and alignments with minimap2 (v2.16).**

(Fig S34a) Alignment of Chromosome 16 *P. x bretschneideri* to *P. communis* (top left)

(Fig S34b) Alignment of Chromosome 16 *P. communis* to *M. domestica* (top right)

(Fig S34c) Alignment of Chromosome 16 *P. x bretschneideri* to *M. domestica* (bottom left)

(Fig S34d) Alignment of Chromosome 16 *P. communis* of Bartlettv1.0 to of BartlettDHv2.0

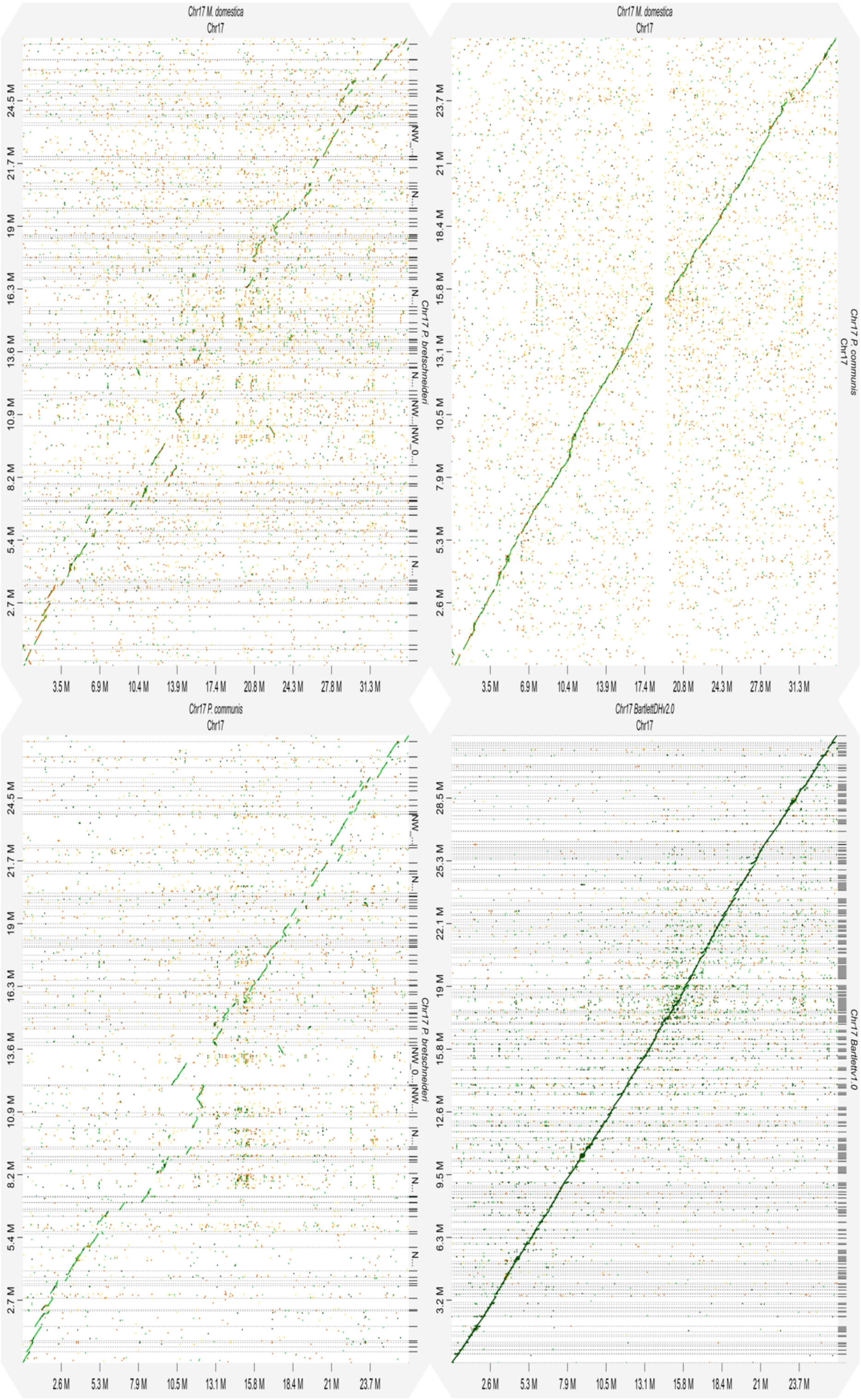

**Figure S35. Chromosome 17 alignment alignment dot plots. Dot plots are produced using the DGENIE software[60]and alignments with minimap2 (v2.16).**

(Fig S35a) Alignment of Chromosome 17 *P. x bretschneideri* to *P. communis* (top left)

(Fig S35b) Alignment of Chromosome 17 *P. communis* to *M. domestica* (top right)

(Fig S35c) Alignment of Chromosome 17 *P. x bretschneideri* to *M. domestica* (bottom left)

(Fig S35d) Alignment of Chromosome 17 *P. communis* of Bartlettv1.0 to of BartlettDHv2.0

**Enriched GO terms**

**Table S1 GO terms enriched in P. communis genes determined to be pomme specific**

414 pomme specific gene families containing 1,447 P. communis genes were enriched for the following GO terms.

| #Type | GO-id | Enrichment | p-value | subset-ratio | description |
|---|---|---|---|---|---|
| MF | GO:0001882 | 0.411353449 | 6.16E-04 | 25.03037667 | nucleoside binding |
| MF | GO:0001883 | 0.409747787 | 7.06E-04 | 24.90886999 | purine nucleoside binding |
| MF | GO:0003824 | 0.259594027 | 9.57E-07 | 59.29526124 | catalytic activity |
| MF | GO:0004872 | 0.98725488 | 0.038161026 | 4.009720535 | receptor activity |
| MF | GO:0004888 | 0.98725488 | 0.038161026 | 4.009720535 | transmembrane receptor activity |
| MF | GO:0005215 | 0.595920778 | 0.043424568 | 8.991494532 | transporter activity |
| MF | GO:0005216 | 1.580096674 | 0.010753402 | 2.187120292 | ion channel activity |
| MF | GO:0005524 | 0.462925951 | 7.16E-05 | 24.54434994 | ATP binding |
| MF | GO:0015075 | 1.066765598 | 2.41E-04 | 5.953827461 | ion transmembrane transporter activity |
| MF | GO:0015082 | 1.81322548 | 0.026644798 | 1.579586877 | di-, tri-valent inorganic cation transmembrane transporter activity |

| | | | | | |
|---|---|---|---|---|---|
| **MF** | **GO:0015267** | **1.562818683** | **0.012551231** | **2.187120292** | **channel activity** |
| **MF** | **GO:0015662** | **2.039783931** | **2.60E-04** | **2.065613609** | **ATPase activity, coupled to transmembrane movement of ions, phosphorylative mechanism** |
| **MF** | **GO:0017076** | **0.362399652** | **0.003413612** | **26.00243013** | **purine nucleotide binding** |
| **MF** | **GO:0022803** | **1.562818683** | **0.012551231** | **2.187120292** | **passive transmembrane transporter activity** |
| **MF** | **GO:0022804** | **0.929996543** | **0.041293348** | **4.374240583** | **active transmembrane transporter activity** |
| **MF** | **GO:0022838** | **1.580096674** | **0.010753402** | **2.187120292** | **substrate-specific channel activity** |
| **MF** | **GO:0022857** | **0.79510714** | **0.002745981** | **7.776427704** | **transmembrane transporter activity** |
| **MF** | **GO:0022891** | **0.958105226** | **3.31E-04** | **6.925880923** | **substrate-specific transmembrane transporter activity** |
| **MF** | **GO:0022892** | **0.883137525** | **8.73E-04** | **7.290400972** | **substrate-specific transporter activity** |

| | | | | | |
|---|---|---|---|---|---|
| **MF** | **GO:0030554** | **0.409747787** | **7.06E-04** | **24.90886999** | **adenyl nucleotide binding** |
| **MF** | **GO:0032553** | **0.414544354** | **3.86E-04** | **25.63791009** | **ribonucleotide binding** |
| **MF** | **GO:0032555** | **0.414544354** | **3.86E-04** | **25.63791009** | **purine ribonucleotide binding** |
| **MF** | **GO:0032559** | **0.462212099** | **7.42E-05** | **24.54434994** | **adenyl ribonucleotide binding** |
| **MF** | **GO:0042625** | **1.59654954** | **0.015306675** | **2.065613609** | **ATPase activity, coupled to transmembrane movement of ions** |
| **MF** | **GO:0046873** | **1.463283009** | **0.019407649** | **2.308626974** | **metal ion transmembrane transporter activity** |
| **BP** | **GO:0006754** | **1.515121941** | **0.044409183** | **2.065613609** | **ATP biosynthetic process** |
| **BP** | **GO:0009141** | **1.495704487** | **0.033186541** | **2.187120292** | **nucleoside triphosphate metabolic process** |
| **BP** | **GO:0009142** | **1.495704487** | **0.033186541** | **2.187120292** | **nucleoside triphosphate biosynthetic process** |

| | | | | | |
|---|---|---|---|---|---|
| BP | GO:0009144 | 1.495704487 | 0.033186541 | 2.187120292 | purine nucleoside triphosphate metabolic process |
| BP | GO:0009145 | 1.495704487 | 0.033186541 | 2.187120292 | purine nucleoside triphosphate biosynthetic process |
| BP | GO:0009199 | 1.495704487 | 0.033186541 | 2.187120292 | ribonucleoside triphosphate metabolic process |
| BP | GO:0009201 | 1.495704487 | 0.033186541 | 2.187120292 | ribonucleoside triphosphate biosynthetic process |
| BP | GO:0009205 | 1.495704487 | 0.033186541 | 2.187120292 | purine ribonucleoside triphosphate metabolic process |
| BP | GO:0009206 | 1.495704487 | 0.033186541 | 2.187120292 | purine ribonucleoside triphosphate biosynthetic process |
| BP | GO:0045087 | 1.037018255 | 0.038395427 | 3.888213852 | innate immune response |
| BP | GO:0046034 | 1.515121941 | 0.044409183 | 2.065613609 | ATP metabolic process |
| CC | GO:0005741 | 2.548171907 | 0.018169086 | 0.85054678 | mitochondrial outer membrane |
